# Using adopted individuals to partition maternal genetic effects into prenatal and postnatal effects on offspring phenotypes

**DOI:** 10.1101/2021.08.04.455178

**Authors:** Liang-Dar Hwang, Gunn-Helen Moen, David M. Evans

## Abstract

Maternal genetic effects can be defined as the effect of a mother’s genotype on the phenotype of her offspring, independent of the offspring’s genotype. Maternal genetic effects can act via the intrauterine environment during pregnancy and/or via the postnatal environment. In this manuscript, we present a simple extension to the basic adoption design that uses structural equation modelling (SEM) to partition maternal genetic effects into prenatal and postnatal effects. We assume that in biological families, offspring phenotypes are influenced prenatally by their mother’s genotype and postnatally by both parents’ genotypes, whereas adopted individuals’ phenotypes are influenced prenatally by their biological mother’s genotype and postnatally by their adoptive parents’ genotypes. Our SEM framework allows us to model the (potentially) unobserved genotypes of biological and adoptive parents as latent variables, permitting us in principle to leverage the thousands of adopted singleton individuals in the UK Biobank. We examine the power, utility and type I error rate of our model using simulations and asymptotic power calculations. We apply our model to polygenic scores of educational attainment and birth weight associated variants, in up to 5178 adopted singletons, 983 trios, 3650 mother-offspring pairs, 1665 father-offspring pairs and 350330 singletons from the UK Biobank. Our results show the expected pattern of maternal genetic effects on offspring birth weight, but unexpectedly large prenatal maternal genetic effects on offspring educational attainment. Sensitivity and simulation analyses suggest this result may be at least partially due to adopted individuals in the UK Biobank being raised by their biological relatives. We show that accurate modelling of these sorts of cryptic relationships is sufficient to bring type I error rate under control and produce unbiased estimates of prenatal and postnatal maternal genetic effects. We conclude that there would be considerable value in following up adopted individuals in the UK Biobank to determine whether they were raised by their biological relatives, and if so, to precisely ascertain the nature of these relationships. These adopted individuals could then be incorporated into informative statistical genetics models like the one described in our manuscript to further elucidate the genetic architecture of complex traits and diseases.

## Introduction

Maternal genetic effects can be defined as the causal influence of maternal genotypes on offspring phenotypes over and above that which results from the transmission of genes from mothers to their offspring (Wolf & Wade, 2009). Over the last few years there has been a resurgence of interest in identifying and quantifying maternal genetic effects on offspring phenotypes, both from the perspective of variance component estimation (Balbona et al., 2021; Bates et al., 2018; Eaves et al., 2014; Eilertsen et al., 2021; Kim et al., 2021; Kong et al., 2018; Qiao et al., 2020; Tubbs et al., 2020) and estimating the causal effect of specific maternal environmental exposures on offspring outcomes through Mendelian randomization based approaches (Evans et al., 2019; Lawlor et al., 2017; Moen et al., 2020; Tyrrell et al., 2016; Warrington et al., 2019; Zhang et al., 2015). Indeed, following the advent of transgenerational genome-wide association studies, maternal genetic effects are beginning to be identified at individual genetic loci (Beaumont et al., 2018; Warrington et al., 2019), a trend that is set to continue as the sample sizes of such studies increase further. Given the increasing number of variants identified, a natural question to ask is whether these loci exert their effects on offspring phenotype through intrauterine mechanisms, the postnatal environment, or both. Indeed, resolving maternal effects into prenatal and postnatal sources of variation could yield valuable insight into the underlying mechanisms behind these associations, and in the case of disease-related phenotypes, potentially important information regarding the optimal timing of interventions.

In this manuscript, we develop a simple method for partitioning maternal genetic effects into prenatal and postnatal components that leverages information provided by adopted individuals. We assume that an adopted individual’s phenotype is influenced by prenatal intrauterine factors as proxied by their biological mother’s genome, and postnatal influences as proxied by their adoptive mother’s and father’s genomes. In contrast, we assume that the phenotype of individuals who have not been adopted are influenced by prenatal intrauterine factors resulting from their biological mother’s genome, and postnatal factors as proxied by their biological mother’s and father’s genomes. This model leads to different expectations for the covariance between an individual’s phenotype and their own and their relatives’ genotypes depending on whether they have been adopted or not.

One of the challenges in applying this kind of framework to real life situations is the paucity of cohorts containing large numbers of adopted families (Horn, 1983; Rhea et al., 2013; Scarr & Weinberg, 1983). Restricted numbers of adopted families will consequently limit the statistical power to partition maternal genetic effects- particularly at single genetic variants which tend to have very small effect sizes. It is important to realize, however, that adopted “singletons” (i.e. adopted individuals whose adoptive and biological parents have not been included in the study) provide important information on partitioning maternal genetic effects into prenatal and postnatal contributions regardless of whether information on their parents has been gathered. The intuition behind this surprising fact is that the covariance between an adopted individual’s genotype and phenotype is a function of prenatal (but not postnatal) maternal genetic effects (Figure 1, G5 to G7). In contrast, the covariance between a non-adopted individual and their own phenotype includes contributions from prenatal and postnatal genetic effects (Figure 1, G1 to G4). Thus, the difference between the genotype-phenotype covariance in adopted and non-adopted singleton individuals provides important information on the likely size of postnatal genetic effects. This is potentially important given that >7000 individuals in the UK Biobank study (Sudlow et al., 2015) report being adopted, and hence this publicly available dataset may represent a powerful resource for identifying prenatal maternal effects and partitioning maternal genetic effects into prenatal and postnatal sources of variation.

In this manuscript, we introduce a new structural equation model (SEM) to estimate prenatal and postnatal maternal genetic effects on offspring phenotypes. The SEM framework allows us to model the contributions from missing individuals as latent variables, facilitating the inclusion of information from adopted singleton individuals whose adoptive and/or biological parents are not available for analysis. We estimate the asymptotic power of our model to partition maternal genetic effects into prenatal and postnatal components and confirm these results by simulation. We code our routines into a freely available R shiny app web utility that researchers can use to perform their own power calculations. Finally, we apply our methods to birth weight and educational attainment data in the UK Biobank- two phenotypes known to be affected by maternal genetic effects. We investigate the effect of inadvertently including adopted singleton individuals whose adoptive parents are genetically related to them in our model, adjust our model to correct for this misspecification and quantify the effect of this adjustment on type 1 error rates and power. We discuss the implications of model misspecification for using the UK Biobank resource in this context in future studies more broadly.

## Methods

### Model Description

We consider 7 different family structures (Figure 1) which we model using SEM (although we note that the SEM framework is flexible enough to include many other sorts of family structures as well). The different family structures we consider are: Biological parent-offspring trios (G1), biological mother-offspring pairs (G2), biological father-offspring pairs (G3), singleton individuals (who were raised by their biological parents) (G4), adopted singleton individuals (who were raised by their adoptive parents) (G5), adoptive mother-adopted child pairs (G6), and biological mother-adopted child pairs (G7). For each of these different family structures, we model missing parental genotypes (whether biological or adoptive) as latent variables.

Let the variables Z_m_, Z_p_ and Z_o_ represent maternal, paternal and offspring genotypes. Likewise, let Z_m_b_ and Z_p_b_ denote the genotypes of the biological mother and father of an adopted individual respectively, and let Z_m_a_ and Z_p_a_ represent the genotypes of the adoptive mother and father of the adopted individual. These variables may be observed (i.e. the square boxes in Figure 1) or unobserved latent variables (i.e. the circles in Figure 1). They may represent genetic risk scores consisting of multiple genetic variants, or single genetic variants if the study is of sufficient size to reliably demonstrate genetic effects at individual loci. The model includes path coefficient terms for maternal genetic effects (*γ_m_*, *β_m_*), paternal genetic effects (*β_p_*) and offspring genetic effects (*β_o_*) on the offspring phenotype (Y). We are specifically interested in partitioning maternal genetic effects into prenatal (γ_m_) and postnatal genetic effects (*β_m_*) and we assume that the effect of the prenatal maternal genetic effect and postnatal maternal genetic effect on the offspring phenotype is additive (i.e. *γ_m_* + *β_m_*). We assume in the case of biological families (i.e. G1 to G4 in Figure 1), that biological mothers exert prenatal and postnatal genetic effects on their offspring. In contrast, in the case of adopted individuals (i.e. G5 to G7 in Figure 1), we assume that prenatal maternal effects from the biological mother and postnatal maternal genetic effects from the adoptive mother contribute to variation in the offspring phenotype. We assume that biological fathers exert only postnatal genetic effects on their offspring (biological families only), and adoptive fathers exert only postnatal genetic effects on their adoptive offspring.

We estimate the genotypic variance (*∅*) of each individual and assume that it is the same in mothers, fathers, their children, and across both biological and adopted individuals under random mating (although the requirement for equal variances across individuals can be relaxed if necessary). We account for possible covariance between maternal and paternal genotypes (e.g. through assortative mating) by estimating the covariance path *ρ*. This is equivalent to modelling one round of assortative mating in the parental generation and has the effect of inflating the total genotypic variance in the offspring generation from *∅* under random mating to *∅* + ½*ρ*. Finally, to accommodate the possibility that adopted individuals may have different phenotypic variances to individuals raised by their biological relatives, we permit the variance of residual sources of variation to differ across adopted and biological family structures (Cheesman et al., 2020).

We caution that for all the parameters in the model to be identified, different constellations of relatives must be sampled from G1-G7. For example, to partition maternal genetic effects into prenatal and postnatal components, information from adopted individuals must be available. In fact, even adopted “singletons”, for whom there is no genotype information from their parents (i.e. biological or adoptive parents), contribute important information for the partitioning of maternal genetic effects, since the covariance between their own genetic score and phenotype is a function of offspring genetic effects and prenatal maternal effects, but not postnatal maternal effects (Figure 1). This contrasts with the situation in non-adopted individuals whose genotype-phenotype covariance is a function of all three sources of variation (plus postnatal paternal genetic effects). Thus, the difference between the genotype-phenotype covariance in adopted and non-adopted individuals provides important information on the size of postnatal maternal genetic effects. Indeed, the inclusion of adopted singleton individuals in our model is sufficient for identification as long as there are at least some (biological) parent-offspring trios (or alternatively both biological mother-child and biological father-child pairs) available. Whilst the presence of adopted individuals’ parents (biological or adoptive) is not a necessary condition for partitioning maternal effects into prenatal and postnatal components, the inclusion of such pairs is advantageous in terms of statistical power as we show below. We include an example R script that fits the sort of SEM described in this article that users can modify to apply to their own data (see Supplementary Materials).

### Asymptotic Power Calculations

We used OpenMx (Boker et al., 2011; Neale et al., 2016) to calculate the asymptotic power to partition maternal genetic effects into prenatal and postnatal maternal genetic effects. Asymptotic covariance matrices were generated assuming certain underlying values for the parameters of the model depicted in Figure 1. The full model was then fitted to the covariance matrices to confirm a perfect fit to the data and to check that the parameter values were correctly estimated. Secondly, a reduced model where the parameter(s) of interest was constrained to zero was fitted to the same covariance matrices. We examined the effect of constraining the prenatal maternal genetic effect or postnatal maternal genetic effect to zero. The difference in minus twice the log-likelihood chi-square between the full and reduced models is equal to the non-centrality parameter of the test for association, with the degrees of freedom equal to the difference in the number of free parameters between the models. Power was then calculated as the area under the curve of a non-central chi-square distribution to the right of the significance threshold of interest:

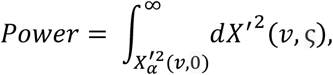

where 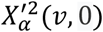 is the quantile of the 100 * (1-α) percentage point of the central *χ^2^* distribution with *v* degrees of freedom, and ς is the non-centrality parameter (Moen et al., 2019).

As we were interested in whether there might be enough adopted individuals in the UK Biobank for the informative partitioning of maternal genetic effects into prenatal and postnatal sources of variation, we first calculated power (α = 0.05) using sample sizes roughly similar to the number of European individuals in the resource who reported their educational attainment (i.e. 1000 parent-offspring trios, 4000 mother-offspring pairs, 1800 father-offspring pairs, 300000 singletons who were raised by their biological parents, 6000 adopted individuals and 50 biological mother-adopted offspring duos-see below). We investigated the effect of different combinations of prenatal maternal genetic effects, postnatal maternal genetic effects, postnatal paternal genetic effects and offspring genetic effects (i.e. *γ*_m_ or *β*_m_ = 0, 0.05, 0.1, 0.3 and 0.5). We also examined the effect of modifying the covariance between parental genotypes (with *ρ* ranging between 0 and 0.2 times the genotypic variance in the parental generation). Finally, we were curious as to the effect of increasing the number of the different family structures on statistical power, and in particular, on whether including increasing numbers of more attainable/accessible biological relatives (i.e. biological trios, pairs, and singletons [G1 – G4]) could increase power to partition maternal genetic effects for a fixed number of adopted individuals. We also investigated the effect of varying the sample size of adopted individuals (G5), adoptive mother-adopted offspring duos (G6), and biological mother-adopted offspring duos (G7) on power. We used our calculator to determine 80% power (α = 0.05).

### Data Simulations

In order to confirm our asymptotic calculations, we simulated a series of datasets of the same size and with the same underlying parameters as in the asymptotic power calculations. Each condition was simulated 1000 times and power was calculated as the number of simulations in which the effect of interest was detected (p<0.05) divided by 1000.

### Online Web Utility

We developed a freely available, online power calculator that allows investigators to explore the power to detect prenatal and postnatal maternal genetic effects using our SEM in their own studies (https://evansgroup.di.uq.edu.au/ADOPTED). Our power calculator is built using the R shiny app (https://shiny.rstudio.com/) running the R studio software (RStudioTeam, 2015) and the OpenMx package (Boker et al., 2011; Neale et al., 2016) in the background.

### Application to UK Biobank Data

We applied our model to self-reported birth weight and educational attainment data from the UK Biobank resource. We only included individuals of European ancestry, determined by: i) projecting their genetic principal components onto the 1000 Genome sample and then K-means clustering or ii) those who self-reported ethnic background of either “British”, “Irish”, “Irish”, “White”, or “Any other white background”. Family relationships were determined by the pairwise kinship estimated across the whole UK Biobank sample using the software KING (Manichaikul et al., 2010). Adoption status was determined based on response to the question “*Were you adopted as a child?*”.

Birth weight is a phenotype that is known to be affected by the maternal genome (Warrington et al., 2019), but should only be influenced by prenatal and not postnatal maternal genetic effects. We only included individuals whose birth weight was between 2.5kg and 4.5kg in analyses. These included 771 biological trios, 2776 biological mother-offspring pairs, 1233 biological father-offspring pairs, 172183 singletons raised by their biological parents, 1084 singleton individuals who reported being adopted, and 18 biological mother-adopted offspring pairs. We calculated each individual’s weighted polygenic risk score for birth weight using 20 SNPs known to influence birth weight via maternal pathways with P < 5 x 10^−8^ from a recent GWAS of birthweight (Warrington et al., 2019). See Supplementary Table 1 for a list of SNPs used to construct polygenic risk scores of birth weight.

We then applied our model to educational attainment in UK Biobank participants of European ancestry. We calculated years of education, as a proxy of educational attainment, based on response to the question “*Which of the following qualifications do you have?*”: College or university degree = 20 years; Vocational Qualification (NVQ), Higher National Diploma (HNC), or equivalent = 19 years; other professional qualification = 15 years; A level/AS level or equivalent = 13 years; O level/General Certificate of Secondary Education (GCSE) or equivalent = 10 years; Certificate of Secondary Education (CSE) or equivalent = 10 years; none of the above = 7 years of education. We regressed years of education on year of birth, sex and the first five genetic principal components and used the residuals in the analysis. The final dataset included 983 biological parent-offspring trios, 3650 biological mother-offspring pairs, 1665 biological father-offspring pairs, 350330 singletons from biological families, 5178 adopted singletons and 41 biological mother-adopted offspring pairs. We calculated unweighted polygenic risk scores (i.e. counts of the educational attainment increasing allele) using 1267 SNPs from the current largest GWAS of educational attainment (Lee et al., 2018). Because this GWAS included individuals from the UK Biobank, we performed a sensitivity analysis using a second set of polygenic risk scores of educational attainment using 72 SNPs from a GWAS of non-UK Biobank individuals (Okbay et al., 2016). See Supplementary Table 2 for a list of SNPs used to construct these polygenic risk scores.

The UK Biobank contains only very limited information regarding the adoption status of individuals (i.e. only whether they report being adopted or not). We were concerned about the possibility that some individuals in UK Biobank might have been adopted by their biological relatives in which case our model in Figure 1 would be misspecified. We therefore also conducted a sensitivity analysis by excluding adopted individuals with known breastfeeding information (on the reasoning that these individuals might be more likely to be fostered by a biologically related individual) and repeating the educational attainment analyses. This reduced the number of adopted “singleton” individuals in the analysis to 2867.

### Investigating the Effect of Model Misspecification

We were concerned about the effect that misclassifying adopted “singleton” individuals as unrelated to their adoptive parents (i.e. when in reality they were raised by their biological relatives) might have on type I error rates and estimates of prenatal and postnatal maternal genetic effects generated by our model. We therefore simulated data under four scenarios: i) adopted singletons where 0%, 20%, 40%, 60% or 80% had adoptive mothers who were siblings or cousins of their biological mothers, ii) adopted singletons where 0%, 20%, 40%, 60% or 80% had adoptive mother who were siblings or cousins of biological fathers, iii) adopted singletons where 0%, 20%, 40%, 60% or 80% had adoptive fathers who were siblings or cousins of biological fathers, and vi) adopted singletons where 0%, 20%, 40%, 60% or 80% had adoptive fathers who were siblings or cousins of biological mothers. We assumed that sample sizes and the number of individuals in the other groups were the same as in our asymptotic power calculations.

We further investigated type 1 error rates and estimates of prenatal and postnatal maternal genetic effects after correctly modelling the relationship between biological and adoptive parents. We created four additional family structures (G8 to G11, Supplementary Figure 1) considering the above four scenarios by adding a covariance path (*r*) between the genotypes of adoptive and biological parents. The covariance between the genotypes of adoptive and biological parents was fixed to *r* = 0.5 х *∅* when the adoptive parent was a sibling of the biological parent and *r* = 0.125 х *∅* when they were cousins.

## Results

### Power calculations

The power to detect prenatal maternal genetic effects increased with increasing size of postnatal maternal genetic effects, paternal genetic effects, and offspring genetic effects (Figure 2). A similar pattern was observed when we quantified the power to detect postnatal maternal genetic effects. Our calculations indicate that a study with a sample size and structure similar to the UK Biobank would have ~80% power (α=0.05) to detect prenatal maternal and postnatal maternal genetic effects that account for 1.5% and 0.9% of the variance in the offspring phenotype respectively (calculated as *γ*_m_^2^ and *β*_m_^2^ assuming all the other genetic effects are null and zero correlation between parental genotypes). The power to detect prenatal and postnatal maternal genetic effects also increased slightly with increasing covariance between maternal and paternal genotypes (Supplementary Figure 2).

The power to detect prenatal and postnatal maternal genetic effects increased with the number of adopted singleton individuals, and more dramatically when the number of adoptive mother-adopted offspring pairs and/or biological mother-adopted offspring pairs increased (Figure 3). Power to detect prenatal maternal genetic effects increased most rapidly with increasing numbers of biological-mother adopted child pairs (G7), whereas the power to detect postnatal maternal genetic effects increased most strongly with increasing numbers of was provided by adoptive mother-adopted child pairs (G6). This makes sense intuitively, since adoptive mother-adopted child pairs (G6) provide a direct estimate of the postnatal maternal genetic effect (*β*_m_), whereas biological mother-adopted child pairs provide a direct estimate of the prenatal maternal genetic effect (*γ*_m_). Likewise, adopted singleton individuals do not provide direct estimates of either prenatal or postnatal maternal genetic effects, and therefore contribute proportionally less in terms of statistical power than adoptive mother-adopted child or biological mother-adopted child pairs.

Interestingly, the power to detect prenatal and postnatal maternal genetic effects also increased with the number of biological families in the analysis (G1 to G4) (Figure 3), however, this increase asymptoted with maximal possible power depending primarily on the number of adopted individuals in the analysis. This result suggests that power to partition maternal genetic effects can be optimized through having large numbers of biological relatives in the analysis, but cannot be increased beyond a level that depends on the number of adopted individuals in the analysis (G5 to G7) and may be less than 100%. Results from simulations showed good concordance with asymptotic power calculations and appropriate type I error rates (Supplementary Tables 3 and 4).

### Analyses in the UK Biobank

Genetic risk scores for birth weight exhibited significant evidence of prenatal maternal genetic effects, offspring genetic effects (in the opposite direction), and no significant evidence for postnatal maternal or paternal genetic effects (Table 1). This pattern of results is expected since (a) birth weight is known to be affected by both maternal and fetal genetic effects, (b) maternal and offspring genetic effects on birth weight for many of the SNPs were in opposite directions, (c) conditional estimates of maternal and offspring genetic effects are negatively correlated, (d) in the present study SNPs comprising the genetic risk scores were selected for their strong maternal effects on offspring birth weight, and (e) birth weight is a perinatal phenotype and so by definition cannot be influenced by postnatal maternal (or paternal) genetic effects (Beaumont et al., 2018; Warrington et al., 2019; Warrington et al., 2018).

**Table 1.**
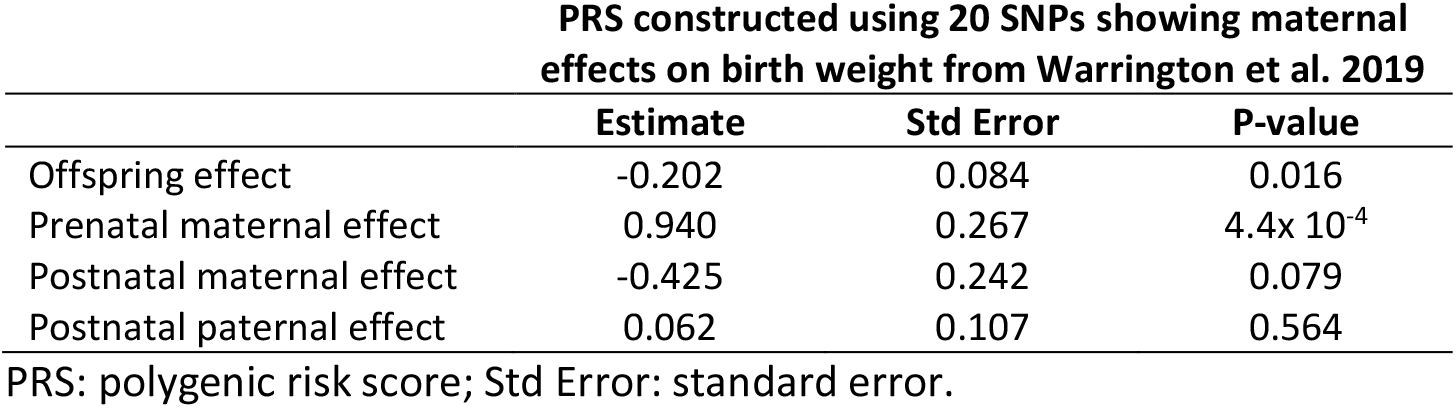
Modelling results of birth weight in the UK Biobank.

In the case of offspring educational attainment, our analyses showed evidence for a prenatal maternal genetic effect, a paternal genetic effect, and an offspring genetic effect (Table 2). The absence of a significant postnatal maternal genetic effect on offspring educational attainment was surprising given that maternal genetic effects should be mediated through maternal educational attainment, and therefore should mostly involve postnatal pathways (although it is possible that some of the relationship between maternal genotype and offspring educational attainment could be mediated through prenatal effects-e.g. less educated mothers consuming more alcohol during pregnancy which then has adverse effects on offspring cognitive development and educational attainment etc). We were concerned that the cause of this surprising result might be because of model misspecification-specifically, adopted children being raised by adoptive parents who are biologically related to them.

**Table 2.** Modelling results of educational attainment in the UK Biobank.

| PRS constructed using 1267 SNPs from Lee et al. 2018 |  |  |  |  |  |  |
| --- | --- | --- | --- | --- | --- | --- |
|  | Full sample |  |  | Excluding adopted individuals with breastfeeding information |  |  |
|  | Estimate | Std Error | P-value | Estimate | Std Error | P-value |
| Offspring effect | 0.026 | 0.004 | $4.41 \times 10^{-10}$ | 0.026 | 0.004 | $7.79 \times 10^{-10}$ |
| Prenatal maternal effect | 0.025 | 0.009 | 0.008 | 0.016 | 0.010 | 0.137 |
| Postnatal maternal effect | -0.004 | 0.007 | 0.567 | 0.006 | 0.009 | 0.518 |
| Postnatal paternal effect | 0.016 | 0.005 | 0.002 | 0.017 | 0.005 | 0.002 |

| PRS constructed using 72 SNPs from Okbay et al. 2016 |  |  |  |  |  |  |
| --- | --- | --- | --- | --- | --- | --- |
|  | Full sample |  |  | Excluding adopted individuals with breastfeeding information |  |  |
|  | Estimate | Std Error | P-value | Estimate | Std Error | P-value |
| Offspring effect | 0.030 | 0.02 | 0.137 | 0.029 | 0.02 | 0.146 |
| Prenatal maternal effect | 0.020 | 0.046 | 0.666 | 0.010 | 0.051 | 0.845 |
| Postnatal maternal effect | -0.017 | 0.034 | 0.620 | -0.007 | 0.041 | 0.862 |
| Postnatal paternal effect | 0.022 | 0.025 | 0.386 | 0.022 | 0.026 | 0.386 |

PRS: polygenic risk score; Std Error: standard error.

We therefore performed sensitivity analyses where we excluded adopted individuals who reported breastfeeding information (i.e. on the hypothesis that knowing if they were breast fed or not is information that an adopted individual could only know if they were adopted by a relative). The results of these analyses showed that estimates of the prenatal maternal genetic effect reduced in size and became non-significant. Whilst the postnatal maternal effect remained non-significant, the direction of effect changed from negative to positive. Sensitivity analyses using polygenic risk scores constructed using the 72 genome-wide significant SNPs from the Okbay et al. (2016) GWAS were underpowered to detect any effect of genetic risk score on educational attainment (Okbay et al., 2016).

### Simulations where adoptive parents are related to biological parents

Our simulations showed that the presence of adoptive parents who were genetically related to their adopted offspring could in some cases increase the type 1 error rate to detect prenatal maternal genetic effects and bias estimates of prenatal and postnatal maternal genetic effects when these relationships were not accurately modelled in the SEM (Supplementary Figures 3 to 6). In general, the effect of including unmodelled related adoptive and biological parents in the SEM depended on whether the adoptive mother or adoptive father was related to the biological parents (i.e. it did not matter whether the adoptive parents were related to the biological mother or father) and the degree of relatedness (i.e. closer relationships had the potential to produce greater bias and type I error rates).

In the case of adoptive mothers being genetically related to biological parents (Supplementary Figures 3 and 4), the presence of postnatal genetic effects (i.e. *β*_*m*_ not equal to 0) was sufficient to bias estimates of the prenatal maternal genetic effect (*γ*_*m*_) and increase type I error rates. In general, when *β*_*m*_ was not equal to zero, then estimates of *γ*_*m*_ were biased towards the true total maternal genetic effect (i.e. *γ*_*m*_ + *β*_*m*_), and estimates of *β*_*m*_ were biased towards zero. The reason for this can be seen intuitively by examining the path models for the G8 and G9 family structures in Supplementary Figure 1 where the presence of biologically related adoptive mothers leads to an unmodelled covariance path between offspring genotype and phenotype (i.e. 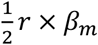). Note that the expected covariance implied by this path is the same in the case of adoptive mothers who are related to their adopted offsprings’ biological mothers (G8 in Supplementary Figure 1), and adoptive mothers who are related to their adopted offsprings’ biological fathers (G9 in Supplementary Figure 1)-implying that failure to model relatedness in both groups should produce the same consequences in terms of bias and type I error. When *β*_*m*_ is not zero, this unmodelled path alters the expected covariance between offspring genotype and phenotype in adopted singleton individuals. In this situation, the model will incorrectly attribute the altered covariance to prenatal maternal genetic effects (i.e. since the 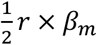 path is not explicitly modelled), and since estimates of the total maternal genetic effect remain unbiased (data not shown), will bias estimates of the postnatal maternal genetic effect towards zero. It follows that results will not be biased when *β*_*m*_ = 0 and the magnitude of any bias will decrease as *r* approaches zero (i.e. with decreasing genetic relationship between adoptive mothers and the biological parents).

Likewise, our simulations showed that the presence of paternal genetic effects (i.e. *β*_*p*_ not equal to 0) was sufficient to bias estimates of the prenatal maternal genetic effect (*γ*_*m*_) and increase type I error rates when unmodelled adoptive fathers who were genetically related to adopted children’s biological parents were included in the analysis (Supplementary Figures 5 and 6). Inspection of the G10 and G11 family structures in Supplementary Figure 1 shows that biologically related adoptive fathers leads to an extra unmodelled covariance path between offspring genotype and phenotype (i.e. 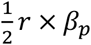), and that this path is the same for both adoptive fathers related to biological fathers (G10 in Supplementary Figure 1) and adoptive fathers related to biological mothers (G11 in Supplementary Figure 1) (again implying that failure to model relatedness in both groups should have the same consequences). It also shows that there should be no bias/inflation of type I error rates in the absence of paternal genetic effects, which is consistent with the results from our simulations. However, if *β*_*p*_ is not zero, then this will affect the covariance between (adopted) offspring genotype and phenotype, and changes may be falsely ascribed to prenatal maternal genetic effects. In this situation, estimates of the prenatal maternal genetic effect 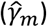 will be biased towards the sum of the true prenatal maternal genetic effect plus the true paternal genetic effect (*γ*_*m*_ + *β*_*p*_), and estimates of the postnatal maternal genetic effect 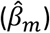 will be biased towards the difference between the true total maternal genetic effect (*γ*_*m*_ + *β*_*m*_) minus the estimated prenatal maternal genetic effect 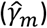.

In general, the presence of unmodelled relationships between biological and adoptive parents did not bias estimates of offspring genetic or paternal genetic effects (results not shown). This makes sense since offspring and paternal genetic effects are estimated directly from the covariance between observed genotypes and phenotypes (i.e. the covariance between observed offspring genotype and offspring phenotype, and the covariance between observed paternal genotype and offspring phenotype respectively), whereas the estimation of prenatal and postnatal maternal genetic effects have to be inferred indirectly using latent (not directly observed) genotypes and the difference in the covariance between offspring genotype and phenotype in biological and adopted families.

When we correctly modelled the relationship between biological and adoptive parents’ genotypes, there was no inflation in type 1 error rates and effect estimates were unbiased (Supplementary Figures 7 to 10). However, correct modelling and inclusion of these individuals in the SEM produced complicated effects on the power to detect true effects relative to if the same number of adopted individuals with biologically unrelated adoptive parents had been included in the model (see Supplementary Figure 11 where 0% on the x-axis corresponds to this situation). For the simulations we examined, power to detect prenatal and postnatal maternal genetic effects decreased when adopted singletons whose adoptive mothers were related to their biological parents were included (and modelled correctly) in the analysis. This makes sense intuitively in that correct modelling of these mothers requires a covariance path between adoptive mother’s genotype and the relevant biological parent’s genotype (G8, G9). The presence of this path decreases the difference between the expected covariance between an adopted offspring’s genotype and phenotype (G8, G9), and the expected covariance between offspring genotype and phenotype from a non-adopted family (G4)-hence lowering the power of the model to discriminate between maternal prenatal and postnatal genetic effects.

In contrast, power to detect prenatal and postnatal maternal genetic effects increased when adopted singletons whose adoptive fathers were related to their biological parents were included (and modelled correctly) in the analysis. In order to understand this result intuitively, it is useful to consider the case where the offspring trait is affected by concordant prenatal and postnatal maternal genetic effects (i.e. *γ*_*m*_ > 0 and *β*_*m*_ > 0), but not by paternal (*β*_*p*_= 0) or offspring genetic effects (*β*_*o*_ = 0). Under the full model, adopted singletons (G5) and adopted singletons whose adoptive father is a genetic relative (G10, G11) produce identical expected covariance matrices and so lead to identical model fits and parameter estimates. However, under the reduced model where *γ*_*m*_ is fixed to zero, the only path that accounts for the covariance between offspring genotype and phenotype is the offspring genetic effect (*β*_*o*_). Thus in order for this covariance to be modelled accurately, the reduced model will produce positive estimates for the offspring genetic effect (*β*_*o*_). Simultaneously, to ensure that the covariance between the observed paternal genotype and offspring phenotype remains close to zero in parent-offspring trios and father-offspring pairs (i.e. G1 and G3), the model will produce negative estimates of the paternal genetic effect (*β*_*p*_). This process presents more of a challenge when modelling adopted singleton individuals whose adoptive fathers are related to their biological parents (i.e. G10, G11) where the paternal genetic effect impacts both the expected covariance between parental genotype and offspring phenotype, and the expected covariance between offspring genotype and phenotype (i.e. because of the presence of the fixed covariance path *r*). The consequence is that the reduced model doesn’t fit the data as well and increased power to detect association. Similar thought experiments can be used to provide intuition for the other results in Supplementary Figure 11.

## Discussion

Genetic studies of adopted individuals have a venerable history in the field of behavior genetics (Leahy, 1935; Richardson, 1912-1913). Traditionally, adoption studies were used to determine the relative importance of genes and the environment on trait variability- the basic idea being that similarity between adopted individuals and their biological parents reflects the effect of genes, whereas the similarity between adopted individuals and their adoptive parents reflects the effect of shared environmental influences. Indeed, large studies of adopted individuals played a valuable role in resolving the “Nature vs Nurture” debate of last century, complementing similar findings from twin studies and other pedigree-based designs (Horn, 1983; Rhea et al., 2013; Scarr & Weinberg, 1983).

In this manuscript, we developed a new statistical model that uses the information provided by adopted individuals to partition maternal genetic effects into prenatal and postnatal influences on offspring phenotypes. Our method uses SEM to model the expected covariance between genotypes and phenotypes in adopted and non-adopted individuals. We have specifically designed our method to capitalize on the large number of adopted “singleton” individuals present in large-scale population-based resources like the UK Biobank, exploiting the fact that the expected covariance between offspring genotype and phenotype differs between adopted and non-adopted individuals as a function of postnatal maternal genetic (and other) effects.

We are not the first to note that information from adopted individuals could be used to provide valuable information on the effect of the prenatal environment on offspring traits (Loehlin, 2016). We are the first, however, to our knowledge to develop a model that specifically quantifies prenatal and postnatal maternal genetic effects on offspring phenotypes. Recently, Dominique and Fletcher (2020) constructed a genetic risk score to proxy educational attainment using maternal genotypes, and tested the strength of the association between the risk score and offspring educational attainment in children from biological and adoptive families separately (Domingue & Fletcher, 2020). The authors showed that the positive association between the score and educational attainment was stronger in children from biological than adoptive families. Our method differs from Dominique and Fletcher in a number of important respects. First, the focus in their study was on demonstrating indirect parental effects on offspring phenotypes, whereas the focus in our manuscript is specifically on partitioning maternal genetic effects into prenatal and postnatal effects. Second, our method provides point estimates and tests of the significance of prenatal and postnatal maternal genetic effects. Lastly, the method used in Dominique and Fletcher requires both adopted and biological parent-child trios (or alternatively mother-offspring pairs). In contrast, our method at a minimum requires biological parent-child trios (or both biological mother-offspring pairs and biological father-offspring pairs) and adopted singletons only to estimate prenatal and postnatal maternal genetic effects on offspring phenotypes. The inclusion of adoptive mother-adopted offspring pairs and/or biological mother-adopted offspring pairs increases power to partition maternal genetic effects dramatically, but is not a strict requirement of our method if adopted singletons are available.

Our power calculations show that in principle the number of adopted individuals in the UK Biobank is unlikely to be sufficient to resolve effects at individual genetic loci but may be large enough to detect evidence for prenatal/postnatal maternal genetic effects in the case of polygenic risk scores. It is worth noting that, whilst primarily a function of the number of adopted families in the analysis, the power to partition maternal genetic effects was also influenced by the number of biological families analysed. Intuitively, this is because the other family structures contribute to more precise estimation of e.g. paternal genetic effects, which in turn provides information on the total size of the maternal genetic component. This observation is important, because the number of adopted individuals in an analysis is likely to be severely constrained, whereas power can sometimes be improved (at least up to a threshold determined by the total number of adopted families/individuals in the analysis) by increasing the sample size of other relative types that may be easier to ascertain. Regardless, the most efficient way to increase the power to detect prenatal and/or postnatal maternal genetic effects is to genotype adoptive mother-adopted child pairs and/or biological mother-adopted child pairs.

Our empirical analyses in the UK Biobank showed the expected pattern of maternal genetic effects on offspring birth weight (i.e. strong evidence for prenatal but not postnatal maternal genetic effects) but not on offspring educational attainment. Parental educational attainment is known to causally affect offspring educational attainment (Bates et al., 2018; Hwang et al., 2020; Kong et al., 2018). Whilst it is possible, even probable, that some loci may affect offspring educational attainment through prenatal maternal pathways, it is not credible that such mechanisms would lead to larger effect sizes than those from postnatal pathways, or that fathers would exhibit strong evidence of postnatal effects but not mothers. Rather we hypothesize it is more likely that our model may have been misspecified in that substantial numbers of adopted individuals in the UK Biobank were in fact raised by their biological relatives. Sensitivity analyses that excluded adopted individuals with known breastfeeding information suggested that including adopted individuals who were raised by genetically related adoptive mothers may lead to inflated prenatal maternal genetic effect estimates. Our simulations with adoptive parents who were siblings or cousins of the biological parents further confirmed that unmodelled relatedness can increase evidence for prenatal maternal genetic effects at the expense of postnatal effects, particularly when paternal genetic and postnatal maternal effects are present, as would be the case with educational attainment (this contrasts with birthweight where we do not expect any paternal or postnatal genetic effects and for which we obtained sensible results). Ideally, one would have detailed information on any genetic relationship between adoptive and biological parents and model these relationships in the SEM (see below).

Our model relies on several strong assumptions, some of which are shown explicitly in Figure 1 and others which are not. Those assumptions explicitly encoded in Figure 1 include that the total maternal genetic effect can be decomposed into the sum of prenatal and postnatal components, that genetic effects are homogenous across biological and adopted families, the absence of genotype x environment interaction, any correlation between the genetic scores in spouses is the same in biological and adoptive families, and that fathers do not exert prenatal effects on offspring traits. Assumptions that are not explicitly shown in Figure 1 include that adopted (and adoptive) individuals are not systematically different from other individuals, adopted individuals are placed randomly within the population (including with individuals not genetically related to themselves), that adoption happened soon after birth and that the adopted individuals were raised in one adoptive family, and adopted individuals do not maintain contact with their biological parents. We also ignore any complexities due to e.g. single mothers raising offspring independently of their fathers etc.

Whilst we expect some of these assumptions to hold (i.e. it is reasonable to expect the absence of substantial prenatal paternal genetic effects for many traits), we expect that others will show varying degrees of violation with differing consequences for the parameter estimates obtained under the misspecified model. For example, it is well appreciated that adopted individuals and adoptive families may differ in several respects to the general population. For example, some studies have reported that adoptive families are better educated and have higher socio-economic status than the population average (Kendler et al., 2015). We argue that violation of this assumption is likely to exert more serious consequences on traditional adoption studies which model the phenotypic correlation between biological, adopted relatives. In contrast, in our design it is more important that genetic effect sizes are homogenous across adopted and non-adopted individuals (i.e. no genotype by environment interaction), and at least currently, there is limited evidence for large genotype by environment interactions at individual genetic loci. Our model also allows for differences in the residual variance across biological and adoptive families (Cheesman et al., 2020). This may be important if for example, adopted children are selectively placed in a reduced range of environments compared to their non-adopted counterparts.

We argue that of more consequence for the validity of our model is that any genetic relationship between adoptive and biological parents is accurately modelled and included in the SEM. Through simulation, we have shown that the consequences of model misspecification depend upon which biological and adoptive parents are related, the nature of this relationship, and the proportion of adopted individuals in the sample who have had their relationship misspecified. Our simulations also showed that correctly modelling this relationship returns unbiased effect estimates and correct type I error rates. Clearly, knowing these cryptic relationships in the UK Biobank would allow us to properly model them and better estimate prenatal and postnatal maternal genetic effects using this resource. We emphasize that accurately modelling these relationships does not require that actual genotypes for adoptive and/or biological parents be obtained (although this would be advantageous in terms of statistical power) as our SEM allows us to model these relationships in terms of latent variables.

We have also not modelled the complex effects of assortative mating in our SEM other than including covariance terms between maternal and paternal genotypes and assuming equal genetic variances in parents and their offspring under random mating (i.e. this is equivalent to assuming one round of assortative mating in the parental generation). Positive assortment induces a number of complications when attempting to decompose the offspring phenotypic variance into its constituent sources of variation, including increasing homozygosity and the genetic variance at loci for the trait undergoing assortment relative to that expected under Hardy-Weinberg equilibrium, and inducing correlations between assorting loci across the rest of the genome both within and between individuals in the same family. Recent work by Balboa et al (2021) and Kim et al (2021) have shown how genetic risk scores of transmitted and non-transmitted alleles in parent-offspring trios can be used to estimate direct and indirect genetic effects on offspring phenotype and the variation attributable to the environmental influence of parents on offspring under phenotypic assortment (Balbona et al., 2021; Kim et al., 2021). It is possible that our basic model could be extended in a similar fashion to incorporate the effect of assortment and estimate some of these effects also.

Finally, it has not been lost on us, that our framework may provide a basis for correcting Mendelian randomization (MR) studies of intrauterine/early life environmental exposures for some of the contaminating effects of horizontal genetic pleiotropy (Figure 4). The presence of latent horizontal pleiotropy is one of the major threats to the validity of causal inference from MR studies (Hemani et al., 2018). As we have shown, the inclusion of adopted individuals alongside biological families in genetic studies permits the partitioning of maternal genetic effects into prenatal and postnatal components. Consequently, if the focus of an MR study is on estimating the causal effect of a prenatal maternal exposure on an offspring outcome, the covariance between adoptive mother’s genotype and offspring (outcome) phenotype should provide an estimate of genetic effects due to postnatal horizontal genetic pleiotropy. These estimates could then be included in an MR model to correct causal estimates for the effect of postnatal pleiotropy (Figure 4). It is important to realize, however, that this procedure will not correct for the effect of prenatal/early life horizontal pleiotropy as these paths will be present in both biological and adopted individuals and so this approach is not a panacea for dealing with latent pleiotropy in MR studies.

In conclusion, we present a simple extension to the basic adoption design that includes measured genotypes and enables partitioning of maternal genetic effects into prenatal and postnatal sources of variation. We show in principle that adopted singleton individuals in the UK Biobank combined with biologically related parent-offspring trios and pairs is sufficient to partition maternal genetic effects into prenatal and postnatal sources of variation. However, power calculations suggest that such a partitioning would currently only be realistic for polygenic risk scores explaining substantial proportions of the variance in offspring phenotype, and that much larger numbers of individuals would be required to achieve such a partitioning at individual loci. In addition, failure to correctly model cryptic relationships between adoptive and biological parents may produce biased estimates of maternal genetic effects and increased type I error rates. Accurate modelling of cryptic relationships is sufficient to bring type I error rate under control and produce unbiased effect estimates. We suggest that there is a possibility that many individuals in the UK Biobank who report being adopted could have been raised by their biological relatives. We conclude that there would be considerable value in following up adopted individuals in the UK Biobank to determine whether they were raised by biological relatives, and if so, to precisely ascertain the nature of the relationship. These adopted individuals could then be used in informative statistical genetics models like the one described in the present manuscript to further elucidate the genetic architecture of complex traits and diseases.

## Acknowledgements

This research has been conducted using the UK Biobank resource (Reference 53641). D.M.E. is funded by an Australian National Health and Medical Research Council Senior Research Fellowship (APP1137714) and this work was funded by NHMRC project grants (GNT1157714, GNT1183074). G.H.M. is supported by the Norwegian Research Council (Post doctorial mobility research grant 287198) and Nils Normans minnegave.

---

**Figure 1.** Path diagrams illustrating the structural equation models (SEM) underlying the seven family structures modelled in this manuscript (G1 – G7). Causal relationships are presented by one headed arrows. Two headed arrows represents correlational relationships. Observed variables and latent variables are shown in squares and circles respectively. Z_M_ represents biological mother’s genotype which influences offspring phenotype (Y) via prenatal (γ_m_) and postnatal (β_m_) pathways. Z_P_ represents the biological father’s genotype which only influences offspring phenotype postnatally (β_p_). Z_o_ represents offspring genotype which influences offspring phenotype (β_o_) and is correlated ½ with the genotypes of its biological parents. Z_Mb_ represents the genotype of a biological mother whose child was adopted and therefore only influences her child’s phenotype through prenatal pathways (γ_m_). Z_Ma_ represents the adoptive mother’s genotype which only influences her adopted offspring’s phenotype via postnatal pathways (β_m_). Z_Pb_ represents the genotype of a biological father whose child was adopted and therefore has no influence on the adopted offspring phenotype. Z_Pa_ represents the adoptive father’s genotype which influences his adopted offspring postnatally (β_p_). *ρ* represents the covariance between parental genotypes, as a result of e.g. assortative mating (it is assumed that this covariance is the same in biological parents and adoptive parents). The total variance of genotypes in the parental generation is set to ∅. Ɛ_1_ and Ɛ_2_ represent residual error terms for the biological and adopted offspring phenotypes respectively that we assume have different variances.

**Figure 2.** Power to detect prenatal maternal genetic effects (γ_m_) (top) or postnatal maternal genetic effects (β_m_) (bottom) whilst also varying the size of prenatal and postnatal maternal genetic effects, paternal genetic effects (β_p_) or offspring genetic effects (β_o_). Effect sizes are parameterized using the path coefficients β and γ. Power was calculated assuming sample sizes approximating the number of white European individuals in the UK Biobank with educational attainment data (i.e. 1000 biological trios, 4000 biological mother-offspring pairs, 1800 biological father-offspring pairs, 300000 singletons, 6000 adopted individuals, and 50 biological mother-adopted offspring pairs and a covariance of 0 (ρ) between maternal and paternal genotypes).

**Figure 3.** Power to detect prenatal maternal (γ_m_) or postnatal maternal genetic (β_m_) effects whilst varying the numbers of each family structure, with the sample sizes of other family structures approximating numbers of white European individuals in the UK Biobank reporting educational attainment (1000 biological trios, 4000 biological mother-offspring pairs, 1800 biological father-offspring pairs, 300000 singletons, 6000 singletons, and 50 biological mother-adopted offspring pairs). Path coefficients representing postnatal or prenatal maternal genetic effects, paternal genetic effects (β_p_) and offspring genetic effects (β_o_) were fixed to 0.1. The covariance between maternal and paternal genotypes was fixed to 0.

**Figure 4.** Diagram illustrating a Mendelian randomization study designed to estimate the causal effect of a maternal prenatal environmental exposure on an offspring later life outcome. Maternal genotype proxies a perinatal exposure of interest and is used as an instrumental variable to estimate the causal effect of the perinatal exposure on the offspring outcome. The pathway of interest is represented by the blue arrows where the SNP influences the outcome prenatally through the perinatal exposure of interest. In this sort of design it is important to control for spurious pathways through the offspring genome since offspring genotype will be correlated ½ with maternal genotype. Maternal genotype could also influence the offspring phenotype via other pleiotropic paths through the intrauterine environment (red arrow) or through the postnatal environment (dashed arrows). The inclusion of adopted individuals into the research design may be useful in controlling for the effect of horizontal pleiotropic influences through the postnatal environment. Intuitively this is because adoptive mother’s genotype provides an estimate of the relationship between maternal genotype and offspring outcome through postnatal pathways only. This estimate could be included in statistical models of the relationship between maternal genotype and offspring outcome to help correct for the effect of horizontal pleiotropy.

## Notes

### Competing Interest Statement

The authors have declared no competing interest.

