## Supplementary material for "Using adopted individuals to partition maternal genetic effects into prenatal and postnatal effects on offspring phenotypes": Figures

### Slide 1
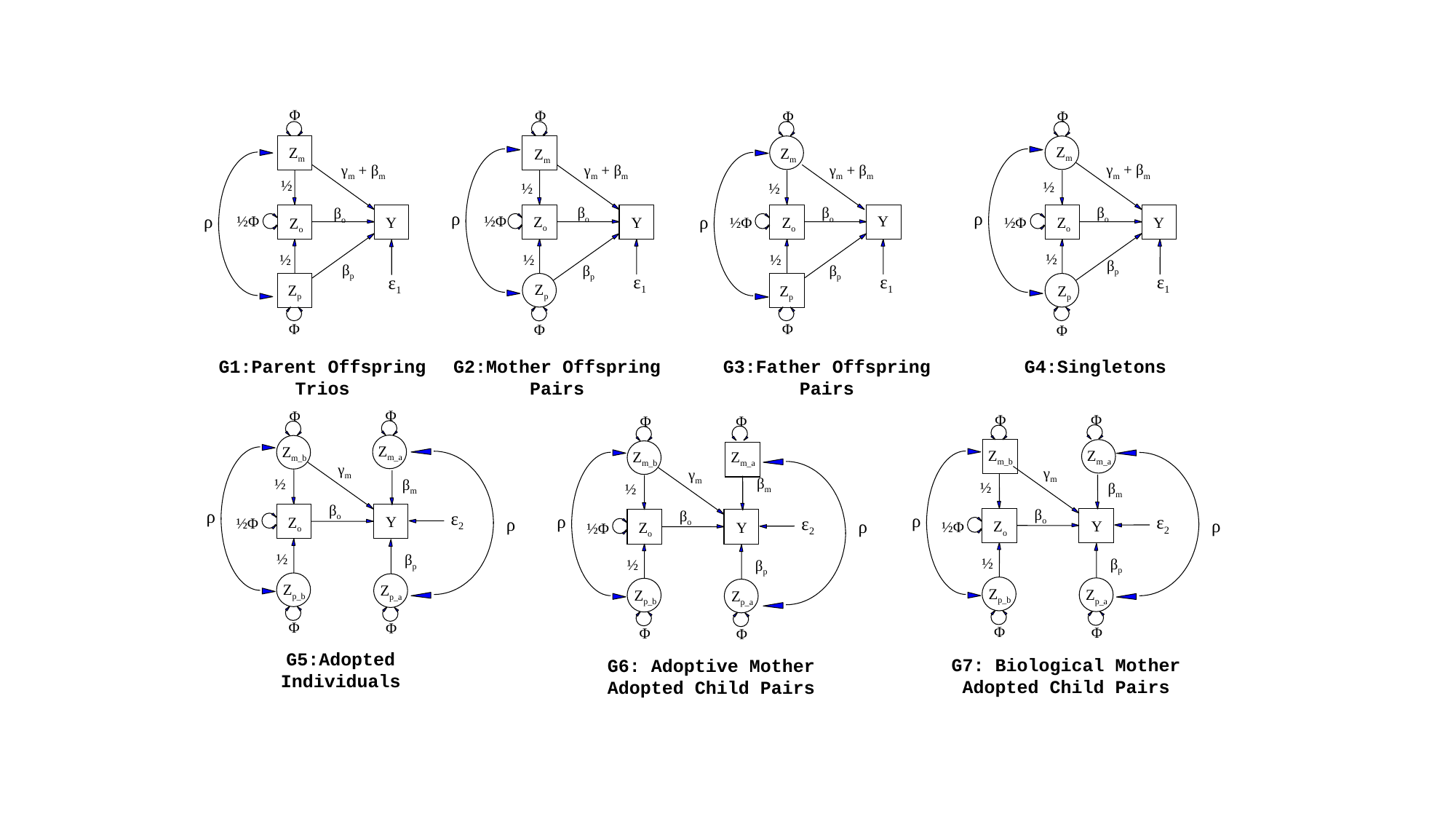

Φ
Z_m_
γ_m_ + β_m_
½
β_o_
½Φ
Y
Z_o_
½
β_p_
ε_1_
Z_p_
Φ
Φ
Z_m_
γ_m_ + β_m_
½
β_o_
Z_o_
Y
½
β_p_
ε_1_
Z_p_
Φ
Φ
Φ
Z_m_
Z_m_
ρ
ρ
ρ
ρ
γ_m_ + β_m_
γ_m_ + β_m_
½
½
β_o_
β_o_
½Φ
Y
Z_o_
½Φ
Y
Z_o_
½Φ
½
½
β_p_
β_p_
ε_1_
ε_1_
Z_p_
Z_p_
Φ
Φ
G1:Parent Offspring
Trios
G2:Mother Offspring
Pairs
G3:Father Offspring
Pairs
G4:Singletons
Φ
Φ
Φ
Φ
Φ
Φ
Z_m_a_
Z_m_b_
Z_m_a_
Z_m_b_
Z_m_a_
Z_m_b_
ρ
ρ
ρ
γ_m_
γ_m_
γ_m_
β_m_
½
β_m_
½
β_m_
½
β_o_
β_o_
β_o_
ε_2_
ε_2_
ε_2_
Y
Z_o_
ρ
½Φ
ρ
ρ
Y
Z_o_
½Φ
Y
Z_o_
½Φ
½
β_p_
½
β_p_
½
β_p_
Z_p_b_
Z_p_a_
Z_p_b_
Z_p_a_
Z_p_b_
Z_p_a_
Φ
Φ
Φ
Φ
Φ
Φ
G5:Adopted
Individuals
G7: Biological Mother
Adopted Child Pairs
G6: Adoptive Mother
Adopted Child Pairs

### Slide 2
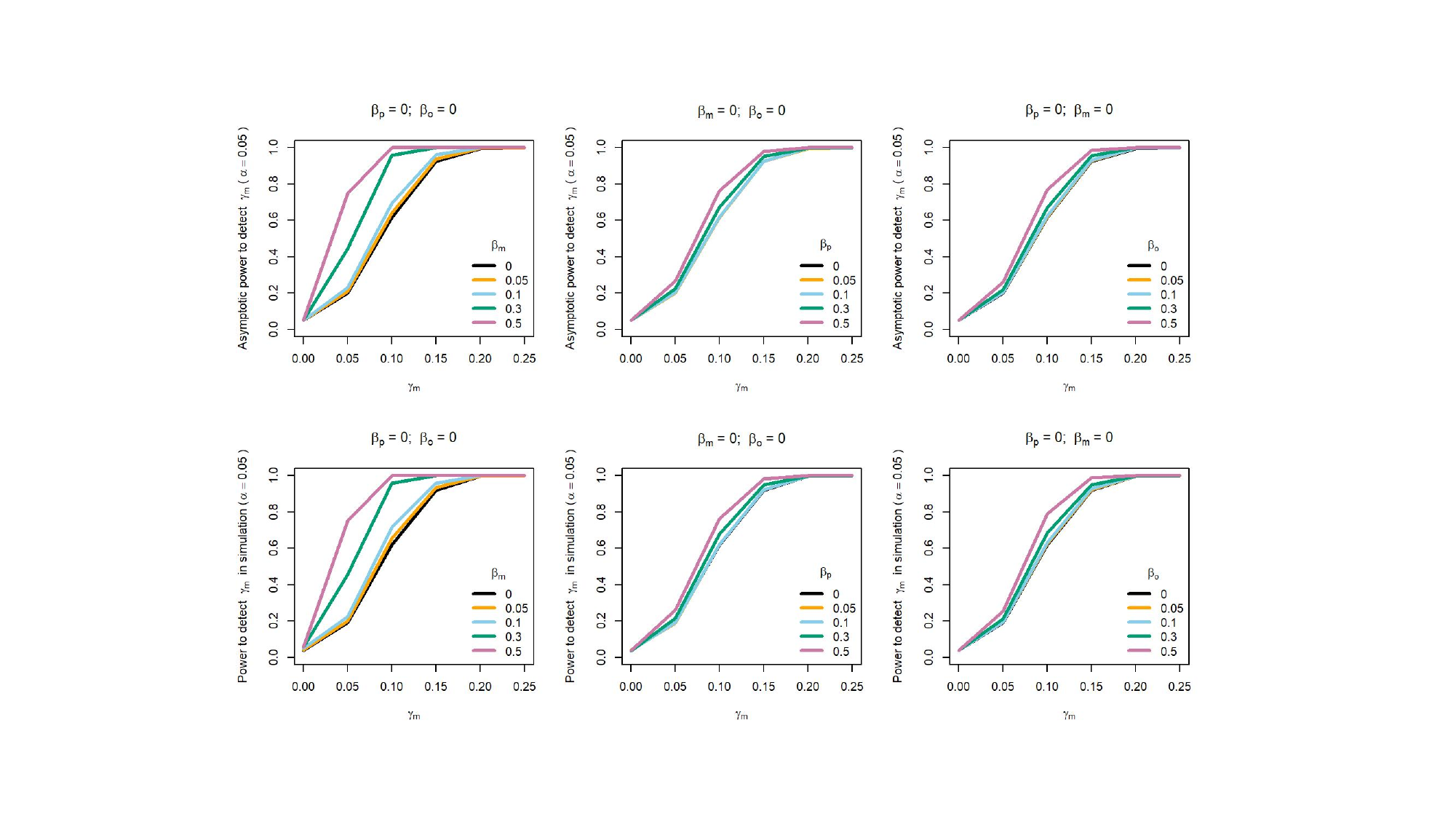

### Slide 3
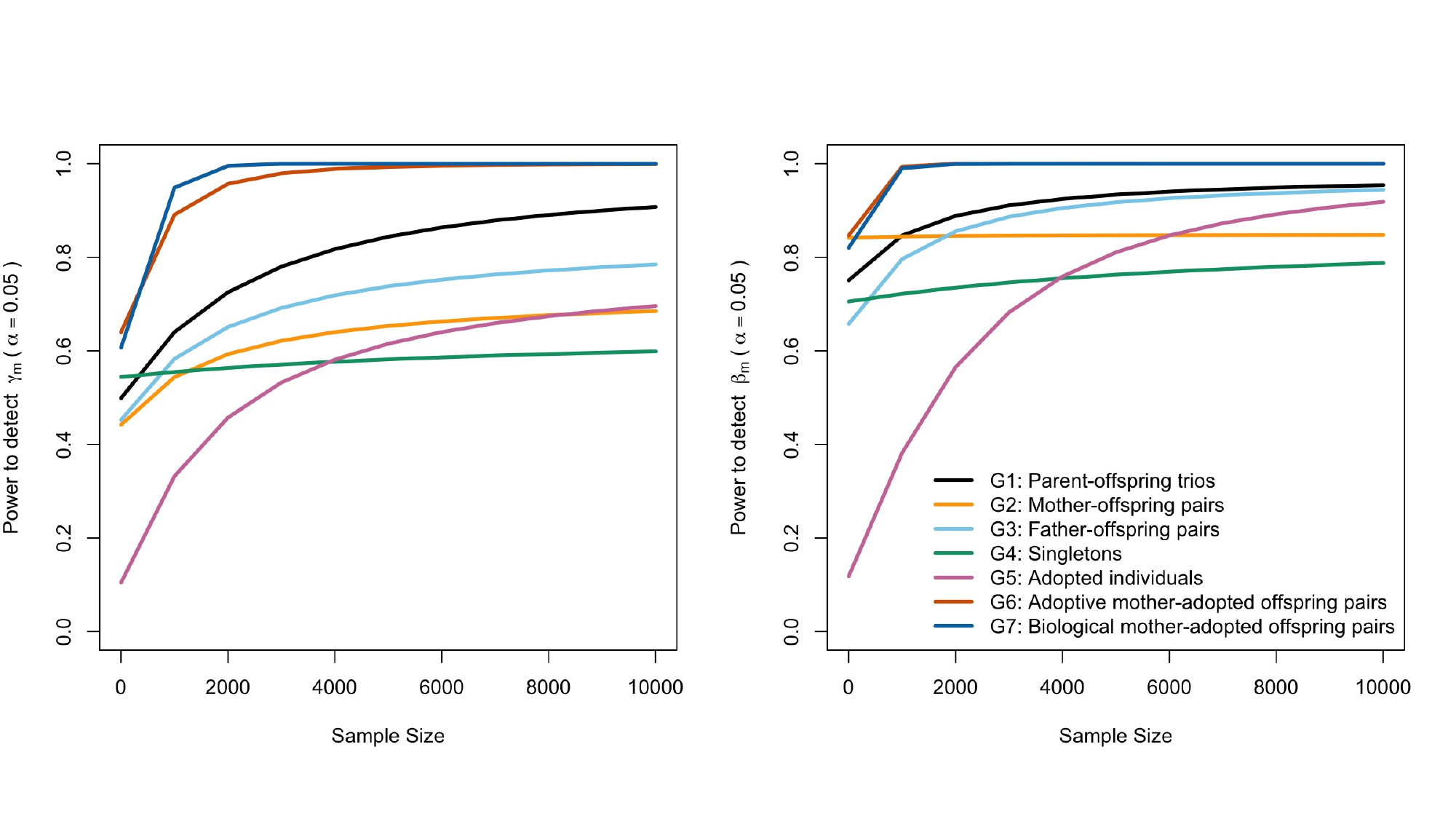

### Slide 4
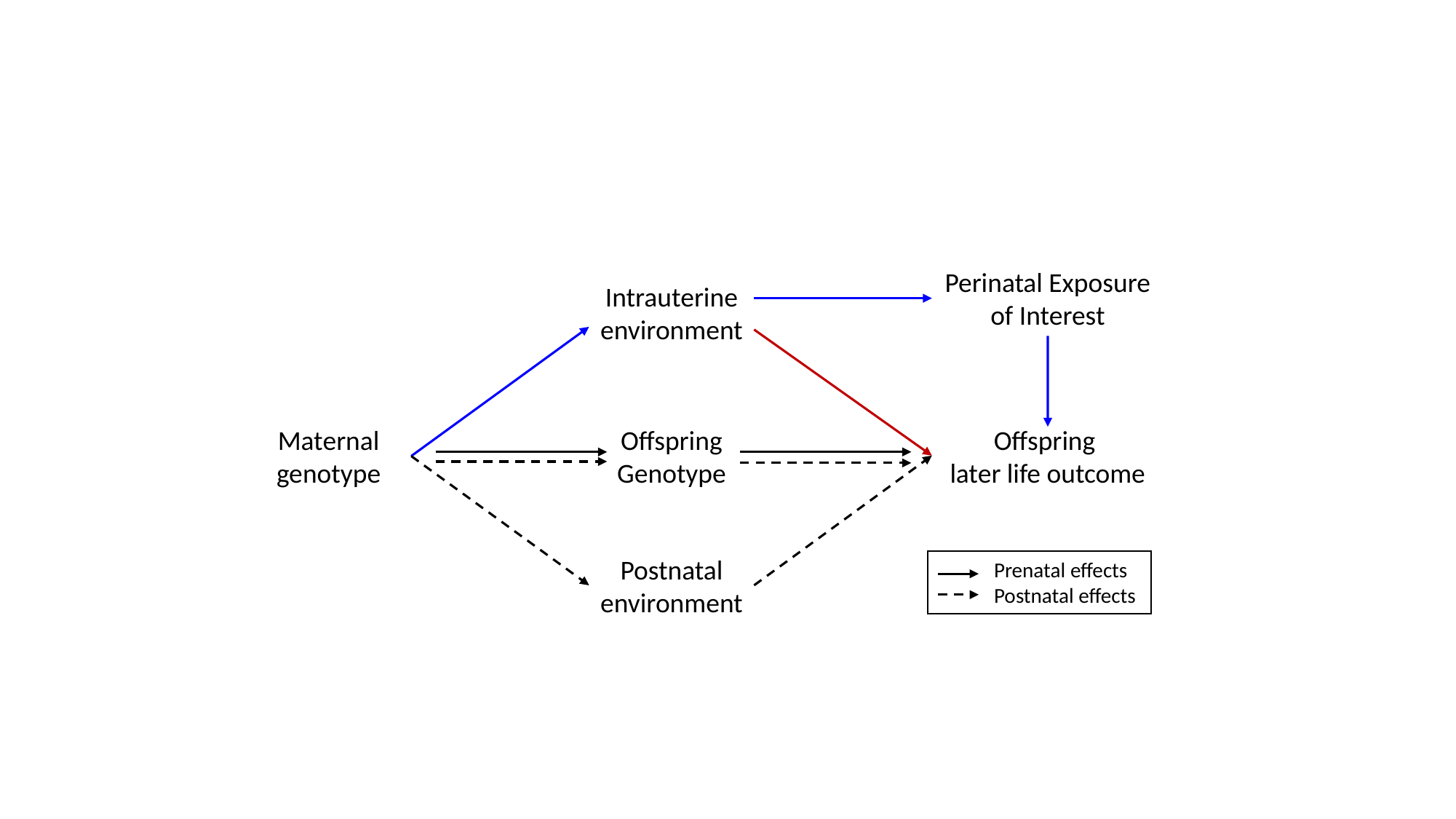

Perinatal Exposure of Interest
Intrauterine environment
Offspring Genotype
Maternal genotype
Offspring
later life outcome
Postnatal environment
Prenatal effects
Postnatal effects
