## Supplementary Figures for "Using adopted individuals to partition maternal genetic effects into prenatal and postnatal effects on offspring phenotypes"

### Slide 1
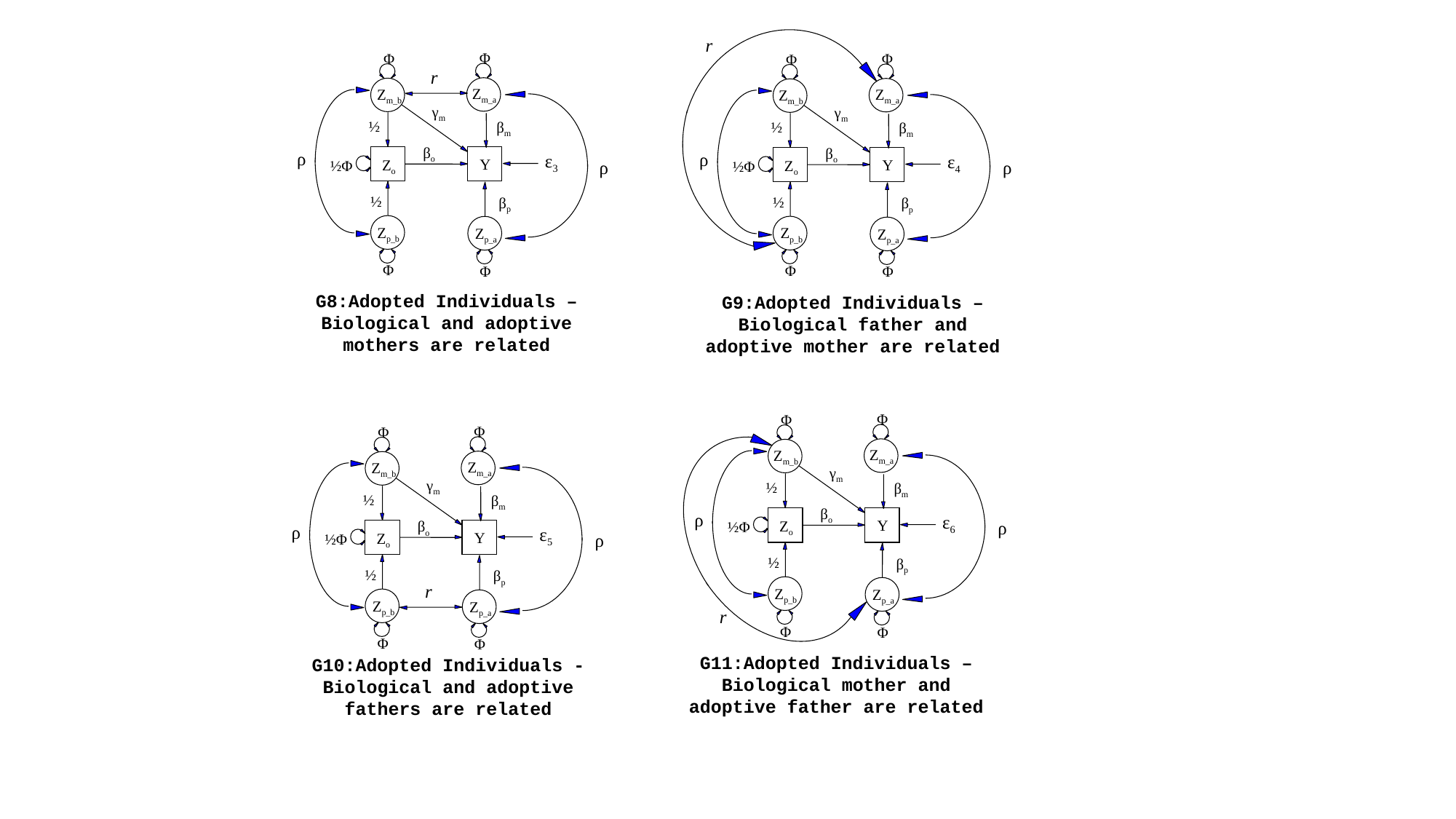

r
Φ
Φ
Φ
Φ
r
Z_m_a_
Z_m_b_
Z_m_a_
Z_m_b_
ρ
ρ
γ_m_
γ_m_
½
β_m_
½
β_m_
β_o_
β_o_
ε_3_
ε_4_
Y
Z_o_
Y
½Φ
ρ
Z_o_
ρ
½Φ
½
½
β_p_
β_p_
Z_p_b_
Z_p_b_
Z_p_a_
Z_p_a_
Φ
Φ
Φ
Φ
G8:Adopted Individuals – Biological and adoptive mothers are related
G9:Adopted Individuals – Biological father and adoptive mother are related
Φ
Φ
Φ
Φ
Z_m_a_
Z_m_b_
ρ
Z_m_a_
Z_m_b_
γ_m_
ρ
γ_m_
½
β_m_
½
β_m_
β_o_
ε_6_
Y
Z_o_
ρ
β_o_
½Φ
ε_5_
Y
Z_o_
ρ
½Φ
½
β_p_
½
β_p_
r
Z_p_b_
Z_p_a_
Z_p_b_
Z_p_a_
r
Φ
Φ
Φ
Φ
G11:Adopted Individuals – Biological mother and adoptive father are related
G10:Adopted Individuals - Biological and adoptive fathers are related

### Slide 2
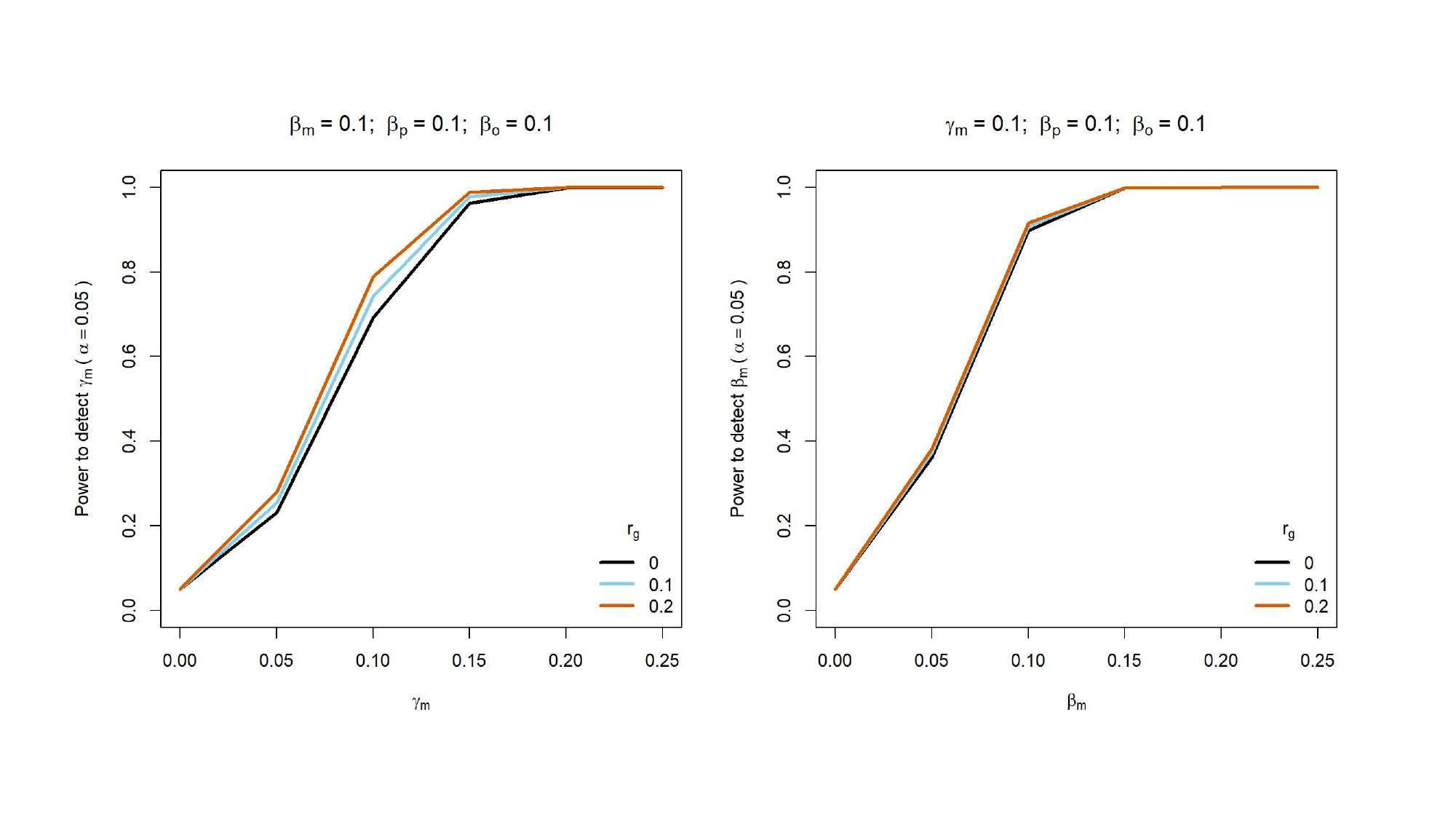

### Slide 3
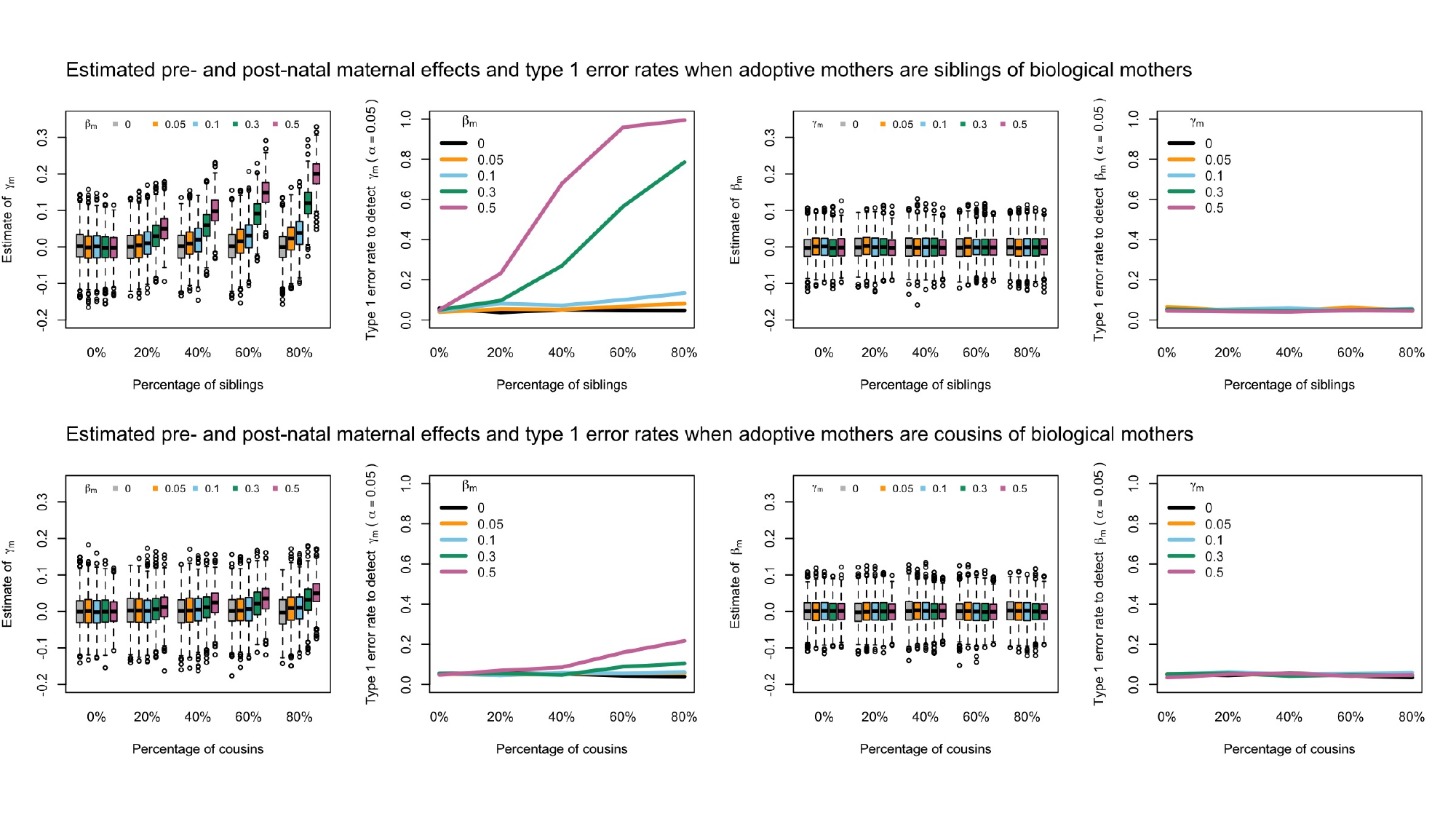

### Slide 4
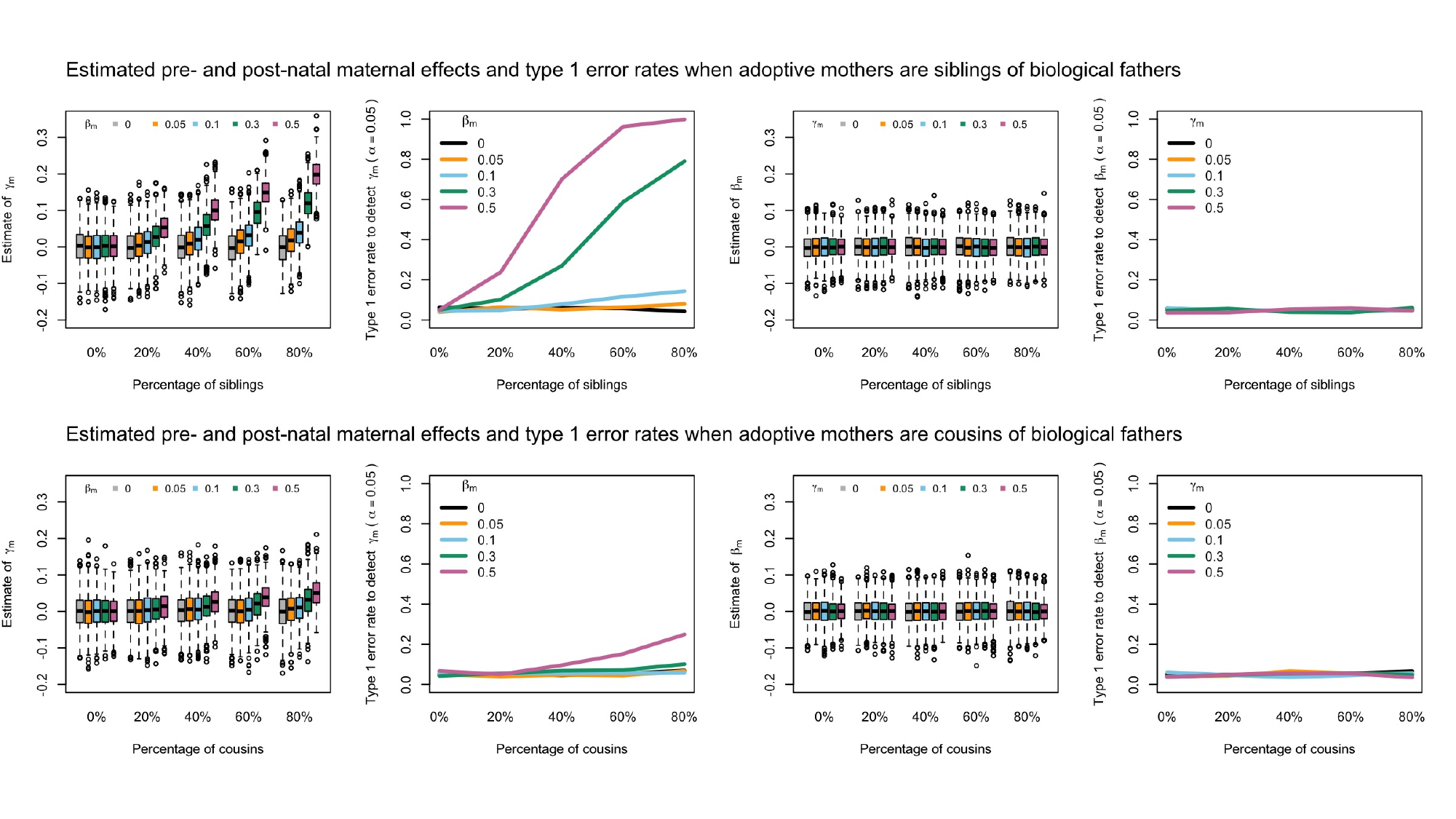

### Slide 5
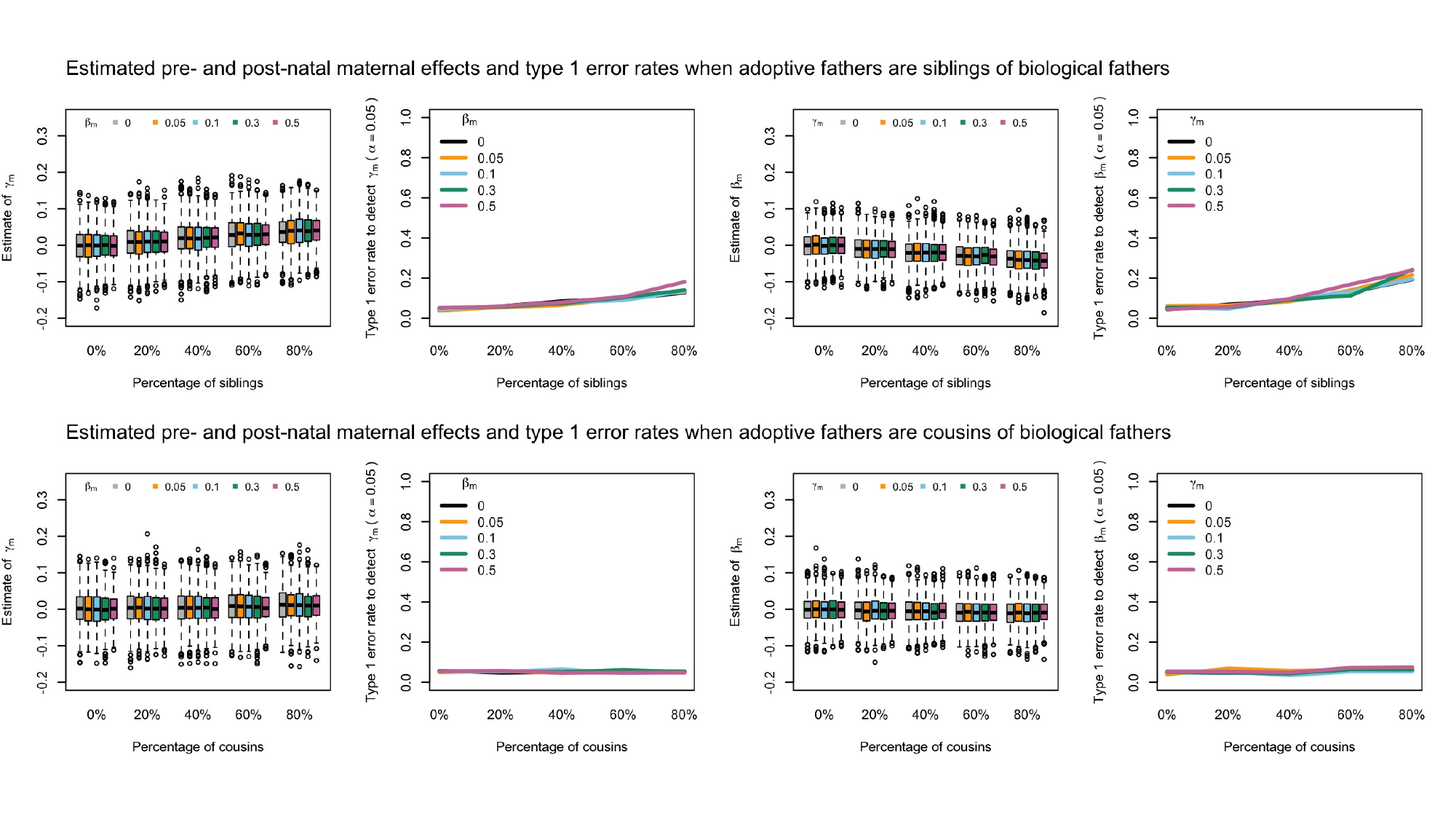

### Slide 6
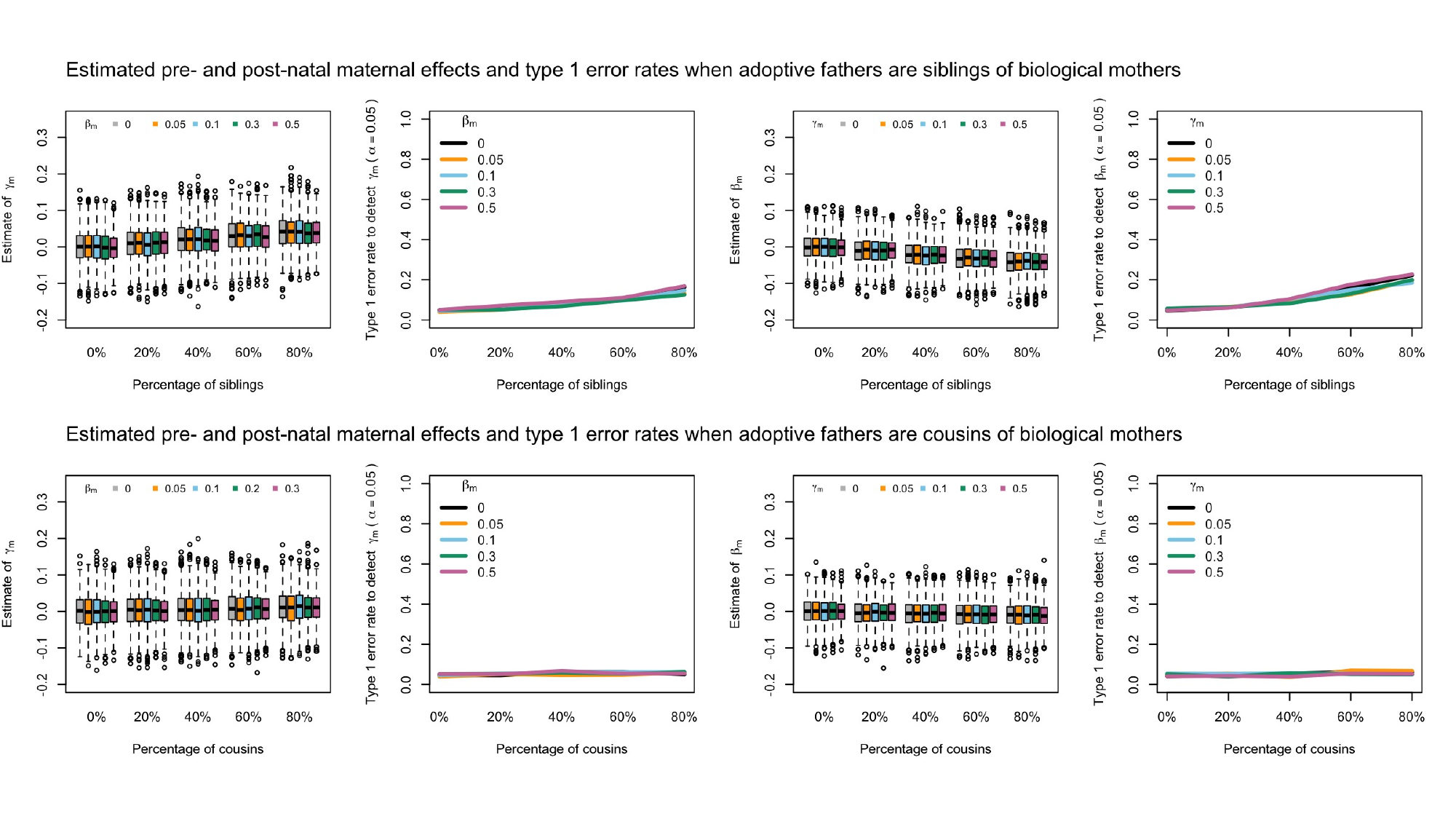

### Slide 7
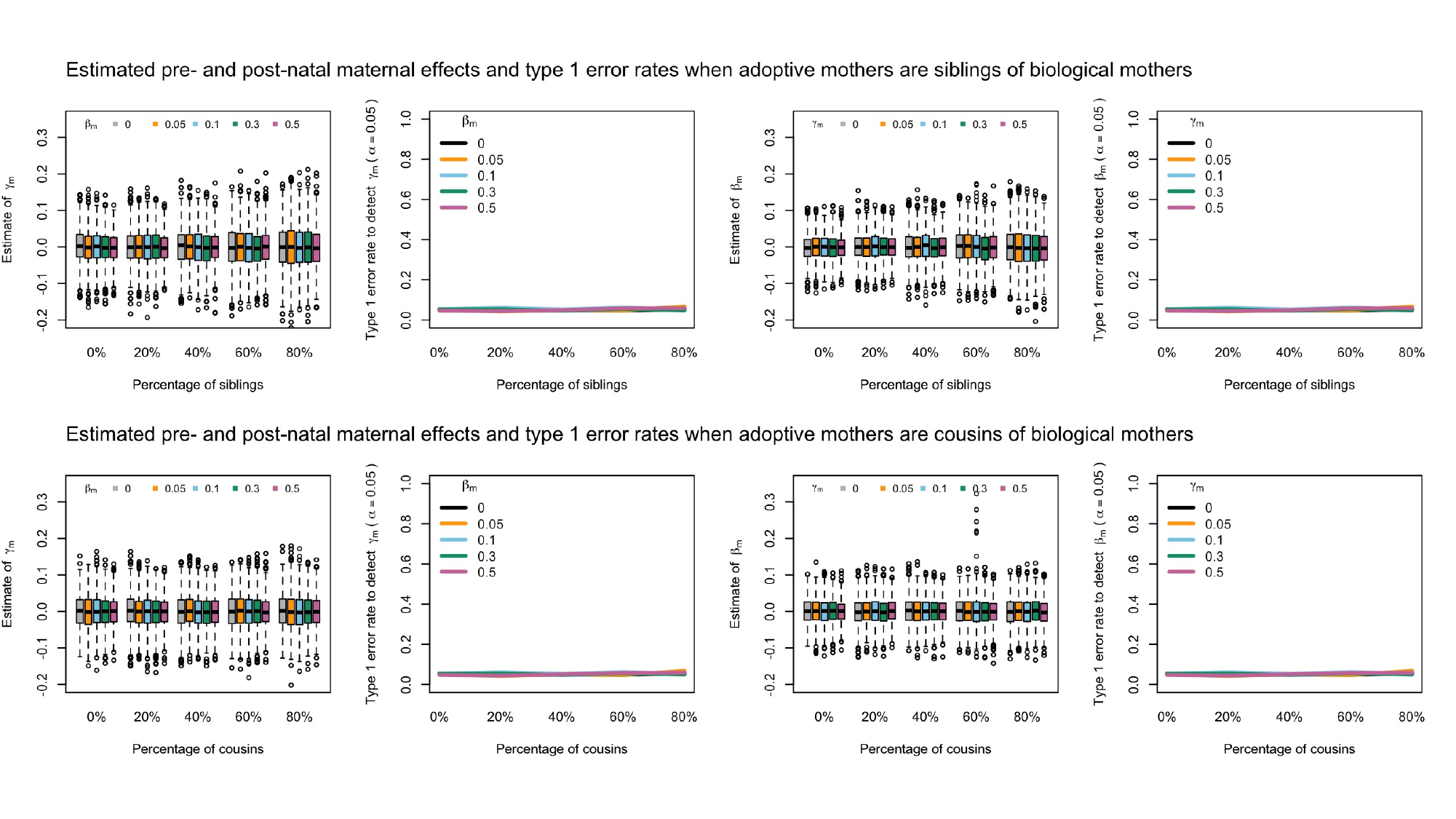

### Slide 8
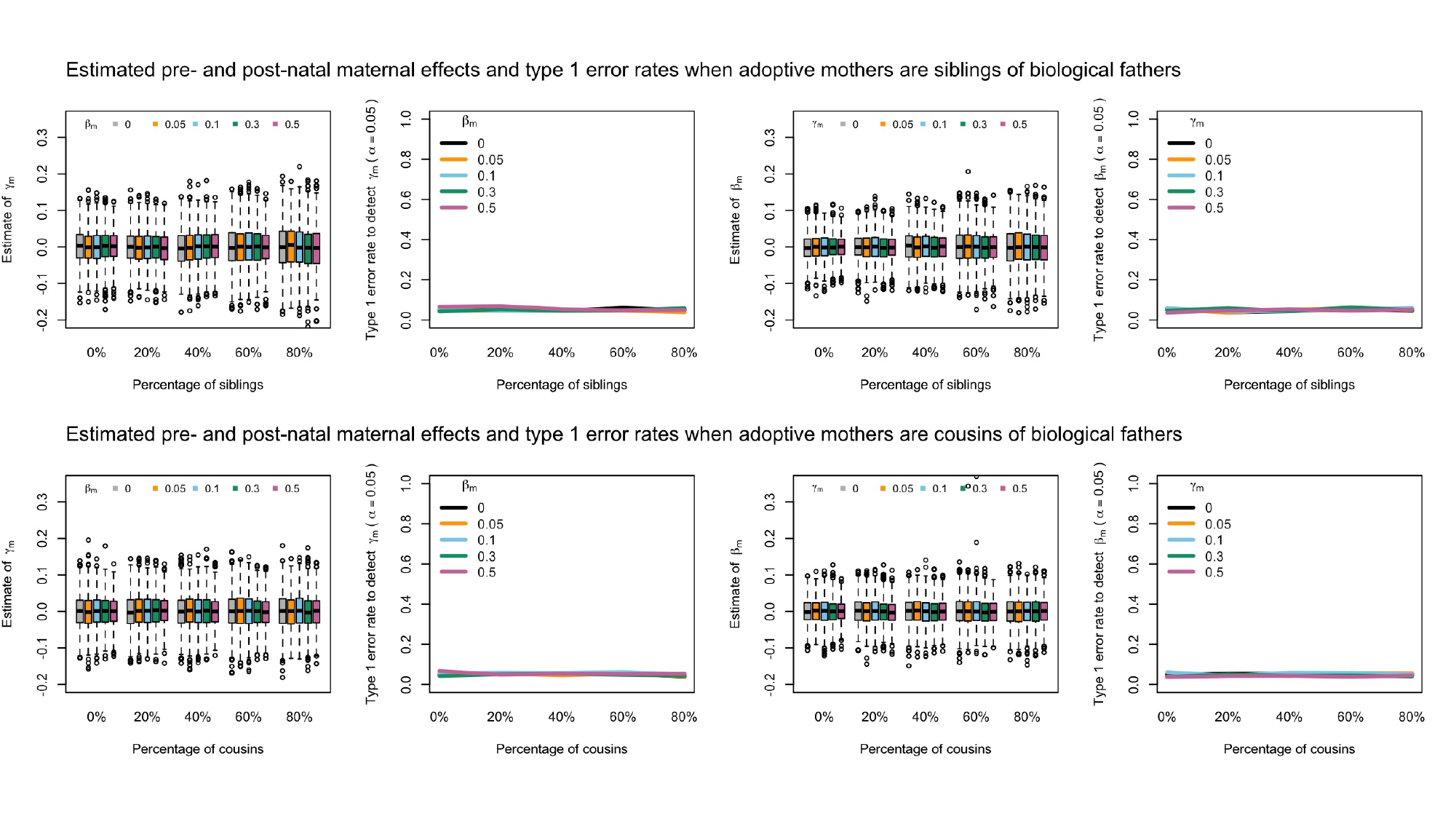

### Slide 9
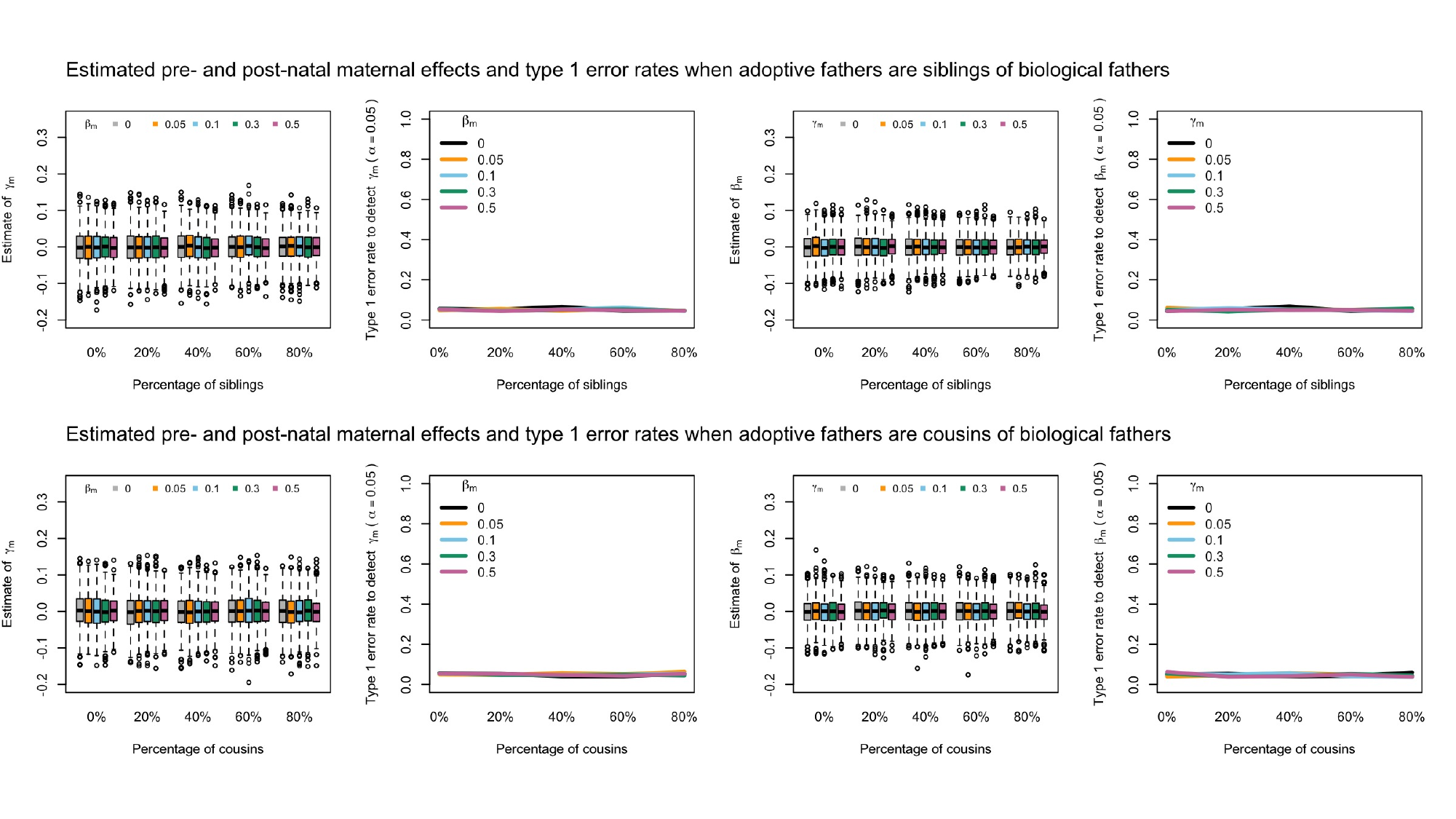

### Slide 10
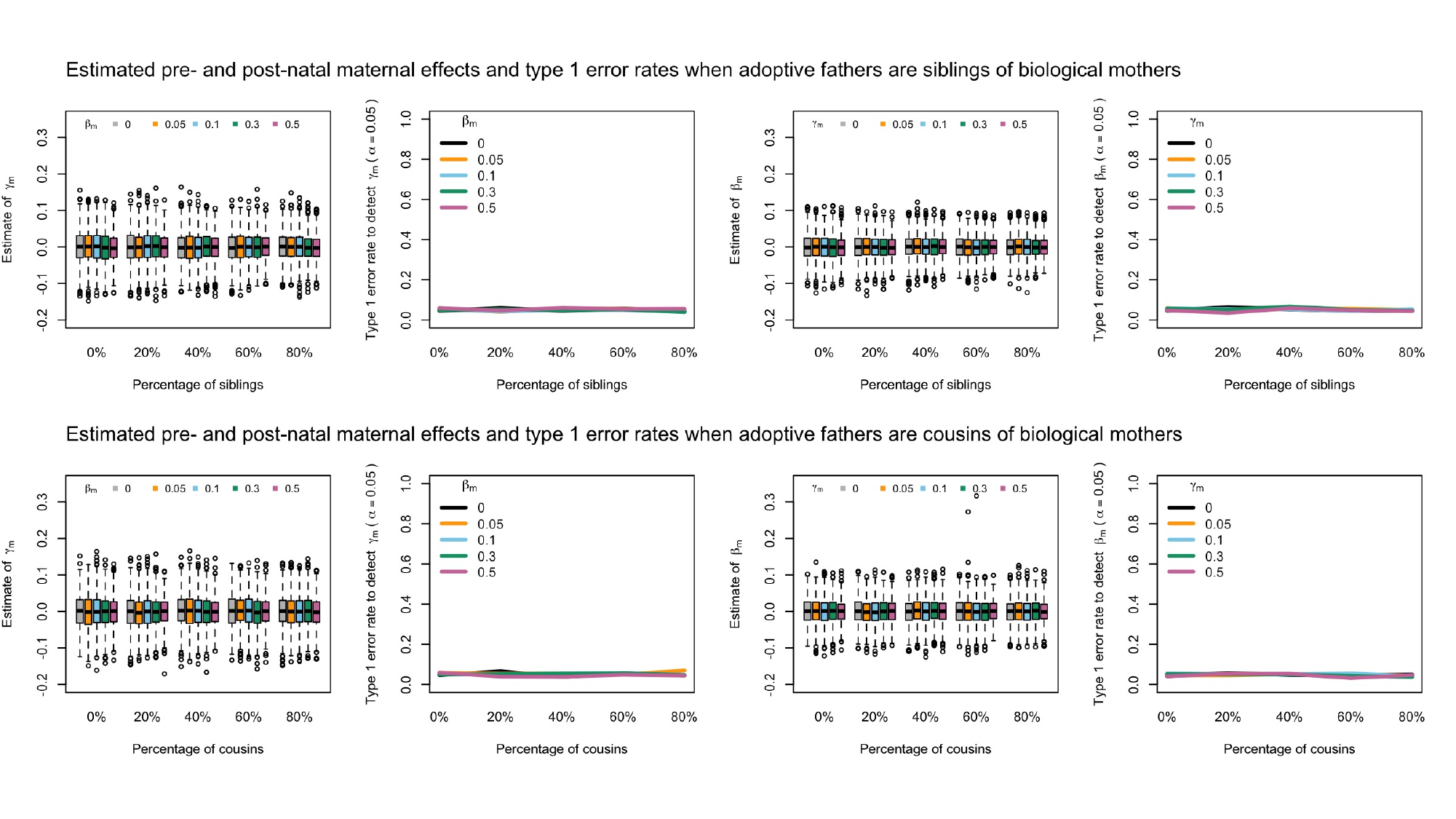

### Slide 11
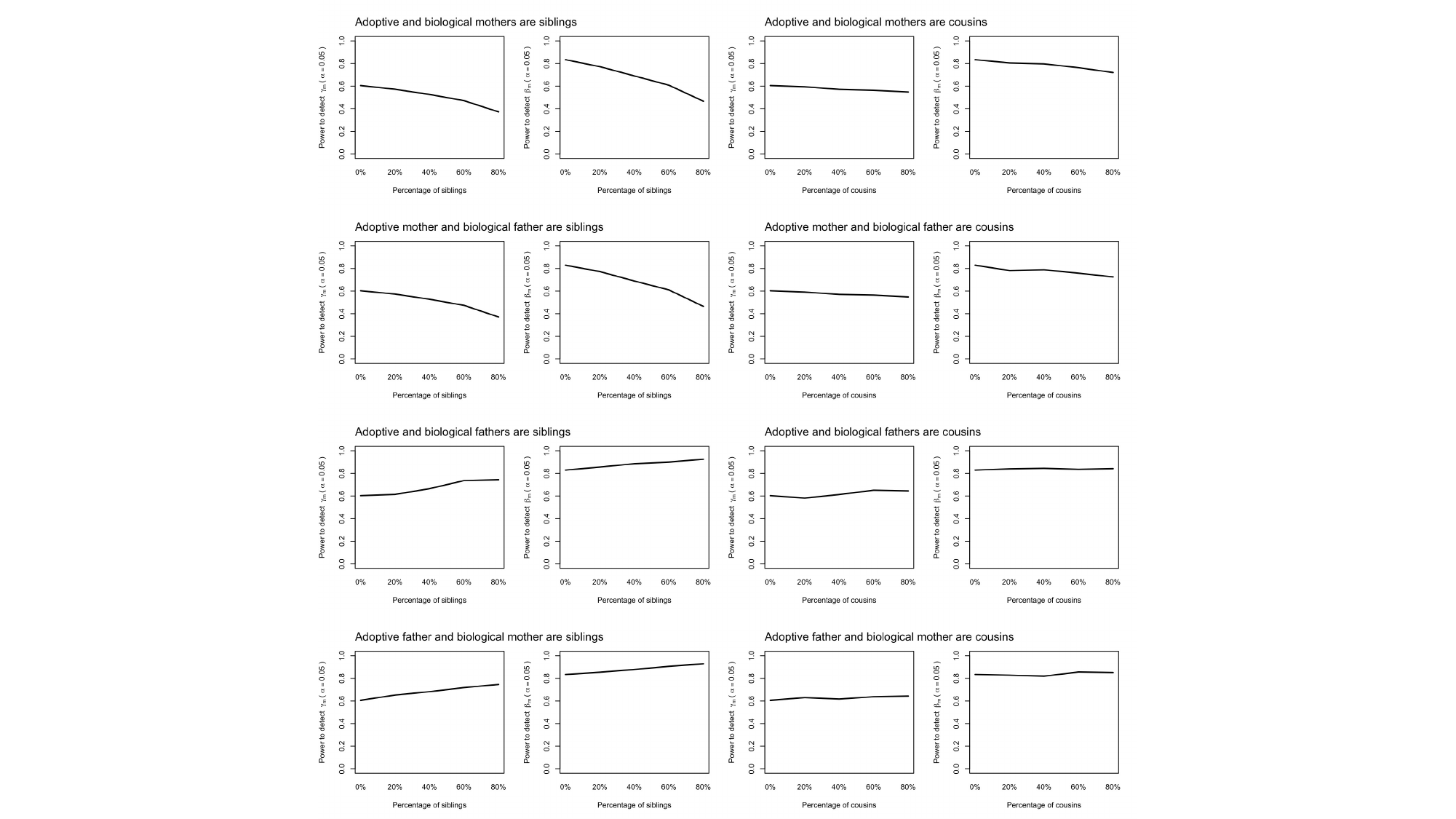
