## Supplementary Materials for "Using adopted individuals to partition maternal genetic effects into prenatal and postnatal effects on offspring phenotypes"

**Legends to Supplementary Figures**

**Supplementary Figure 1.** Path diagrams illustrating the structural equation models (SEM) underlying four additional family structures (G8 – G11) where biological and adoptive parents are related. Observed variables and latent variables are shown in squares and circles respectively. Causal relationships are presented by one headed arrows. Two headed arrows represents correlational relationships. Z_M_ represents biological mother’s genotype which influences offspring phenotype (Y) via prenatal (γ_m_) and postnatal (β_m_) pathways. Z_P_ represents the biological father’s genotype which only influences offspring phenotype postnatally (β_p_). Z_o_ represents offspring genotype which influences offspring phenotype (β_o_) and is correlated ½ with the genotypes of its biological parents. Z_Mb_ represents the genotype of a biological mother whose child was adopted and therefore only influences her child’s phenotype through prenatal pathways (γ_m_). Z_Ma_ represents the adoptive mother’s genotype which only influences her adopted offspring’s phenotype via postnatal pathways (β_m_). Z_Pb_ represents the genotype of a biological father whose child was adopted and therefore has no influence on the adopted offspring phenotype. Z_Pa_ represents the adoptive father’s genotype which influences his adopted offspring postnatally (β_p_). *ρ* represents the covariance between parental genotypes, as a result of e.g. assortative mating (it is assumed that this covariance is the same in biological parents and adoptive parents). The total variance of genotypes in the parental generation is set to Φ. Ɛ_3_ through Ɛ_6_ represent residual error terms for the adopted offspring that we assume have different variances. The (fixed) parameter *r* represents the covariance between biological and adoptive parents’ genotypes.

**Supplementary Figure 2.** Power to detect prenatal maternal (γ_m_) or postnatal maternal (β_m_) genetic effects whilst varying the correlation (r_g_) between maternal and paternal genotypes. Power was calculated using sample sizes approximating the number of white European individuals in the UK Biobank reporting their own educational attainment (1000 biological trios, 4000 biological mother-offspring pairs, 1800 biological father-offspring pairs, 300000 singletons, 6000 adopted individuals, and 50 biological mother-adopted offspring pairs). Path coefficients representing prenatal maternal (γ_m_), postnatal maternal (β_m_), paternal (β_p_) and offspring genetic effects (β_o_) were fixed to 0.1.

**Supplementary Figure 3.** Estimated prenatal (γ_m_) and postnatal (β_m_) maternal effects and type 1 error rates whilst varying the percentage of adoptive mothers being siblings (top) or cousins (bottom) of biological mothers. An SEM which did not correctly model this relationship was fit to the data. Power was calculated using simulated data with 1000 biological trios, 4000 biological mother-offspring pairs, 1800 biological father-offspring pairs, 300000 singletons, and 6000 adopted individuals. Prenatal maternal genetic effects were fixed to zero whilst the size of postnatal maternal genetic effects were varied (left panels); postnatal maternal genetic effects were fixed to zero whilst the size of prenatal maternal genetic effects were varied (right panels). Paternal effects (β_p_) and offspring effects (β_o_) were fixed to 0.1. The covariance between maternal and paternal genotypes was fixed to zero.

**Supplementary Figure 4.** Estimated prenatal (γ_m_) and postnatal (β_m_) maternal effects and type 1 error rates whilst varying the percentage of adoptive mothers being siblings (top) or cousins (bottom) of biological fathers. An SEM which did not correctly model this relationship was fit to the data. Power was calculated using simulated data with 1000 biological trios, 4000 biological mother-offspring pairs, 1800 biological father-offspring pairs, 300000 singletons, and 6000 adopted individuals. Prenatal maternal genetic effects were fixed to zero whilst the size of postnatal maternal genetic effects were varied (left panels); postnatal maternal genetic effects were fixed to zero whilst the size of prenatal maternal genetic effects were varied (right panels). Paternal effects (β_p_) and offspring effects (β_o_) were fixed to 0.1. The covariance between maternal and paternal genotypes was fixed to zero.

**Supplementary Figure 5.** Estimated prenatal (γ_m_) and postnatal (β_m_) maternal effects and type 1 error rates whilst varying the percentage of adoptive fathers being siblings (top) or cousins (bottom) of biological fathers. An SEM which did not correctly model this relationship was fit to the data. Power was calculated using simulated data with 1000 biological trios, 4000 biological mother-offspring pairs, 1800 biological father-offspring pairs, 300000 singletons, and 6000 adopted individuals. Prenatal maternal genetic effects were fixed to zero whilst the size of postnatal maternal genetic effects were varied (left panels); postnatal maternal genetic effects were fixed to zero whilst the size of prenatal maternal genetic effects were varied (right panels). Paternal effects (β_p_) and offspring effects (β_o_) were fixed to 0.1. The covariance between maternal and paternal genotypes was fixed to zero.

**Supplementary Figure 6.** Estimated prenatal (γ_m_) and postnatal (β_m_) maternal effects and type 1 error rates whilst varying the percentage of adoptive fathers being siblings (top) or cousins (bottom) of biological mothers. An SEM which did not correctly model this relationship was fit to the data. Power was calculated using simulated data with 1000 biological trios, 4000 biological mother-offspring pairs, 1800 biological father-offspring pairs, 300000 singletons, and 6000 adopted individuals. Prenatal maternal genetic effects were fixed to zero whilst the size of postnatal maternal genetic effects were varied (left panels); postnatal maternal genetic effects were fixed to zero whilst the size of prenatal maternal genetic effects were varied (right panels). Paternal effects (β_p_) and offspring effects (β_o_) were fixed to 0.1. The covariance between maternal and paternal genotypes was fixed to zero.

**Supplementary Figure 7.** Estimated prenatal (γ_m_) and postnatal (β_m_) maternal effects and type 1 error rates whilst varying the percentage of adoptive mothers being siblings (top) or cousins (bottom) of biological mothers. An SEM which modelled this relationship correctly was fit to the data. Power was calculated using simulated data with 1000 biological trios, 4000 biological mother-offspring pairs, 1800 biological father-offspring pairs, 300000 singletons, and 6000 adopted individuals. Prenatal maternal genetic effects were fixed to zero whilst the size of postnatal maternal genetic effects were varied (left panels); postnatal maternal genetic effects were fixed to zero whilst the size of prenatal maternal genetic effects were varied (right panels). Paternal effects (β_p_) and offspring effects (β_o_) were fixed to 0.1. The covariance between maternal and paternal genotypes was fixed to zero.

**Supplementary Figure 8.** Estimated prenatal (γ_m_) and postnatal (β_m_) maternal effects and type 1 error rates whilst varying the percentage of adoptive mothers being siblings (top) or cousins (bottom) of biological fathers. An SEM which modelled this relationship correctly was fit to the data. Power was calculated using simulated data with 1000 biological trios, 4000 biological mother-offspring pairs, 1800 biological father-offspring pairs, 300000 singletons, and 6000 adopted individuals. Prenatal maternal genetic effects were fixed to zero whilst the size of postnatal maternal genetic effects were varied (left panels); postnatal maternal genetic effects were fixed to zero whilst the size of prenatal maternal genetic effects were varied (right panels). Paternal effects (β_p_) and offspring effects (β_o_) were fixed to 0.1. The covariance between maternal and paternal genotypes was fixed to zero.

**Supplementary Figure 9.** Estimated prenatal (γ_m_) and postnatal (β_m_) maternal effects and type 1 error rates whilst varying the percentage of adoptive fathers being siblings (top) or cousins (bottom) of biological fathers. An SEM which modelled this relationship correctly was fit to the data. Power was calculated using simulated data with 1000 biological trios, 4000 biological mother-offspring pairs, 1800 biological father-offspring pairs, 300000 singletons, and 6000 adopted individuals. Prenatal maternal genetic effects were fixed to zero whilst the size of postnatal maternal genetic effects were varied (left panels); postnatal maternal genetic effects were fixed to zero whilst the size of prenatal maternal genetic effects were varied (right panels). Paternal effects (β_p_) and offspring effects (β_o_) were fixed to 0.1. The covariance between maternal and paternal genotypes was fixed to zero.

**Supplementary Figure 10.** Estimated prenatal (γ_m_) and postnatal (β_m_) maternal effects and type 1 error rates whilst varying the percentage of adoptive fathers being siblings (top) or cousins (bottom) of biological mothers. An SEM which modelled this relationship correctly was fit to the data. Power was calculated using simulated data with 1000 biological trios, 4000 biological mother-offspring pairs, 1800 biological father-offspring pairs, 300000 singletons, and 6000 adopted individuals. Prenatal maternal genetic effects were fixed to zero whilst the size of postnatal maternal genetic effects were varied (left panels); postnatal maternal genetic effects were fixed to zero whilst the size of prenatal maternal genetic effects were varied (right panels). Paternal effects (β_p_) and offspring effects (β_o_) were fixed to 0.1. The covariance between maternal and paternal genotypes was fixed to zero.

**Supplementary Figure 11.** Power to detect prenatal (γ_m_) and postnatal maternal (β_m_) genetic effects when the relationship between adoptive and biological parents are correctly specified in the model. Power was calculated using simulated data assuming 1000 biological parent-offspring trios, 4000 biological mother-offspring pairs, 1800 biological father-offspring pairs, 300,000 singletons from non-adopted families, and 6000 adopted individuals. Prenatal maternal genetic effects , postnatal maternal genetic effects, paternal genetic effects (β_p_) and offspring genetic effects (β_o_) were fixed to 0.1. The covariance between maternal and paternal genotypes was fixed to 0. The percentage of adopted individuals whose adoptive parents were genetically related to their biological parents was varied (x-axis). Note that 0% corresponds to the situation where biological and adoptive parents are unrelated.

\# This is code to partition maternal genetic effects into pre- and postnatal effects from Hwang et al. (2021) Using adopted individuals to partition maternal genetic effects into prenatal and postnatal effects on offspring phenotypes

\# Please contact Liang-Dar (Daniel) Hwang <> or David Evans <> for questions or if you want to fit a more complicated model, e.g., where biological and adoptive parents are related.

\# Input data: variance-covariance matrices between offspring phenotype and their own and their relatives' genotypes (or polygenic risk scores) from 7 different family structures.

#Fictitious data used in this example!

\# Required R package: "OpenMx"

\# Clear global environmental variables

rm(list=ls())

\# Load the R package "OpenMx"

library(OpenMx)

\# Read in a 4x4 variance-covariance matrix for G1: Biological Parent-offspring trios

\# The diagonal contains the variances offspring genotype (Zo), maternal genotype (Zm), paternal genotype (Zp), and offspring phenotype (Y), and the off-diagonal contains their covariances.

group_1 <- matrix(c(1.00, 0.00, 0.50, 0.25, 0.00, 1.00, 0.50, 0.15, 0.50, 0.50, 1.00, 0.25, 0.25, 0.15, 0.25, 1.00), nrow=4, byrow=TRUE) # This is where to read in the variance-covariance matrix

rownames(group_1)<- c("Zm", "Zp", "Zo", "Y")

colnames(group_1)<- c("Zm", "Zp", "Zo", "Y")

N_G1 <- 1000 # This is the number of biological parent-offspring trios used to derive the variance-covariance matrix

\# Read in a 3x3 variance-covariance matrix for G2: Mother-offspring pairs

group_2 <- matrix(c(1.00, 0.50, 0.25, 0.50, 1.00, 0.25, 0.25, 0.25, 1.00), nrow=3, byrow=TRUE) # This is where to read in the variance-covariance matrix

rownames(group_2)<- c("Zm", "Zo", "Y")

colnames(group_2)<- c("Zm", "Zo", "Y")

N_G2 <- 4000 # This is the number of biological Mother-offspring pairs used to derive the variance-covariance matrix

\# Read in a 3x3 variance-covariance matrix for G3: Father-offspring pairs

group_3 <- matrix(c(1.00, 0.50, 0.15, 0.50, 1.00, 0.25, 0.15, 0.25, 1.00), nrow=3, byrow=TRUE) # This is where to read in the variance-covariance matrix

rownames(group_3)<- c("Zp", "Zo", "Y")

colnames(group_3)<- c("Zp", "Zo", "Y")

N_G3 <- 1800 # This is the number of Father-offspring pairs used to derive the variance-covariance matrix

\# Read in a 2x2 variance-covariance matrix for G4: Singletons with Biological Parents

group_4 <- matrix(c(1.00, 0.25, 0.25, 1.00), nrow=2, byrow=TRUE) # This is where to read in the variance-covariance matrix

rownames(group_4)<- c("Zo", "Y")

colnames(group_4)<- c("Zo", "Y")

N_G4 <- 300000 # This is the number of Singletons with Biological Parents used to derive the variance-covariance matrix

\# Read in a 2x2 variance-covariance matrix for G5: Singletons with Adoptive Parents

\# Zo is adopted individual's genotype and Y is their phenotype

group_5 <- matrix(c(1.00, 0.15, 0.15, 0.96), nrow=2, byrow=TRUE) # This is where to read in the variance-covariance matrix

rownames(group_5)<- c("Zo", "Y")

colnames(group_5)<- c("Zo", "Y")

N_G5 <- 6000 # This is the number of Singletons with Adoptive Parents used to derive the variance-covariance matrix

\# Read in a 3x3 variance-covariance matrix for G6: Adoptive Mother - Adopted Child Pairs

\# Zo is adopted child's genotype. Zmf is adoptive mother's genotype. Y is adopted child's phenotype.

group_6 <- matrix(c(1.00, 0.0, 0.15, 0.00, 1.0, 0.10, 0.15, 0.1, 0.96), nrow=3, byrow=TRUE) # This is where to read in the variance-covariance matrix

rownames(group_6)<- c("Zo", "Zmf", "Y")

colnames(group_6)<- c("Zo", "Zmf", "Y")

N_G6 <- 0 # This is the number of Adoptive Mother - Adopted Child Pairs used to derive the variance-covariance matrix

\# Read in a 3x3 variance-covariance matrix for G7: Biological Mother - Adopted Child Pairs

\# Zo is adopted child's genotype. Zmb is biological mother's genotype. Y is adopted child's phenotype.

group_7 <- matrix(c(1.00, 0.50, 0.15, 0.50, 1.00, 0.15, 0.15, 0.15, 0.96), nrow=3, byrow=TRUE) # This is where to read in the variance-covariance matrix

rownames(group_7)<- c("Zo", "Zmb", "Y")

colnames(group_7)<- c("Zo", "Zmb", "Y")

N_G7 <- 50 # This is the number of Biological Mother - Adopted Child Pairs used to derive the variance-covariance matrix

\# Model parameters

V <- mxMatrix(type="Full", nrow=1, ncol=1, free=TRUE, values=1, label="v", name="V") #Variance of a SNP or a polygenic risk score

B_OY <-	mxMatrix(type="Full", nrow=1, ncol=1, free=TRUE, values=0, label="b_oy", name="B_OY") #Offspring genetic effect

B_MY <-	mxMatrix(type="Full", nrow=1, ncol=1, free=TRUE, values=0, label="b_my", name="B_MY") #Post-natal maternal genetic effect

B_PY <-	mxMatrix(type="Full", nrow=1, ncol=1, free=TRUE, values=0, label="b_py", name="B_PY") #Paternal genetic effect

G_MY <-	mxMatrix(type="Full", nrow=1, ncol=1, free=TRUE, values=0, label="g_my", name="G_MY") #Pre-natal maternal genetic effect

E1 <-	mxMatrix(type="Full", nrow=1, ncol=1, free=TRUE, values=0.9, label="e1", name="E1") #Phenotypic error variance in biological families

E2 <-	mxMatrix(type="Full", nrow=1, ncol=1, free=TRUE, values=0.9, label="e2", name="E2") #Phenotypic error variance in adopted families

R <- mxMatrix(type="Full", nrow=1, ncol=1, free=TRUE, values=0, label="rho", name="R") #Covariance between maternal and paternal genotypes

\# Define elements of expected covariance matrix in terms of parameters

\# G1 Parent offspring trios

c11_G1 <- mxAlgebra(expression=V, name="C11_G1")

c12_G1 <- mxAlgebra(expression=R, name="C12_G1")

c13_G1 <- mxAlgebra(expression=0.5*V+0.5*R, name="C13_G1")

c14_G1 <- mxAlgebra(expression=(G_MY + B_MY)*V + 0.5*B_OY*V + B_PY*R + 0.5*B_OY*R, name="C14_G1")

c21_G1 <- mxAlgebra(expression=R, name="C21_G1")

c22_G1 <- mxAlgebra(expression=V, name="C22_G1")

c23_G1 <- mxAlgebra(expression=0.5*V+0.5*R, name="C23_G1")

c24_G1 <- mxAlgebra(expression=B_PY*V + 0.5*B_OY*V + (G_MY+B_MY)*R + 0.5*B_OY*R, name="C24_G1")

c31_G1 <- mxAlgebra(expression=0.5*V+0.5*R, name="C31_G1")

c32_G1 <- mxAlgebra(expression=0.5*V+0.5*R, name="C32_G1")

c33_G1 <- mxAlgebra(expression=V+0.5*R, name="C33_G1")

c34_G1 <- mxAlgebra(expression=B_OY*(V+0.5*R) + 0.5*V*(G_MY + B_MY) + 0.5*V*B_PY + 0.5*(G_MY + B_MY)*R + 0.5*B_PY*R, name="C34_G1")

c41_G1 <- mxAlgebra(expression=(G_MY + B_MY)*V + 0.5*B_OY*V + B_PY*R + 0.5*B_OY*R, name="C41_G1")

c42_G1 <- mxAlgebra(expression=B_PY*V + 0.5*B_OY*V + (G_MY+B_MY)*R + 0.5*B_OY*R, name="C42_G1")

c43_G1 <- mxAlgebra(expression=B_OY*(V+0.5*R) + 0.5*V*(G_MY + B_MY) + 0.5*V*B_PY + 0.5*(G_MY+B_MY)*R + 0.5*B_PY*R, name="C43_G1")

c44_G1 <- mxAlgebra(expression=B_PY^2*V + B_OY^2*(V+0.5*R) + (G_MY + B_MY)^2*V + B_OY*V*(G_MY + B_MY) + B_OY*V*B_PY + E1 + 2*B_PY*R*(G_MY+B_MY) + B_PY*B_OY*R + B_OY*(G_MY+B_MY)*R, name="C44_G1")

\# G2 Mother offspring pairs

c11_G2 <- mxAlgebra(expression=V, name="C11_G2")

c12_G2 <- mxAlgebra(expression=0.5*V+0.5*R, name="C12_G2")

c13_G2 <- mxAlgebra(expression=(G_MY + B_MY)*V + 0.5*B_OY*V + B_PY*R + 0.5*B_OY*R, name="C13_G2")

c21_G2 <- mxAlgebra(expression=0.5*V+0.5*R, name="C21_G2")

c22_G2 <- mxAlgebra(expression=V+0.5*R, name="C22_G2")

c23_G2 <- mxAlgebra(expression=B_OY*(V+0.5*R) + 0.5*V*(G_MY + B_MY) + 0.5*V*B_PY + 0.5*(G_MY + B_MY)*R + 0.5*B_PY*R, name="C23_G2")

c31_G2 <- mxAlgebra(expression=(G_MY + B_MY)*V + 0.5*B_OY*V + B_PY*R + 0.5*B_OY*R, name="C31_G2")

c32_G2 <- mxAlgebra(expression=B_OY*(V+0.5*R) + 0.5*V*(G_MY + B_MY) + 0.5*V*B_PY + 0.5*(G_MY+B_MY)*R + 0.5*B_PY*R, name="C32_G2")

c33_G2 <- mxAlgebra(expression=B_PY^2*V + B_OY^2*(V+0.5*R) + (G_MY + B_MY)^2*V + B_OY*V*(G_MY + B_MY) + B_OY*V*B_PY + E1 + 2*B_PY*R*(G_MY+B_MY) + B_PY*B_OY*R + B_OY*(G_MY+B_MY)*R, name="C33_G2")

\# G3 Father offspring pairs

c11_G3 <- mxAlgebra(expression=V, name="C11_G3")

c12_G3 <- mxAlgebra(expression=0.5*V+0.5*R, name="C12_G3")

c13_G3 <- mxAlgebra(expression=B_PY*V + 0.5*B_OY*V + (G_MY+B_MY)*R + 0.5*B_OY*R, name="C13_G3")

c21_G3 <- mxAlgebra(expression=0.5*V+0.5*R, name="C21_G3")

c22_G3 <- mxAlgebra(expression=V+0.5*R, name="C22_G3")

c23_G3 <- mxAlgebra(expression=B_OY*(V+0.5*R) + 0.5*V*(G_MY + B_MY) + 0.5*V*B_PY + 0.5*(G_MY + B_MY)*R + 0.5*B_PY*R, name="C23_G3")

c31_G3 <- mxAlgebra(expression=B_PY*V + 0.5*B_OY*V + (G_MY+B_MY)*R + 0.5*B_OY*R, name="C31_G3")

c32_G3 <- mxAlgebra(expression=B_OY*(V+0.5*R) + 0.5*V*(G_MY + B_MY) + 0.5*V*B_PY + 0.5*(G_MY+B_MY)*R + 0.5*B_PY*R, name="C32_G3")

c33_G3 <- mxAlgebra(expression=B_PY^2*V + B_OY^2*(V+0.5*R) + (G_MY + B_MY)^2*V + B_OY*V*(G_MY + B_MY) + B_OY*V*B_PY + E1 + 2*B_PY*R*(G_MY+B_MY) + B_PY*B_OY*R + B_OY*(G_MY+B_MY)*R, name="C33_G3")

\# G4 Singletons (biological)

c11_G4 <- mxAlgebra(expression=V+0.5*R, name="C11_G4")

c12_G4 <- mxAlgebra(expression=B_OY*(V+0.5*R) + 0.5*V*(G_MY + B_MY) + 0.5*V*B_PY + 0.5*(G_MY + B_MY)*R + 0.5*B_PY*R, name="C12_G4")

c21_G4 <- mxAlgebra(expression=B_OY*(V+0.5*R) + 0.5*V*(G_MY + B_MY) + 0.5*V*B_PY + 0.5*(G_MY+B_MY)*R + 0.5*B_PY*R, name="C21_G4")

c22_G4 <- mxAlgebra(expression=B_PY^2*V + B_OY^2*(V+0.5*R) + (G_MY + B_MY)^2*V + B_OY*V*(G_MY + B_MY) + B_OY*V*B_PY + E1 + 2*B_PY*R*(G_MY+B_MY) + B_PY*B_OY*R + B_OY*(G_MY+B_MY)*R, name="C22_G4")

\# G5 Singletons (adopted)

c11_G5 <- mxAlgebra(expression=V+0.5*R, name="C11_G5")

c12_G5 <- mxAlgebra(expression=B_OY*(V+0.5*R) + 0.5*V*G_MY + 0.5*G_MY*R, name="C12_G5")

c21_G5 <- mxAlgebra(expression=B_OY*(V+0.5*R) + 0.5*V*G_MY + 0.5*G_MY*R, name="C21_G5")

c22_G5 <- mxAlgebra(expression=B_OY^2*(V+0.5*R) + G_MY^2*V + B_MY^2*V + B_PY^2*V + B_OY*V*G_MY + B_OY*G_MY*R + E2 + 2*B_MY*B_PY*R, name="C22_G5")

\# G6 Foster Mother - Adopted Child Pairs

c11_G6 <- mxAlgebra(expression=V+0.5*R, name="C11_G6")

c12_G6 <- mxAlgebra(expression=0, name="C12_G6")

c13_G6 <- mxAlgebra(expression=B_OY*(V+0.5*R) + 0.5*V*G_MY + 0.5*G_MY*R, name="C13_G6")

c21_G6 <- mxAlgebra(expression=0, name="C21_G6")

c22_G6 <- mxAlgebra(expression=V, name="C22_G6")

c23_G6 <- mxAlgebra(expression=B_MY*V, name="C23_G6")

c31_G6 <- mxAlgebra(expression=B_OY*(V+0.5*R) + 0.5*V*G_MY + 0.5*G_MY*R, name="C31_G6")

c32_G6 <- mxAlgebra(expression=B_MY*V, name="C32_G6")

c33_G6 <- mxAlgebra(expression=B_OY^2*(V+0.5*R) + G_MY^2*V + B_MY^2*V + B_PY^2*V + B_OY*V*G_MY + B_OY*G_MY*R + E2 + 2*B_MY*B_PY*R, name="C33_G6")

\# G7 Biological Mother - Adopted Child Pairs

c11_G7 <- mxAlgebra(expression=V+0.5*R, name="C11_G7")

c12_G7 <- mxAlgebra(expression=0.5*V+0.5*R, name="C12_G7")

c13_G7 <- mxAlgebra(expression=B_OY*(V+0.5*R) + 0.5*V*G_MY + 0.5*G_MY*R, name="C13_G7")

c21_G7 <- mxAlgebra(expression=0.5*V+0.5*R, name="C21_G7")

c22_G7 <- mxAlgebra(expression=V, name="C22_G7")

c23_G7 <- mxAlgebra(expression=G_MY*V + 0.5*B_OY*V + 0.5*B_OY*R, name="C23_G7")

c31_G7 <- mxAlgebra(expression=B_OY*(V+0.5*R) + 0.5*V*G_MY + 0.5*G_MY*R, name="C31_G7")

c32_G7 <- mxAlgebra(expression=G_MY*V + 0.5*B_OY*V + 0.5*B_OY*R, name="C32_G7")

c33_G7 <- mxAlgebra(expression=B_OY^2*(V+0.5*R) + G_MY^2*V + B_MY^2*V + B_PY^2*V + B_OY*V*G_MY + B_OY*G_MY*R + E2 + 2*B_MY*B_PY*R, name="C33_G7")

\# Expected covariance matrices

expCov_G1 <- mxAlgebra( expression= rbind( cbind(C11_G1, C12_G1, C13_G1, C14_G1), cbind(C21_G1, C22_G1, C23_G1, C24_G1), cbind(C31_G1, C32_G1, C33_G1, C34_G1), cbind(C41_G1, C42_G1, C43_G1, C44_G1)), name="expCov_G1")

expCov_G2 <- mxAlgebra( expression= rbind( cbind(C11_G2, C12_G2, C13_G2), cbind(C21_G2, C22_G2, C23_G2), cbind(C31_G2, C32_G2, C33_G2)), name="expCov_G2")

expCov_G3 <- mxAlgebra( expression= rbind( cbind(C11_G3, C12_G3, C13_G3), cbind(C21_G3, C22_G3, C23_G3), cbind(C31_G3, C32_G3, C33_G3)), name="expCov_G3")

expCov_G4 <- mxAlgebra( expression= rbind( cbind(C11_G4, C12_G4), cbind(C21_G4, C22_G4)), name="expCov_G4")

expCov_G5 <- mxAlgebra( expression= rbind( cbind(C11_G5, C12_G5), cbind(C21_G5, C22_G5)), name="expCov_G5")

expCov_G6 <- mxAlgebra( expression= rbind( cbind(C11_G6, C12_G6, C13_G6), cbind(C21_G6, C22_G6, C23_G6), cbind(C31_G6, C32_G6, C33_G6)), name="expCov_G6")

expCov_G7 <- mxAlgebra( expression= rbind( cbind(C11_G7, C12_G7, C13_G7), cbind(C21_G7, C22_G7, C23_G7), cbind(C31_G7, C32_G7, C33_G7)), name="expCov_G7")

\# Create Data Objects for Multiple Groups

dataG1	<- mxData(observed=(group_1), type="cov", numObs=N_G1)

dataG2	<- mxData(observed=(group_2), type="cov", numObs=N_G2)

dataG3	<- mxData(observed=(group_3), type="cov", numObs=N_G3)

dataG4	<- mxData(observed=(group_4), type="cov", numObs=N_G4)

dataG5	<- mxData(observed=(group_5), type="cov", numObs=N_G5)

dataG6	<- mxData(observed=(group_6), type="cov", numObs=N_G6)

dataG7	<- mxData(observed=(group_7), type="cov", numObs=N_G7)

\# Create Expectation Objects for Multiple Groups

expG1	<- mxExpectationNormal( covariance="expCov_G1", dimnames=c("Zm","Zp","Zo","Y") )

expG2	<- mxExpectationNormal( covariance="expCov_G2", dimnames=c("Zm","Zo", "Y") )

expG3	<- mxExpectationNormal( covariance="expCov_G3", dimnames=c("Zp","Zo","Y") )

expG4	<- mxExpectationNormal( covariance="expCov_G4", dimnames=c("Zo","Y") )

expG5	<- mxExpectationNormal( covariance="expCov_G5", dimnames=c("Zo","Y") )

expG6	<- mxExpectationNormal( covariance="expCov_G6", dimnames=c("Zo","Zmf","Y") )

expG7	<- mxExpectationNormal( covariance="expCov_G7", dimnames=c("Zo","Zmb","Y") )

funML	<- mxFitFunctionML()

\# Create Model Objects for Multiple Groups

pars	<- list( V, B_OY, B_MY, B_PY, G_MY, E1, E2, R)

modelG1	<- mxModel( V, B_OY, B_MY, B_PY, G_MY, E1, R, c11_G1, c12_G1, c13_G1, c14_G1, c21_G1, c22_G1, c23_G1, c24_G1, c31_G1, c32_G1, c33_G1, c34_G1, c41_G1, c42_G1, c43_G1, c44_G1, expG1, expCov_G1, dataG1, funML, name="G1" )

modelG2	<- mxModel( V, B_OY, B_MY, B_PY, G_MY, E1, R, c11_G2, c12_G2, c13_G2, c21_G2, c22_G2, c23_G2, c31_G2, c32_G2, c33_G2, expG2, expCov_G2, dataG2, funML, name="G2" )

modelG3	<- mxModel( V, B_OY, B_MY, B_PY, G_MY, E1, R, c11_G3, c12_G3, c13_G3, c21_G3, c22_G3, c23_G3, c31_G3, c32_G3, c33_G3, expG3, expCov_G3, dataG3, funML, name="G3" )

modelG4	<- mxModel( V, B_OY, B_MY, B_PY, G_MY, E1, R, c11_G4, c12_G4, c21_G4, c22_G4, expG4, expCov_G4, dataG4, funML, name="G4" )

modelG5	<- mxModel( V, B_OY, B_MY, B_PY, G_MY, E2, R, c11_G5, c12_G5, c21_G5, c22_G5, expG5, expCov_G5, dataG5, funML, name="G5" )

modelG6	<- mxModel( V, B_OY, B_MY, B_PY, G_MY, E2, R, c11_G6, c12_G6, c13_G6, c21_G6, c22_G6, c23_G6, c31_G6, c32_G6, c33_G6, expG6, expCov_G6, dataG6, funML, name="G6" )

modelG7	<- mxModel( V, B_OY, B_MY, B_PY, G_MY, E2, R, c11_G7, c12_G7, c13_G7, c21_G7, c22_G7, c23_G7, c31_G7, c32_G7, c33_G7, expG7, expCov_G7, dataG7, funML, name="G7" )

multi	<- mxFitFunctionMultigroup( c("G1","G2","G3","G4","G5","G6","G7") )

modelFull <- mxModel( "Full_Model", pars, modelG1, modelG2, modelG3, modelG4, modelG5, modelG6, modelG7, multi)

\# RUN MODEL

fitFull	<- mxRun( modelFull, intervals=F )

sumFull	<- summary( fitFull )

sumFull

\# v, variance of the SNP (polygenic risk score)

\# B_OY, offspring genetic effect

\# b_my, postnatal maternal genetic effect

\# b_py, paternal genetic effect

\# g_my, prenatal maternal genetic effect

\# e1, error variance of biological children's phenotype

\# e2, error variance of adopted children's phenotype

\# rho, covariance between maternal and paternal genotypes
