## Supplementary Table 1 for "Using adopted individuals to partition maternal genetic effects into prenatal and postnatal effects on offspring phenotypes"

**Table S1.** SNPs used to construct a weighted polygenic score of maternal birthweight. SNPs were selected fro

| SNP | Chromosome | Position hg19 (bp) | Effect Allele | Other Allele | Beta | Error |
|---|---|---|---|---|---|---|
| rs560887 | 2 | 169763148 | C | T | 0.038 | 0.005 |
| rs9855896 | 3 | 14287150 | G | A | 0.033 | 0.006 |
| rs11708067 | 3 | 123065778 | A | G | 0.029 | 0.005 |
| rs4679760 | 3 | 155855418 | G | C | 0.038 | 0.005 |
| rs2946179 | 5 | 157886627 | C | T | 0.045 | 0.005 |
| rs9379084 | 6 | 7231843 | G | A | 0.040 | 0.007 |
| rs45446698 | 7 | 99332948 | G | T | 0.077 | 0.012 |
| rs6995390 | 8 | 77611012 | T | A | 0.039 | 0.006 |
| rs72760655 | 9 | 116916214 | C | A | 0.031 | 0.005 |
| rs10509669 | 10 | 95969913 | A | T | 0.039 | 0.005 |
| rs3740360 | 10 | 96025491 | C | A | 0.044 | 0.007 |
| rs2168101 | 11 | 8255408 | C | A | 0.039 | 0.005 |
| rs10830963 | 11 | 92708710 | G | C | 0.046 | 0.005 |
| rs180438 | 12 | 47187260 | G | A | 0.039 | 0.006 |
| rs3184504 | 12 | 111884608 | C | T | 0.034 | 0.005 |
| rs3784789 | 15 | 75082552 | G | C | 0.030 | 0.005 |
| rs12909648 | 15 | 86224570 | G | A | 0.027 | 0.005 |
| rs34717629 | 17 | 17610404 | A | G | 0.032 | 0.006 |
| rs2967677 | 19 | 8789721 | C | T | 0.050 | 0.007 |
| rs6608539 | 23 | 115132834 | A | G | 0.027 | 0.004 |

*rs2967676 was not included because it is in complete linkage disequilibrium with rs2967677.

m Supplemetary Table 5 in Warrington et al. (2019) if they have "SEM-adjusted Maternal Effects" wit

| **P-value** |
|---|
| 5.4E-14 |
| 2.6E-09 |
| 3.8E-08 |
| 6.5E-15 |
| 3.7E-17 |
| 4.7E-08 |
| 1.1E-10 |
| 3.9E-10 |
| 7.6E-10 |
| 3.6E-13 |
| 1.6E-09 |
| 5.9E-14 |
| 4.6E-19 |
| 4.1E-11 |
| 1.8E-13 |
| 1.2E-09 |
| 3.4E-09 |
| 4.4E-08 |
| 3.8E-14 |
| 1.1E-09 |

h $p<5.0 \times 10^{-8}$.
