## Supplementary Table 2 for "Using adopted individuals to partition maternal genetic effects into prenatal and postnatal effects on offspring phenotypes"

**Table S2.** SNPs used to construct unweighted polygenic scores of educational attainment.

| Lee et al. 2018 | | | | | | | | |
|---|---|---|---|---|---|---|---|---|
| **SNP** | **Chromosome** | **Position hg19 (bp)** | **Effect Allele** | **Beta** | **Standard Error** | **P-value** | **Note** | **SNP** |
| rs780569 | 1 | 4569436 | A | -0.0085 | 0.0016 | 4.98E-08 | | rs301800 |
| rs3439405 | 1 | 6853091 | A | -0.0117 | 0.002 | 2.30E-09 | | rs1121086 |
| rs1112117 | 1 | 8445946 | A | 0.0151 | 0.0018 | 1.44E-16 | | rs3430537 |
| rs4846010 | 1 | 11531059 | A | 0.0099 | 0.0018 | 3.35E-08 | | rs2568955 |
| rs7811607 | 1 | 18434125 | C | 0.0086 | 0.0016 | 3.33E-08 | | rs1008078 |
| rs1079961 | 1 | 20541370 | A | -0.0089 | 0.0016 | 4.11E-08 | | rs1158885 |
| rs1112359 | 1 | 20871750 | T | 0.0133 | 0.0024 | 2.21E-08 | | rs1777827 |
| rs1212792 | 1 | 28708529 | T | 0.0108 | 0.0018 | 1.69E-09 | | rs2992632 |
| rs590013 | 1 | 29155738 | T | 0.01 | 0.0015 | 2.47E-11 | | rs7607633 |
| rs1079888 | 1 | 32199099 | T | -0.014 | 0.0019 | 5.15E-14 | | rs1168926 |
| rs1202801 | 1 | 41764471 | T | 0.0164 | 0.0017 | 9.78E-23 | | rs1606974 |
| rs1120989 | 1 | 41798668 | A | 0.0119 | 0.0021 | 8.03E-09 | | rs1169017 |
| rs2364544 | 1 | 41833162 | A | -0.0105 | 0.0014 | 2.91E-13 | | rs2457660 |
| rs2992037 | 1 | 43949810 | A | 0.0149 | 0.0015 | 4.18E-23 | | rs1145988 |
| rs5885955 | 1 | 44010456 | T | 0.0187 | 0.0028 | 2.82E-11 | | rs1049609 |
| rs1207663 | 1 | 44026656 | C | 0.0207 | 0.0017 | 3.33E-34 | | rs1340290 |
| rs7267309 | 1 | 44087925 | A | 0.0257 | 0.0042 | 8.90E-10 | | rs4851251 |
| rs7720169 | 1 | 44347087 | A | 0.0156 | 0.0022 | 1.28E-12 | | rs1298766 |
| rs803619 | 1 | 44465419 | T | -0.0173 | 0.0023 | 2.19E-14 | | rs1782424 |
| rs1738050 | 1 | 44707295 | C | -0.0093 | 0.0014 | 1.42E-10 | | rs1684558 |
| rs1121112 | 1 | 45957609 | A | 0.0104 | 0.0017 | 7.62E-10 | | rs4500960 |
| rs6696068 | 1 | 53740797 | T | -0.0088 | 0.0014 | 9.50E-10 | | rs6739979 |
| rs548897 | 1 | 57718030 | A | 0.0081 | 0.0014 | 1.10E-08 | | rs2245901 |
| rs852771 | 1 | 58288317 | T | 0.0092 | 0.0015 | 1.84E-09 | | rs5583072 |
| rs2764684 | 1 | 58537919 | T | 0.0143 | 0.0019 | 9.99E-15 | | rs3576124 |
| rs2989476 | 1 | 61059259 | C | 0.0078 | 0.0014 | 3.73E-08 | | rs6225953 |
| rs4915735 | 1 | 61808444 | A | 0.0116 | 0.002 | 9.56E-09 | | rs1487347 |
| rs5567558 | 1 | 66224881 | T | -0.0103 | 0.0018 | 5.92E-09 | | rs1171205 |
| rs1392816 | 1 | 66481188 | T | 0.0082 | 0.0014 | 1.61E-08 | | rs1126343 |
| rs7266746 | 1 | 66536012 | T | 0.0197 | 0.0031 | 2.44E-10 | | rs6226392 |
| rs1744088 | 1 | 67208622 | A | 0.016 | 0.0029 | 2.42E-08 | | rs6799130 |
| rs7552964 | 1 | 69331299 | A | 0.0148 | 0.0023 | 1.78E-10 | | rs1264680 |
| rs1078928 | 1 | 69788482 | T | 0.0091 | 0.0016 | 2.07E-08 | | rs2610986 |
| rs1024268 | 1 | 70117404 | T | -0.0084 | 0.0014 | 3.40E-09 | | rs3407209 |
| rs663251 | 1 | 70566021 | T | 0.0081 | 0.0015 | 4.02E-08 | | rs3101246 |
| rs481940 | 1 | 71442457 | T | 0.01 | 0.0016 | 4.94E-10 | | rs4863692 |
| rs7267717 | 1 | 72120034 | A | 0.01 | 0.0014 | 3.03E-12 | | rs4493682 |
| rs3412291 | 1 | 72509742 | C | 0.0084 | 0.0014 | 4.16E-09 | | rs2964197 |
| rs3430537 | 1 | 72733610 | A | 0.0314 | 0.0024 | 1.10E-39 | | rs6116018 |
| rs2568955 | 1 | 72762169 | T | -0.0165 | 0.0016 | 2.51E-24 | | rs324886 |
| rs1445591 | 1 | 72960011 | A | 0.0085 | 0.0015 | 4.02E-08 | | rs1006178 |
| rs1202822 | 1 | 72986136 | T | 0.0099 | 0.0016 | 1.67E-09 | | rs2431108 |
| rs1121022 | 1 | 73859290 | T | -0.0105 | 0.0019 | 2.20E-08 | | rs1402025 |

| | | | | | | | |
|---|---|---|---|---|---|---|---|
| rs7409167 | 1 | 73978572 | A | -0.0153 | 0.0022 | 6.46E-12 | rs6237983 |
| rs1121040 | 1 | 74490459 | A | 0.012 | 0.0014 | 1.63E-17 | rs5623133 |
| rs1569092 | 1 | 74858724 | A | 0.016 | 0.002 | 2.12E-16 | rs9320913 |
| rs2848208 | 1 | 75226252 | A | 0.0086 | 0.0015 | 4.51E-09 | rs7767938 |
| rs7999473 | 1 | 75562222 | T | 0.024 | 0.0035 | 5.88E-12 | rs2615691 |
| rs1078265 | 1 | 77934335 | T | -0.0082 | 0.0014 | 6.37E-09 | rs1253145 |
| rs1335482 | 1 | 78603463 | T | 0.0096 | 0.0014 | 1.01E-11 | rs1267193 |
| rs6428587 | 1 | 90825188 | T | 0.0084 | 0.0015 | 8.59E-09 | rs1135204 |
| rs4480339 | 1 | 91096799 | A | -0.0097 | 0.0015 | 1.53E-10 | rs1716717 |
| rs1713112 | 1 | 91126097 | A | -0.0243 | 0.0039 | 6.29E-10 | rs1176823 |
| rs5617499 | 1 | 91145181 | A | -0.0143 | 0.002 | 2.81E-12 | rs1268229 |
| rs1008078 | 1 | 91189731 | T | -0.0182 | 0.0014 | 4.50E-37 | rs1871109 |
| rs7526112 | 1 | 93747683 | T | 0.0124 | 0.0015 | 2.07E-17 | rs1329443 |
| rs236318 | 1 | 94050221 | C | -0.0102 | 0.0017 | 7.40E-10 | rs895606 |
| rs6690195 | 1 | 96221653 | T | -0.0113 | 0.0014 | 7.89E-16 | rs7854982 |
| rs1378583 | 1 | 96454977 | A | 0.0193 | 0.0029 | 1.55E-11 | rs1119119 |
| rs635754 | 1 | 96484940 | A | 0.0119 | 0.0014 | 6.36E-17 | rs1277237 |
| rs1879519 | 1 | 98412713 | A | -0.0315 | 0.0058 | 4.87E-08 | rs7945718 |
| rs1087512 | 1 | 98417446 | C | 0.0174 | 0.0019 | 4.46E-20 | rs7955289 |
| rs1843815 | 1 | 98589715 | A | 0.0077 | 0.0014 | 4.40E-08 | rs2456973 |
| rs1275068 | 1 | 98757620 | C | 0.0116 | 0.0016 | 6.64E-13 | rs7131944 |
| rs9435340 | 1 | 107593201 | A | 0.0086 | 0.0015 | 5.99E-09 | rs572016 |
| rs575113 | 1 | 110046373 | A | 0.0133 | 0.0015 | 7.52E-18 | rs7306755 |
| rs3768480 | 1 | 110530995 | C | 0.0101 | 0.0014 | 8.40E-13 | rs9537821 |
| rs6689641 | 1 | 110720400 | A | 0.0078 | 0.0014 | 3.29E-08 | rs1043209 |
| rs2761438 | 1 | 110752139 | A | 0.0098 | 0.0015 | 1.59E-11 | rs8005528 |
| rs7657742 | 1 | 110761099 | C | 0.0203 | 0.0024 | 7.78E-18 | rs1711997 |
| rs4839155 | 1 | 112161489 | T | 0.0115 | 0.0017 | 4.66E-12 | rs1928185 |
| rs1538389 | 1 | 112327144 | T | -0.0109 | 0.0018 | 1.27E-09 | rs1296929 |
| rs3790609 | 1 | 113056990 | T | 0.0107 | 0.0019 | 8.35E-09 | rs2837992 |
| rs1274655 | 1 | 114374966 | C | 0.0248 | 0.0041 | 1.44E-09 | rs165633 |
| rs2458370 | 1 | 117874944 | T | -0.0086 | 0.0015 | 4.23E-09 | |
| rs6174895 | 1 | 156038058 | A | -0.0273 | 0.0048 | 1.31E-08 | |
| rs3026996 | 1 | 159167290 | A | 0.0138 | 0.0016 | 5.42E-17 | |
| rs2254681 | 1 | 159248048 | A | 0.0101 | 0.0016 | 4.25E-10 | |
| rs1126519 | 1 | 159400359 | T | 0.0083 | 0.0015 | 2.14E-08 | |
| rs3472038 | 1 | 171455322 | T | -0.018 | 0.0024 | 1.90E-13 | |
| rs3817923 | 1 | 171809211 | A | -0.0121 | 0.0022 | 4.11E-08 | |
| rs6672986 | 1 | 174076864 | C | 0.0094 | 0.0014 | 6.42E-11 | |
| rs4652135 | 1 | 175837673 | A | 0.0103 | 0.0016 | 4.35E-11 | |
| rs6697584 | 1 | 181509718 | T | -0.0126 | 0.0017 | 1.43E-13 | |
| rs7550087 | 1 | 181596751 | T | 0.0207 | 0.0033 | 2.48E-10 | |
| rs3766979 | 1 | 181636598 | A | 0.0085 | 0.0015 | 9.94E-09 | |
| rs1212623 | 1 | 184698816 | A | 0.0095 | 0.0014 | 4.82E-11 | |
| rs1171040 | 1 | 190400984 | A | -0.0113 | 0.0017 | 5.07E-11 | |
| rs4658019 | 1 | 196149936 | T | 0.0084 | 0.0014 | 2.19E-09 | |
| rs1747817 | 1 | 197711492 | T | -0.0092 | 0.0016 | 2.29E-08 | |

| | | | | | | |
|---|---|---|---|---|---|---|
| rs1214507 | 1 | 197822977 | A | -0.0127 | 0.0017 | 2.56E-13 |
| rs3478070 | 1 | 197907554 | A | 0.0111 | 0.0017 | 2.05E-11 |
| rs1890132 | 1 | 199427321 | T | -0.0089 | 0.0015 | 6.56E-09 |
| rs8024 | 1 | 201845575 | A | -0.0115 | 0.0015 | 1.13E-14 |
| rs1158885 | 1 | 204587047 | A | 0.0199 | 0.0017 | 4.45E-31 |
| rs884108 | 1 | 204591237 | A | -0.0094 | 0.0017 | 2.39E-08 |
| rs1212329 | 1 | 210912835 | T | 0.0086 | 0.0015 | 4.48E-09 |
| rs3111251 | 1 | 211409844 | T | -0.009 | 0.0014 | 3.88E-10 |
| rs7836524 | 1 | 211737950 | T | 0.0181 | 0.0033 | 4.38E-08 |
| rs7164614 | 1 | 212369228 | T | 0.0104 | 0.0018 | 5.69E-09 |
| rs1256815 | 1 | 213151580 | T | -0.0091 | 0.0015 | 3.52E-09 |
| rs1391513 | 1 | 216693263 | T | -0.0095 | 0.0015 | 5.25E-10 |
| rs4846724 | 1 | 221967817 | A | 0.0098 | 0.0014 | 2.89E-12 |
| rs6695132 | 1 | 234734655 | T | 0.0097 | 0.0017 | 1.01E-08 |
| rs1329125 | 1 | 234740880 | T | -0.0097 | 0.0015 | 8.37E-11 |
| rs1240588 | 1 | 235597252 | T | -0.0098 | 0.0014 | 2.30E-12 |
| rs1145931 | 1 | 241876565 | A | -0.0122 | 0.0017 | 1.65E-12 |
| rs3897821 | 1 | 243420388 | A | 0.0154 | 0.0015 | 3.58E-25 |
| rs2994326 | 1 | 243651026 | T | -0.01 | 0.0018 | 3.38E-08 |
| rs1092705 | 1 | 243811321 | A | -0.0125 | 0.0022 | 2.13E-08 |
| rs622169 | 1 | 244437407 | T | 0.0086 | 0.0014 | 2.73E-09 |
| rs1261426 | 2 | 4123797 | A | 0.0083 | 0.0014 | 3.38E-09 |
| rs7603132 | 2 | 4951548 | A | 0.0137 | 0.0018 | 1.20E-14 |
| rs7590368 | 2 | 10961474 | T | -0.0146 | 0.0016 | 3.98E-20 |
| rs7607633 | 2 | 10977585 | T | 0.0206 | 0.0021 | 1.71E-23 |
| rs1734370 | 2 | 10978710 | A | 0.0099 | 0.0016 | 2.42E-10 |
| rs5582649 | 2 | 12836640 | T | -0.0127 | 0.0021 | 3.16E-09 |
| rs1991585 | 2 | 16647170 | T | -0.0101 | 0.0015 | 6.26E-11 |
| rs312945 | 2 | 21328825 | A | -0.0091 | 0.0015 | 2.34E-09 |
| rs6738860 | 2 | 22442099 | A | -0.0088 | 0.0014 | 4.65E-10 |
| rs4358081 | 2 | 29100642 | A | -0.0095 | 0.0014 | 1.28E-11 |
| rs1302960 | 2 | 29574875 | T | -0.0092 | 0.0014 | 1.33E-10 |
| rs6543810 | 2 | 34380023 | T | 0.0082 | 0.0015 | 2.11E-08 |
| rs7288111 | 2 | 44754020 | A | 0.0146 | 0.0027 | 3.59E-08 |
| rs1246804 | 2 | 44854981 | T | 0.0137 | 0.0014 | 3.12E-21 |
| rs7279239 | 2 | 44886144 | T | -0.0145 | 0.0022 | 1.09E-10 |
| rs163229 | 2 | 45157163 | C | -0.026 | 0.0047 | 4.13E-08 |
| rs6138783 | 2 | 48688635 | A | 0.0088 | 0.0015 | 1.11E-08 |
| rs6744040 | 2 | 49908151 | A | -0.008 | 0.0014 | 1.15E-08 |
| rs939400 | 2 | 50645890 | T | -0.0095 | 0.0015 | 7.52E-11 |
| rs1339886 | 2 | 50692940 | A | 0.0086 | 0.0015 | 5.89E-09 |
| rs1750293 | 2 | 50994255 | T | -0.0167 | 0.002 | 2.12E-17 |
| rs1757400 | 2 | 51230274 | A | -0.0129 | 0.0021 | 1.69E-09 |
| rs1262079 | 2 | 51287849 | A | 0.0121 | 0.0019 | 9.25E-11 |
| rs6214289 | 2 | 51539113 | A | 0.0108 | 0.0016 | 1.05E-11 |
| rs1301028 | 2 | 51824512 | T | 0.0198 | 0.0021 | 3.44E-21 |
| rs3948495 | 2 | 57383133 | T | -0.0089 | 0.0014 | 3.95E-10 |

| | | | | | | |
|---|---|---|---|---|---|---|
| rs1106090 | 2 | 58068741 | A | 0.0094 | 0.0014 | 1.00E-10 |
| rs6058953 | 2 | 58072605 | A | 0.0168 | 0.0028 | 3.03E-09 |
| rs6736025 | 2 | 58797634 | T | 0.0079 | 0.0014 | 2.77E-08 |
| rs1169388 | 2 | 60122046 | A | -0.0091 | 0.0014 | 1.22E-10 |
| rs5598678 | 2 | 60655551 | T | 0.0187 | 0.0026 | 2.27E-13 |
| rs1018985 | 2 | 60713235 | A | 0.0158 | 0.0014 | 4.69E-29 |
| rs2665668 | 2 | 60768978 | A | -0.0094 | 0.0015 | 1.25E-10 |
| rs356999 | 2 | 60811584 | A | -0.0101 | 0.0014 | 2.91E-12 |
| rs1163867 | 2 | 61018515 | C | 0.0249 | 0.0045 | 4.35E-08 |
| rs1049609 | 2 | 61482261 | A | -0.0131 | 0.0016 | 3.71E-17 |
| rs3549393 | 2 | 65571139 | C | 0.0124 | 0.0017 | 9.73E-14 |
| rs6731373 | 2 | 68503044 | A | -0.0115 | 0.0015 | 1.39E-14 |
| rs3436386 | 2 | 73490412 | A | 0.0087 | 0.0014 | 8.69E-10 |
| rs2916490 | 2 | 80192352 | A | -0.0092 | 0.0015 | 1.92E-09 |
| rs1128064 | 2 | 80421805 | C | -0.016 | 0.0025 | 1.72E-10 |
| rs1189442 | 2 | 80588091 | A | 0.0209 | 0.0033 | 2.14E-10 |
| rs1838692 | 2 | 81751340 | C | 0.0174 | 0.003 | 4.72E-09 |
| rs7291945 | 2 | 81999568 | T | -0.0132 | 0.0023 | 1.67E-08 |
| rs3575169 | 2 | 98242555 | T | -0.0246 | 0.004 | 8.04E-10 |
| rs1303434 | 2 | 98329197 | T | -0.0082 | 0.0014 | 9.29E-09 |
| rs6215535 | 2 | 98932980 | A | 0.0338 | 0.0053 | 1.69E-10 |
| rs6215577 | 2 | 99981222 | A | -0.0196 | 0.0032 | 6.88E-10 |
| rs6715321 | 2 | 100109001 | T | -0.0097 | 0.0014 | 7.62E-12 |
| rs7782640 | 2 | 100251389 | T | 0.0151 | 0.0027 | 2.05E-08 |
| rs6715849 | 2 | 100306378 | A | -0.0122 | 0.0014 | 6.87E-18 |
| rs1300991 | 2 | 100423158 | T | 0.0212 | 0.0037 | 1.36E-08 |
| rs7281911 | 2 | 100451357 | T | -0.0231 | 0.0024 | 8.29E-22 |
| rs7520341 | 2 | 100513153 | T | 0.0135 | 0.0019 | 7.68E-13 |
| rs4851263 | 2 | 100801867 | A | -0.0151 | 0.0023 | 7.86E-11 |
| rs1301864 | 2 | 100821545 | T | -0.0215 | 0.0014 | 1.05E-50 |
| rs7141387 | 2 | 100924822 | A | 0.0351 | 0.0036 | 2.66E-22 |
| rs1016628 | 2 | 101128095 | T | 0.0151 | 0.0019 | 3.79E-16 |
| rs3410669 | 2 | 101151830 | C | 0.0159 | 0.0019 | 1.62E-16 |
| rs2942884 | 2 | 101323578 | A | -0.009 | 0.0014 | 2.27E-10 |
| rs7770281 | 2 | 101328728 | T | 0.0187 | 0.0025 | 4.17E-14 |
| rs4850954 | 2 | 101573214 | T | 0.0077 | 0.0014 | 3.73E-08 |
| rs1020405 | 2 | 103775435 | T | -0.0101 | 0.0014 | 1.92E-12 |
| rs3474802 | 2 | 104060003 | A | 0.0185 | 0.0025 | 3.72E-13 |
| rs2570497 | 2 | 104441546 | T | -0.0121 | 0.0015 | 1.27E-16 |
| rs6215587 | 2 | 105969362 | T | -0.0151 | 0.0021 | 1.65E-12 |
| rs1271226 | 2 | 107564115 | T | -0.0107 | 0.0014 | 6.88E-14 |
| rs1340249 | 2 | 115826318 | A | -0.008 | 0.0014 | 1.33E-08 |
| rs7891815 | 2 | 117611525 | T | -0.008 | 0.0015 | 4.26E-08 |
| rs5899689 | 2 | 123735005 | A | -0.0094 | 0.0016 | 4.23E-09 |
| rs4848924 | 2 | 124991887 | A | -0.0117 | 0.0015 | 3.56E-14 |
| rs1301056 | 2 | 125873208 | A | -0.0091 | 0.0014 | 1.03E-10 |
| rs6743032 | 2 | 126024787 | A | -0.0165 | 0.0024 | 8.36E-12 |

| | | | | | | | |
|---|---|---|---|---|---|---|---|
| rs1302661 | 2 | 128153335 | A | -0.0084 | 0.0015 | 1.05E-08 | |
| rs1765063 | 2 | 139420392 | T | -0.0104 | 0.0019 | 2.19E-08 | |
| rs7297296 | 2 | 140454376 | A | 0.0083 | 0.0015 | 2.94E-08 | |
| rs1016900 | 2 | 140563368 | A | -0.008 | 0.0014 | 1.15E-08 | |
| rs7530881 | 2 | 140642378 | A | 0.0287 | 0.0047 | 9.39E-10 | |
| rs1464297 | 2 | 140653749 | T | -0.0109 | 0.0015 | 1.49E-13 | |
| rs7478792 | 2 | 141598681 | A | 0.0182 | 0.0028 | 8.97E-11 | |
| rs7597412 | 2 | 142069014 | C | 0.0088 | 0.0015 | 1.68E-09 | |
| rs7760976 | 2 | 142358950 | A | 0.0236 | 0.0028 | 7.71E-17 | |
| rs1018107 | 2 | 142862830 | A | 0.0102 | 0.0019 | 3.31E-08 | |
| rs7561705 | 2 | 143446472 | A | 0.0078 | 0.0014 | 2.71E-08 | |
| rs1320139 | 2 | 144158177 | C | -0.0149 | 0.0014 | 7.62E-26 | |
| rs1092819 | 2 | 144376033 | T | -0.0087 | 0.0014 | 7.76E-10 | |
| rs7396184 | 2 | 144478983 | A | -0.0114 | 0.0014 | 2.52E-15 | |
| rs3462479 | 2 | 144916804 | T | 0.0116 | 0.0021 | 1.94E-08 | |
| rs1427298 | 2 | 145214421 | T | 0.0086 | 0.0014 | 1.33E-09 | |
| rs7560871 | 2 | 145616899 | A | -0.0193 | 0.0027 | 5.24E-13 | |
| rs7575637 | 2 | 145796251 | A | 0.0107 | 0.0014 | 3.43E-14 | |
| rs1774234 | 2 | 148633936 | A | -0.0119 | 0.0018 | 1.59E-11 | |
| rs6729612 | 2 | 152780167 | T | 0.0092 | 0.0015 | 9.39E-10 | |
| rs1342267 | 2 | 155474355 | T | -0.011 | 0.0014 | 6.48E-15 | |
| rs6720515 | 2 | 156603535 | A | 0.0103 | 0.0017 | 7.95E-10 | |
| rs1554798 | 2 | 156835059 | A | -0.008 | 0.0014 | 1.22E-08 | |
| rs7290612 | 2 | 157076201 | T | 0.0185 | 0.003 | 6.44E-10 | |
| rs1368250 | 2 | 157485517 | T | 0.025 | 0.0045 | 3.00E-08 | |
| rs1342558 | 2 | 157487273 | C | -0.0089 | 0.0014 | 2.32E-10 | |
| rs4664983 | 2 | 159454438 | T | -0.0103 | 0.0018 | 4.11E-09 | |
| rs1339752 | 2 | 161235981 | C | -0.0097 | 0.0017 | 7.22E-09 | |
| rs6217735 | 2 | 161290337 | A | -0.0275 | 0.0038 | 6.87E-13 | |
| rs1019236 | 2 | 161380888 | A | -0.0078 | 0.0014 | 2.36E-08 | |
| rs6752228 | 2 | 161547697 | T | -0.008 | 0.0014 | 1.26E-08 | |
| rs7999716 | 2 | 161808690 | A | 0.0172 | 0.003 | 7.14E-09 | |
| rs1149529 | 2 | 161812352 | T | 0.0265 | 0.004 | 4.44E-11 | |
| rs6760887 | 2 | 161859760 | T | 0.0155 | 0.0022 | 1.17E-12 | |
| rs6219417 | 2 | 161866504 | A | 0.0111 | 0.0017 | 2.41E-10 | |
| rs1267062 | 2 | 162000266 | C | 0.0113 | 0.0018 | 7.40E-10 | |
| rs1167898 | 2 | 162101261 | A | -0.0166 | 0.0014 | 1.60E-31 | |
| rs4500960 | 2 | 162818621 | T | -0.0131 | 0.0014 | 1.04E-20 | |
| rs2098526 | 2 | 162845276 | A | -0.0241 | 0.0043 | 1.62E-08 | |
| rs7607716 | 2 | 162893338 | A | 0.0165 | 0.0025 | 2.62E-11 | |
| rs1247815 | 2 | 164379658 | T | 0.0108 | 0.0015 | 1.24E-13 | |
| rs1247738 | 2 | 166144850 | T | 0.0109 | 0.0017 | 7.15E-11 | |
| rs7575938 | 2 | 166910677 | A | 0.009 | 0.0015 | 1.13E-09 | |
| rs1450188 | 2 | 172616423 | A | 0.0095 | 0.0015 | 6.00E-10 | Not available |
| rs1742807 | 2 | 172851936 | C | 0.0137 | 0.0016 | 6.41E-17 | |
| rs1247398 | 2 | 173682948 | T | -0.0085 | 0.0015 | 3.57E-08 | |
| rs1019349 | 2 | 174094345 | A | 0.0097 | 0.0016 | 3.10E-09 | |

| | | | | | | |
|---|---|---|---|---|---|---|
| rs711793 | 2 | 174175921 | T | 0.0103 | 0.0015 | 5.01E-12 |
| rs7291750 | 2 | 175225018 | T | -0.0178 | 0.003 | 2.91E-09 |
| rs4972748 | 2 | 176098010 | T | 0.0105 | 0.0018 | 9.50E-09 |
| rs1261350 | 2 | 180928465 | C | 0.0098 | 0.0014 | 4.82E-12 |
| rs1766754 | 2 | 181614913 | A | -0.0087 | 0.0015 | 9.94E-09 |
| rs1835339 | 2 | 183393680 | T | 0.0084 | 0.0014 | 5.59E-09 |
| rs1527878 | 2 | 183446535 | A | -0.0105 | 0.0016 | 1.49E-10 |
| rs7289663 | 2 | 183543531 | T | 0.0162 | 0.0028 | 4.88E-09 |
| rs6217497 | 2 | 185053385 | A | -0.0105 | 0.0018 | 2.76E-09 |
| rs1154382 | 2 | 185883877 | T | -0.0169 | 0.003 | 2.96E-08 |
| rs1020505 | 2 | 186139826 | T | 0.009 | 0.0014 | 3.67E-10 |
| rs6217965 | 2 | 189135884 | A | 0.0112 | 0.0016 | 5.66E-13 |
| rs1066769 | 2 | 189580148 | A | -0.023 | 0.0041 | 1.38E-08 |
| rs6672197 | 2 | 191701185 | A | -0.0087 | 0.0015 | 1.51E-08 |
| rs5640852 | 2 | 193706511 | T | -0.0082 | 0.0014 | 7.47E-09 |
| rs1167547 | 2 | 193853269 | T | -0.0127 | 0.0014 | 1.37E-19 |
| rs4502401 | 2 | 199323526 | T | 0.0085 | 0.0015 | 7.55E-09 |
| rs1093182 | 2 | 199495830 | A | -0.0141 | 0.0014 | 8.65E-24 |
| rs6711399 | 2 | 200462840 | T | -0.0107 | 0.0019 | 1.16E-08 |
| rs1729412 | 2 | 201083598 | T | -0.0107 | 0.0014 | 4.05E-14 |
| rs4675248 | 2 | 202880230 | A | -0.0094 | 0.0014 | 4.50E-11 |
| rs6435326 | 2 | 207002559 | A | -0.0091 | 0.0014 | 7.56E-11 |
| rs1019283 | 2 | 207725616 | T | 0.0083 | 0.0015 | 2.05E-08 |
| rs6218299 | 2 | 212599303 | T | 0.0137 | 0.0015 | 2.55E-19 |
| rs6757087 | 2 | 212680523 | T | -0.0085 | 0.0014 | 2.55E-09 |
| rs7597126 | 2 | 215009358 | T | -0.0094 | 0.0014 | 2.46E-11 |
| rs1685177 | 2 | 215189950 | T | -0.0153 | 0.0027 | 9.24E-09 |
| rs4673840 | 2 | 215363241 | T | -0.0138 | 0.0019 | 5.86E-13 |
| rs3496708 | 2 | 215382654 | A | -0.008 | 0.0014 | 1.77E-08 |
| rs5701687 | 2 | 220310112 | T | 0.0209 | 0.0037 | 1.90E-08 |
| rs1542354 | 2 | 221814085 | A | -0.0084 | 0.0014 | 2.72E-09 |
| rs1301631 | 2 | 225337188 | A | -0.0138 | 0.0024 | 1.67E-08 |
| rs1168773 | 2 | 225405122 | A | -0.0105 | 0.0015 | 4.66E-12 |
| rs5714820 | 2 | 225689421 | A | 0.0095 | 0.0016 | 5.17E-09 |
| rs1301549 | 2 | 226290705 | C | 0.0119 | 0.0018 | 5.34E-11 |
| rs1269468 | 2 | 226609241 | T | 0.0102 | 0.0015 | 2.13E-11 |
| rs4500930 | 2 | 228985505 | T | -0.0109 | 0.0015 | 1.67E-13 |
| rs7474762 | 2 | 229061868 | T | 0.0153 | 0.0022 | 6.29E-12 |
| rs1246717 | 2 | 229189860 | T | -0.0081 | 0.0014 | 6.90E-09 |
| rs6704768 | 2 | 233592501 | A | -0.0116 | 0.0014 | 1.84E-16 |
| rs3581158 | 2 | 233743794 | T | 0.0217 | 0.0031 | 5.47E-12 |
| rs2250660 | 2 | 233752551 | C | 0.008 | 0.0014 | 1.20E-08 |
| rs4663617 | 2 | 236744626 | A | 0.0104 | 0.0017 | 4.12E-10 |
| rs6219091 | 2 | 236788175 | T | 0.0089 | 0.0015 | 1.25E-09 |
| rs7248213 | 2 | 237054356 | T | 0.0141 | 0.0019 | 4.59E-13 |
| rs6706275 | 2 | 240265617 | T | 0.0083 | 0.0015 | 2.81E-08 |
| rs7299379 | 2 | 240321051 | T | 0.0127 | 0.0022 | 1.12E-08 |

| | | | | | | | |
|---|---|---|---|---|---|---|---|
| rs1338833 | 2 | 242400597 | A | 0.0085 | 0.0015 | 6.48E-09 | |
| rs1704885 | 3 | 8258174 | A | 0.0108 | 0.0015 | 3.10E-13 | |
| rs382196 | 3 | 9145554 | T | -0.0082 | 0.0014 | 1.47E-08 | |
| rs164938 | 3 | 10315103 | T | -0.0088 | 0.0014 | 1.05E-09 | |
| rs9882532 | 3 | 16865845 | T | 0.0124 | 0.0015 | 2.15E-17 | |
| rs4685405 | 3 | 16981683 | T | -0.0111 | 0.0018 | 8.69E-10 | |
| rs7303907 | 3 | 17036387 | C | 0.0092 | 0.0016 | 1.75E-08 | |
| rs9866123 | 3 | 20631298 | A | 0.0086 | 0.0014 | 1.02E-09 | |
| rs399821 | 3 | 21500324 | A | -0.0078 | 0.0014 | 2.34E-08 | |
| rs1125127 | 3 | 24905460 | T | -0.0166 | 0.0027 | 9.91E-10 | |
| rs1308546 | 3 | 24950387 | C | 0.0103 | 0.0014 | 2.40E-13 | |
| rs7305555 | 3 | 28027538 | A | 0.0111 | 0.002 | 4.95E-08 | |
| rs1518890 | 3 | 34271042 | T | -0.009 | 0.0016 | 2.68E-08 | |
| rs4328757 | 3 | 36938180 | T | 0.0096 | 0.0014 | 2.95E-11 | |
| rs6226076 | 3 | 47990500 | C | -0.0132 | 0.0016 | 3.45E-16 | |
| rs1380961 | 3 | 48179876 | A | 0.0234 | 0.0042 | 1.95E-08 | |
| rs1407115 | 3 | 48469441 | C | 0.0366 | 0.0055 | 2.23E-11 | |
| rs6442126 | 3 | 48525955 | A | -0.0121 | 0.0015 | 4.72E-16 | |
| rs1919036 | 3 | 48578053 | A | 0.0335 | 0.0061 | 4.08E-08 | |
| rs3895736 | 3 | 48658467 | A | 0.0144 | 0.0019 | 9.92E-15 | |
| rs7308232 | 3 | 48981024 | C | -0.0252 | 0.0044 | 1.24E-08 | |
| rs7262491 | 3 | 49064576 | T | 0.0282 | 0.0034 | 5.59E-17 | |
| rs1502522 | 3 | 49296234 | A | 0.0204 | 0.0035 | 3.82E-09 | |
| rs1166563 | 3 | 49413266 | T | -0.0242 | 0.0038 | 2.95E-10 | |
| rs9859556 | 3 | 49455986 | T | 0.029 | 0.0015 | 4.61E-82 | |
| rs1384843 | 3 | 49511263 | T | -0.0306 | 0.0051 | 1.40E-09 | |
| rs6226267 | 3 | 49649873 | A | 0.0213 | 0.002 | 2.22E-25 | |
| rs7771938 | 3 | 49917021 | A | -0.0403 | 0.0058 | 3.78E-12 | |
| rs1112261 | 3 | 49931760 | T | -0.0201 | 0.0018 | 1.35E-27 | |
| rs2014830 | 3 | 50172397 | T | 0.011 | 0.0015 | 4.72E-13 | |
| rs2526398 | 3 | 50187596 | C | -0.0222 | 0.0015 | 2.47E-51 | |
| rs736471 | 3 | 50519392 | T | 0.0083 | 0.0014 | 4.69E-09 | |
| rs1172009 | 3 | 50562042 | T | 0.015 | 0.0026 | 6.48E-09 | |
| rs1171639 | 3 | 50699715 | A | 0.0159 | 0.0021 | 1.89E-14 | |
| rs1823553 | 3 | 50920457 | A | -0.0525 | 0.0084 | 3.67E-10 | Excluded Low MAF |
| rs1504216 | 3 | 51047621 | A | 0.0349 | 0.0061 | 1.04E-08 | |
| rs7445387 | 3 | 52046707 | A | 0.0214 | 0.0037 | 6.48E-09 | |
| rs2336721 | 3 | 53028375 | T | 0.0095 | 0.0015 | 1.05E-10 | |
| rs6225281 | 3 | 53440128 | A | -0.0094 | 0.0017 | 1.20E-08 | |
| rs4687735 | 3 | 53684371 | T | -0.02 | 0.0033 | 1.63E-09 | |
| rs1113038 | 3 | 53758950 | T | 0.009 | 0.0014 | 4.17E-10 | |
| rs6445633 | 3 | 54230638 | A | 0.0088 | 0.0015 | 6.37E-09 | |
| rs7625428 | 3 | 56570907 | T | 0.0098 | 0.0014 | 1.07E-11 | |
| rs2736752 | 3 | 60817322 | T | 0.0095 | 0.0017 | 3.24E-08 | |
| rs6774533 | 3 | 62471086 | T | 0.009 | 0.0015 | 4.18E-09 | |
| rs853286 | 3 | 64285502 | T | 0.0124 | 0.0023 | 4.13E-08 | |
| rs6224744 | 3 | 64430551 | C | 0.0114 | 0.0014 | 8.79E-16 | |

| | | | | | | |
|---|---|---|---|---|---|---|
| rs1404549 | 3 | 65454923 | A | -0.0089 | 0.0015 | 2.38E-09 |
| rs1263807 | 3 | 65654480 | A | -0.0107 | 0.0015 | 3.13E-12 |
| rs7755409 | 3 | 65711935 | T | -0.0186 | 0.0027 | 6.63E-12 |
| rs6780414 | 3 | 67384670 | T | 0.0105 | 0.0018 | 7.10E-09 |
| rs9849884 | 3 | 68869923 | A | -0.009 | 0.0014 | 2.51E-10 |
| rs1150005 | 3 | 69930625 | A | -0.025 | 0.0031 | 1.09E-15 |
| rs5630688 | 3 | 70253693 | A | 0.0086 | 0.0016 | 4.82E-08 |
| rs6225628 | 3 | 70526305 | T | 0.0111 | 0.0017 | 1.32E-10 |
| rs7626786 | 3 | 70540347 | A | 0.0104 | 0.0017 | 1.64E-09 |
| rs9877225 | 3 | 70631590 | T | 0.009 | 0.0016 | 4.22E-08 |
| rs1191983 | 3 | 70954273 | C | 0.0089 | 0.0015 | 7.95E-10 |
| rs5573631 | 3 | 71586293 | C | -0.0154 | 0.0014 | 4.59E-27 |
| rs5996735 | 3 | 71639294 | T | -0.0127 | 0.0021 | 2.34E-09 |
| rs9820604 | 3 | 72363674 | T | 0.0115 | 0.0018 | 4.80E-10 |
| rs1918394 | 3 | 74903904 | T | 0.0119 | 0.0019 | 3.04E-10 |
| rs1123794 | 3 | 76230061 | C | -0.0082 | 0.0014 | 7.10E-09 |
| rs9844755 | 3 | 78440564 | A | 0.0083 | 0.0015 | 4.64E-08 |
| rs1331898 | 3 | 82533639 | A | 0.0124 | 0.0015 | 1.17E-16 |
| rs9830359 | 3 | 85397118 | T | -0.0147 | 0.0022 | 4.70E-11 |
| rs6656892 | 3 | 85672018 | T | -0.0181 | 0.0015 | 7.60E-34 |
| rs5743740 | 3 | 86173623 | A | -0.0145 | 0.0023 | 2.67E-10 |
| rs4396896 | 3 | 86261800 | A | -0.0107 | 0.0014 | 1.12E-13 |
| rs9853928 | 3 | 103295384 | T | -0.0127 | 0.0017 | 1.59E-13 |
| rs2851388 | 3 | 107296628 | A | -0.011 | 0.0018 | 2.03E-09 |
| rs7640424 | 3 | 107820063 | T | 0.0113 | 0.0015 | 1.02E-13 |
| rs1149582 | 3 | 108039298 | T | -0.022 | 0.0039 | 1.36E-08 |
| rs2290601 | 3 | 108112760 | T | -0.0149 | 0.0017 | 4.49E-19 |
| rs1156933 | 3 | 108284214 | A | 0.0309 | 0.0053 | 7.22E-09 |
| rs7430651 | 3 | 116582186 | T | -0.0111 | 0.0016 | 9.38E-13 |
| rs1191770 | 3 | 117174568 | A | -0.0087 | 0.0014 | 9.45E-10 |
| rs2088913 | 3 | 117310069 | A | 0.0081 | 0.0014 | 9.61E-09 |
| rs1717204 | 3 | 118461145 | A | -0.0117 | 0.0018 | 1.20E-10 |
| rs6792805 | 3 | 118675196 | A | -0.009 | 0.0016 | 1.70E-08 |
| rs2332179 | 3 | 122079430 | A | 0.0121 | 0.002 | 2.76E-09 |
| rs1172098 | 3 | 122214045 | T | 0.0085 | 0.0015 | 2.57E-08 |
| rs3406738 | 3 | 123594419 | T | -0.0104 | 0.0015 | 9.77E-13 |
| rs7630133 | 3 | 123923288 | A | 0.0082 | 0.0015 | 2.80E-08 |
| rs9821664 | 3 | 127003110 | A | 0.0094 | 0.0017 | 4.52E-08 |
| rs9289300 | 3 | 127144988 | T | -0.0164 | 0.0019 | 1.00E-17 |
| rs2929860 | 3 | 128691941 | T | 0.011 | 0.0018 | 1.33E-09 |
| rs7650602 | 3 | 141147414 | T | -0.0089 | 0.0014 | 2.77E-10 |
| rs2885198 | 3 | 143636421 | A | 0.0088 | 0.0014 | 4.63E-10 |
| rs1205416 | 3 | 150093445 | C | 0.0096 | 0.0016 | 2.30E-09 |
| rs7387433 | 3 | 157933483 | T | -0.0178 | 0.0029 | 1.29E-09 |
| rs7617204 | 3 | 160821969 | A | -0.0097 | 0.0014 | 4.33E-12 |
| rs9858921 | 3 | 161148946 | A | -0.0087 | 0.0014 | 5.19E-10 |
| rs893522 | 3 | 165560618 | A | 0.0149 | 0.0025 | 1.39E-09 |

| | | | | | | | |
|---|---|---|---|---|---|---|---|
| rs1309916 | 3 | 168740425 | T | -0.0118 | 0.0017 | 1.09E-11 | |
| rs4894658 | 3 | 173928736 | C | 0.0091 | 0.0016 | 1.34E-08 | |
| rs7262255 | 3 | 175672897 | T | -0.012 | 0.0017 | 5.50E-13 | |
| rs1157327 | 3 | 180691034 | A | -0.0415 | 0.0072 | 1.06E-08 | Excluded Low MAF |
| rs7702523 | 3 | 180734185 | A | -0.014 | 0.0019 | 4.65E-13 | |
| rs2718791 | 3 | 180931395 | T | -0.0081 | 0.0015 | 4.02E-08 | |
| rs9870317 | 3 | 182514636 | A | 0.0086 | 0.0015 | 9.35E-09 | |
| rs1093724 | 3 | 185802823 | T | -0.0123 | 0.0018 | 6.95E-12 | |
| rs1332748 | 3 | 196788363 | A | -0.0121 | 0.0018 | 4.47E-11 | |
| rs1505375 | 4 | 705629 | A | -0.0151 | 0.0026 | 9.50E-09 | |
| rs9683585 | 4 | 2702804 | C | 0.0084 | 0.0014 | 2.46E-09 | |
| rs6824567 | 4 | 2883172 | T | 0.013 | 0.0016 | 8.91E-17 | |
| rs363096 | 4 | 3180021 | T | -0.0143 | 0.0014 | 5.77E-24 | |
| rs362307 | 4 | 3241845 | T | -0.0226 | 0.0027 | 1.55E-17 | |
| rs3415584 | 4 | 3252130 | A | 0.0135 | 0.0016 | 1.43E-16 | |
| rs6853599 | 4 | 3445610 | T | 0.0108 | 0.0019 | 6.27E-09 | |
| rs1120924 | 4 | 5228939 | T | 0.0108 | 0.0015 | 2.79E-13 | |
| rs2702576 | 4 | 15058245 | A | -0.009 | 0.0014 | 4.41E-10 | |
| rs1003294 | 4 | 15402115 | T | -0.008 | 0.0015 | 3.42E-08 | |
| rs1313321 | 4 | 15646794 | A | 0.0096 | 0.0014 | 7.57E-12 | |
| rs1980129 | 4 | 17045911 | A | 0.0085 | 0.0014 | 1.17E-09 | |
| rs1141928 | 4 | 17969142 | T | -0.0279 | 0.0051 | 3.26E-08 | |
| rs2011603 | 4 | 18025484 | A | -0.0091 | 0.0016 | 1.12E-08 | |
| rs4467547 | 4 | 21945933 | T | 0.0121 | 0.0014 | 4.88E-17 | |
| rs9291437 | 4 | 22165255 | C | -0.0081 | 0.0014 | 1.82E-08 | |
| rs1173686 | 4 | 23729566 | A | 0.0142 | 0.0018 | 2.56E-15 | |
| rs3481147 | 4 | 25408838 | A | 0.0104 | 0.0017 | 2.75E-09 | |
| rs6240939 | 4 | 25605036 | T | 0.0101 | 0.0018 | 3.55E-08 | |
| rs1002173 | 4 | 28609476 | T | -0.0164 | 0.0027 | 9.45E-10 | |
| rs1173343 | 4 | 30534692 | A | -0.0096 | 0.0017 | 2.10E-08 | |
| rs4358358 | 4 | 30970822 | A | 0.0109 | 0.0015 | 8.69E-13 | |
| rs868456 | 4 | 32150029 | A | 0.0089 | 0.0015 | 1.07E-08 | |
| rs317050 | 4 | 35446302 | T | 0.0078 | 0.0014 | 2.50E-08 | |
| rs6179858 | 4 | 36691695 | A | -0.0112 | 0.002 | 2.05E-08 | |
| rs1000951 | 4 | 38591860 | A | 0.0088 | 0.0014 | 3.02E-10 | |
| rs6812533 | 4 | 39687838 | T | 0.0106 | 0.0016 | 4.85E-11 | |
| rs1313076 | 4 | 45163333 | C | -0.0091 | 0.0014 | 1.89E-10 | |
| rs1264621 | 4 | 45980056 | T | 0.0084 | 0.0014 | 4.56E-09 | |
| rs2055940 | 4 | 46997913 | A | 0.0082 | 0.0015 | 4.95E-08 | |
| rs1505676 | 4 | 62127291 | C | 0.0084 | 0.0015 | 6.67E-09 | |
| rs1440930 | 4 | 65902342 | C | 0.0081 | 0.0014 | 8.94E-09 | |
| rs7494485 | 4 | 66496914 | T | -0.0089 | 0.0015 | 3.62E-09 | |
| rs1173216 | 4 | 67070824 | A | 0.0105 | 0.0016 | 1.46E-11 | |
| rs1250622 | 4 | 67899235 | T | -0.013 | 0.0014 | 4.23E-20 | |
| rs7263669 | 4 | 68016609 | T | -0.014 | 0.002 | 1.31E-12 | |
| rs2034631 | 4 | 82225529 | T | 0.0089 | 0.0015 | 2.64E-09 | |
| rs7692359 | 4 | 83209346 | T | -0.0109 | 0.0017 | 1.40E-10 | |

| | | | | | | |
|---|---|---|---|---|---|---|
| rs1718610 | 4 | 91703241 | T | 0.0108 | 0.0018 | 3.95E-09 |
| rs2870281 | 4 | 91945572 | A | -0.0083 | 0.0014 | 4.35E-09 |
| rs1250352 | 4 | 94543233 | T | -0.0113 | 0.0016 | 3.93E-13 |
| rs151381 | 4 | 103118768 | T | -0.0087 | 0.0014 | 7.01E-10 |
| rs1310732 | 4 | 103188709 | T | -0.0239 | 0.0027 | 1.98E-18 |
| rs7683416 | 4 | 106152984 | T | 0.0139 | 0.0014 | 6.05E-23 |
| rs2522545 | 4 | 106378284 | T | -0.0083 | 0.0014 | 6.23E-09 |
| rs1490612 | 4 | 106410855 | T | 0.0193 | 0.0026 | 1.55E-13 |
| rs925161 | 4 | 112323249 | C | -0.0078 | 0.0014 | 3.38E-08 |
| rs1955250 | 4 | 118514236 | A | -0.014 | 0.0025 | 2.26E-08 |
| rs1314565 | 4 | 122963344 | T | -0.0173 | 0.0025 | 8.99E-12 |
| rs6534338 | 4 | 123026869 | T | 0.0103 | 0.0015 | 1.46E-11 |
| rs1000623 | 4 | 130670108 | T | -0.0093 | 0.0016 | 4.48E-09 |
| rs1704880 | 4 | 137529847 | A | -0.0106 | 0.0014 | 2.75E-13 |
| rs1051950 | 4 | 140640283 | T | 0.0119 | 0.0019 | 7.31E-10 |
| rs7624160 | 4 | 140717160 | A | 0.0139 | 0.0022 | 6.48E-10 |
| rs1264377 | 4 | 140753103 | T | 0.013 | 0.0015 | 1.32E-17 |
| rs3531965 | 4 | 140903155 | T | 0.0119 | 0.0015 | 8.01E-16 |
| rs969512 | 4 | 147872742 | A | -0.0106 | 0.0015 | 9.84E-13 |
| rs7672622 | 4 | 157705551 | A | -0.0088 | 0.0016 | 4.72E-08 |
| rs2081652 | 4 | 159862913 | A | 0.0123 | 0.0015 | 1.09E-16 |
| rs1110023 | 4 | 160477041 | A | -0.0101 | 0.0014 | 8.17E-13 |
| rs4691601 | 4 | 160599341 | A | -0.0115 | 0.0014 | 2.46E-16 |
| rs1841023 | 4 | 163735584 | A | -0.0092 | 0.0015 | 1.64E-09 |
| rs5640513 | 4 | 166126745 | A | 0.0153 | 0.0022 | 1.31E-12 |
| rs1264652 | 4 | 170944698 | T | -0.0132 | 0.0016 | 3.79E-16 |
| rs1137316 | 4 | 170984229 | T | 0.0339 | 0.0061 | 2.36E-08 |
| rs2082317 | 4 | 171824549 | T | -0.0078 | 0.0014 | 3.19E-08 |
| rs1173265 | 4 | 172427073 | A | -0.0126 | 0.0016 | 6.24E-15 |
| rs1311785 | 4 | 174008759 | A | 0.0084 | 0.0015 | 4.64E-08 |
| rs1759867 | 4 | 176647637 | T | -0.0116 | 0.0014 | 1.37E-16 |
| rs6234063 | 4 | 176940335 | T | 0.0103 | 0.0014 | 1.47E-12 |
| rs7270956 | 4 | 178540163 | A | 0.0085 | 0.0015 | 2.03E-08 |
| rs1128956 | 4 | 183724005 | T | -0.014 | 0.0019 | 5.34E-14 |
| rs1172469 | 4 | 186764747 | T | 0.0093 | 0.0015 | 2.08E-09 |
| rs1316384 | 5 | 3264389 | T | -0.0149 | 0.002 | 4.66E-14 |
| rs1687180 | 5 | 3437297 | T | 0.009 | 0.0015 | 1.59E-09 |
| rs162445 | 5 | 7943246 | A | 0.0152 | 0.0026 | 5.72E-09 |
| rs2434672 | 5 | 11471879 | A | 0.0091 | 0.0014 | 1.06E-10 |
| rs1690168 | 5 | 11507368 | T | 0.0111 | 0.0019 | 5.17E-09 |
| rs4263475 | 5 | 26774142 | A | 0.0078 | 0.0014 | 4.40E-08 |
| rs1150171 | 5 | 26896955 | A | -0.0206 | 0.0038 | 4.72E-08 |
| rs1316813 | 5 | 26898629 | A | -0.0092 | 0.0017 | 2.57E-08 |
| rs1756346 | 5 | 26913774 | A | -0.0147 | 0.0017 | 1.74E-17 |
| rs1094092 | 5 | 30808645 | T | 0.0111 | 0.0014 | 1.31E-14 |
| rs1316918 | 5 | 51716957 | A | -0.0079 | 0.0014 | 2.01E-08 |
| rs6237051 | 5 | 52789198 | A | -0.0109 | 0.0019 | 5.59E-09 |

| | | | | | | |
|---|---|---|---|---|---|---|
| rs702606 | 5 | 53167117 | T | 0.0118 | 0.0021 | 9.19E-09 |
| rs1094054 | 5 | 56677646 | A | 0.0078 | 0.0014 | 3.44E-08 |
| rs981883 | 5 | 57107989 | A | 0.0099 | 0.0014 | 6.67E-12 |
| rs2964199 | 5 | 57532775 | T | -0.0099 | 0.0015 | 4.61E-11 |
| rs6450476 | 5 | 57771087 | A | -0.0116 | 0.0015 | 4.62E-14 |
| rs1336104 | 5 | 58318963 | T | -0.0158 | 0.0028 | 1.84E-08 |
| rs981230 | 5 | 59039858 | T | 0.01 | 0.0014 | 1.28E-12 |
| rs7979816 | 5 | 59152140 | A | 0.0172 | 0.0025 | 3.63E-12 |
| rs2910823 | 5 | 59498175 | T | 0.0115 | 0.0014 | 2.50E-16 |
| rs4283754 | 5 | 59570258 | A | -0.0084 | 0.0014 | 7.55E-09 |
| rs1315442 | 5 | 59608950 | C | -0.0136 | 0.0023 | 5.02E-09 |
| rs27220 | 5 | 59775136 | A | -0.0114 | 0.0015 | 7.52E-15 |
| rs6449503 | 5 | 60095272 | A | 0.0185 | 0.0014 | 1.84E-39 |
| rs7985592 | 5 | 60095384 | T | -0.0194 | 0.0033 | 3.10E-09 |
| rs1135521 | 5 | 60644403 | C | 0.0294 | 0.0053 | 2.51E-08 |
| rs4490539 | 5 | 60685757 | A | 0.0142 | 0.0015 | 1.31E-20 |
| rs1317703 | 5 | 60853523 | A | -0.0085 | 0.0015 | 4.88E-09 |
| rs347661 | 5 | 62549141 | T | -0.0081 | 0.0014 | 1.58E-08 |
| rs1080538 | 5 | 63034606 | A | -0.0103 | 0.0014 | 2.22E-13 |
| rs6722496 | 5 | 63787810 | A | 0.0113 | 0.0017 | 7.19E-11 |
| rs1319023 | 5 | 65964783 | T | -0.0138 | 0.0024 | 5.66E-09 |
| rs34309 | 5 | 67564383 | A | 0.008 | 0.0014 | 2.94E-08 |
| rs1004140 | 5 | 67805272 | T | 0.0151 | 0.0019 | 1.55E-15 |
| rs7716161 | 5 | 72179574 | C | 0.0105 | 0.0019 | 3.00E-08 |
| rs42302 | 5 | 74965554 | A | 0.0086 | 0.0015 | 5.96E-09 |
| rs1910005 | 5 | 81110866 | T | -0.009 | 0.0015 | 5.66E-09 |
| rs7356536 | 5 | 87083961 | T | -0.0129 | 0.0021 | 2.95E-10 |
| rs6452793 | 5 | 87818174 | T | 0.0175 | 0.0017 | 4.47E-26 |
| rs324885 | 5 | 87897271 | A | 0.0142 | 0.0014 | 4.11E-24 |
| rs7277186 | 5 | 87974682 | T | -0.0211 | 0.0029 | 3.64E-13 |
| rs1417296 | 5 | 87999371 | T | 0.0195 | 0.0026 | 1.49E-13 |
| rs6110461 | 5 | 88163771 | A | -0.0172 | 0.0014 | 1.56E-34 |
| rs1004282 | 5 | 88308777 | A | -0.0145 | 0.0022 | 1.06E-10 |
| rs2365376 | 5 | 88740323 | A | 0.0092 | 0.0015 | 3.93E-10 |
| rs1004606 | 5 | 89115576 | A | 0.0127 | 0.0022 | 4.40E-09 |
| rs2962378 | 5 | 90950210 | A | 0.0101 | 0.0017 | 4.30E-09 |
| rs1766933 | 5 | 92187932 | T | -0.011 | 0.0014 | 1.10E-14 |
| rs6881733 | 5 | 92586991 | T | -0.0086 | 0.0016 | 4.08E-08 |
| rs1144685 | 5 | 92870246 | A | 0.0293 | 0.0036 | 1.99E-16 |
| rs2029401 | 5 | 92891029 | A | -0.0091 | 0.0014 | 1.53E-10 |
| rs1233273 | 5 | 93033082 | A | 0.0135 | 0.0018 | 5.96E-14 |
| rs1007417 | 5 | 93927705 | A | 0.0079 | 0.0014 | 1.58E-08 |
| rs187580 | 5 | 102627355 | T | -0.0104 | 0.0017 | 3.56E-10 |
| rs7494427 | 5 | 102726073 | T | 0.0231 | 0.0035 | 4.44E-11 |
| rs171697 | 5 | 103956516 | C | 0.0133 | 0.0015 | 4.85E-19 |
| rs1006142 | 5 | 104678079 | A | 0.0082 | 0.0014 | 6.63E-09 |
| rs7714719 | 5 | 106498927 | C | 0.0116 | 0.002 | 1.12E-08 |

| | | | | | | |
|---|---|---|---|---|---|---|
| rs1593022 | 5 | 106685563 | T | -0.0099 | 0.0017 | 4.69E-09 |
| rs252991 | 5 | 106767346 | A | 0.0095 | 0.0015 | 6.30E-11 |
| rs1316306 | 5 | 106955451 | T | 0.0115 | 0.0014 | 4.43E-16 |
| rs34410 | 5 | 107469684 | C | -0.0083 | 0.0014 | 3.86E-09 |
| rs2416214 | 5 | 108943485 | A | 0.0079 | 0.0014 | 2.09E-08 |
| rs1748964 | 5 | 109156184 | A | 0.0127 | 0.0015 | 2.77E-17 |
| rs4705763 | 5 | 112394503 | C | 0.0089 | 0.0014 | 2.08E-10 |
| rs660001 | 5 | 113866598 | A | -0.0164 | 0.0017 | 1.66E-21 |
| rs6237983 | 5 | 120102028 | T | 0.0121 | 0.0015 | 1.77E-15 |
| rs1006002 | 5 | 124304677 | T | 0.011 | 0.0015 | 1.88E-13 |
| rs1005759 | 5 | 124335247 | A | 0.0106 | 0.0014 | 4.08E-14 |
| rs6871635 | 5 | 133830395 | A | -0.0087 | 0.0014 | 1.16E-09 |
| rs713584 | 5 | 136416957 | A | -0.008 | 0.0014 | 1.48E-08 |
| rs1434630 | 5 | 136557055 | T | -0.0137 | 0.002 | 4.63E-12 |
| rs1251907 | 5 | 136776762 | T | -0.0107 | 0.0017 | 1.16E-10 |
| rs6881581 | 5 | 140946355 | A | -0.0096 | 0.0015 | 1.08E-10 |
| rs32940 | 5 | 141132286 | T | -0.0096 | 0.0015 | 3.72E-10 |
| rs3519210 | 5 | 145705669 | T | -0.0091 | 0.0017 | 3.77E-08 |
| rs1251699 | 5 | 145927688 | C | -0.0099 | 0.0017 | 7.90E-09 |
| rs2964255 | 5 | 152052332 | A | 0.0088 | 0.0015 | 6.34E-09 |
| rs1829021 | 5 | 153304758 | A | -0.03 | 0.0053 | 1.17E-08 |
| rs42210 | 5 | 166408788 | C | -0.0098 | 0.0016 | 2.74E-10 |
| rs1706964 | 5 | 167452251 | T | 0.0085 | 0.0015 | 2.01E-08 |
| rs2569041 | 5 | 167510357 | T | 0.0077 | 0.0014 | 4.85E-08 |
| rs1265977 | 5 | 167621274 | T | 0.0085 | 0.0014 | 1.64E-09 |
| rs29792 | 5 | 168978148 | A | -0.0093 | 0.0015 | 1.06E-09 |
| rs6861925 | 5 | 176324729 | C | -0.009 | 0.0014 | 1.76E-10 |
| rs2545795 | 5 | 176887106 | A | 0.0125 | 0.0014 | 1.03E-18 |
| rs2858777 | 5 | 177030177 | T | 0.0088 | 0.0015 | 2.55E-09 |
| rs1252479 | 6 | 3464074 | T | 0.0102 | 0.0014 | 5.28E-13 |
| rs1126037 | 6 | 6069588 | A | -0.0131 | 0.0022 | 2.72E-09 |
| rs7282475 | 6 | 14619148 | T | 0.0133 | 0.0024 | 1.93E-08 |
| rs9349956 | 6 | 14718260 | A | -0.0159 | 0.0018 | 4.89E-18 |
| rs1094926 | 6 | 14886977 | A | -0.0091 | 0.0016 | 2.33E-08 |
| rs7772172 | 6 | 16662928 | A | 0.0091 | 0.0014 | 1.74E-10 |
| rs7766240 | 6 | 16954368 | T | -0.0134 | 0.0024 | 2.00E-08 |
| rs7282985 | 6 | 16966052 | A | -0.0149 | 0.0016 | 1.52E-19 |
| rs7282851 | 6 | 19036035 | T | -0.0168 | 0.0019 | 1.38E-19 |
| rs2092248 | 6 | 19205458 | T | -0.0083 | 0.0015 | 2.43E-08 |
| rs4712371 | 6 | 19227667 | A | -0.0124 | 0.0021 | 6.63E-09 |
| rs9465509 | 6 | 19744129 | A | -0.0084 | 0.0014 | 2.35E-09 |
| rs767943 | 6 | 23446691 | A | -0.0122 | 0.0016 | 1.54E-14 |
| rs7871422 | 6 | 25793055 | T | 0.0172 | 0.003 | 1.55E-08 |
| rs7283469 | 6 | 26176517 | A | 0.0117 | 0.002 | 4.91E-09 |
| rs2179152 | 6 | 26325888 | T | -0.0131 | 0.0015 | 1.68E-19 |
| rs6918506 | 6 | 26577857 | C | 0.0113 | 0.0014 | 9.57E-16 |
| rs6009664 | 6 | 27636313 | A | 0.0159 | 0.0023 | 2.24E-12 |

| | | | | | | |
|---|---|---|---|---|---|---|
| rs7451726 | 6 | 28382494 | A | 0.0109 | 0.0017 | 2.86E-10 |
| rs1148107 | 6 | 30398390 | A | -0.0121 | 0.002 | 1.98E-09 |
| rs2256965 | 6 | 31555130 | A | 0.0106 | 0.0014 | 2.40E-13 |
| rs9267658 | 6 | 31845985 | T | 0.0138 | 0.002 | 5.25E-12 |
| rs9267677 | 6 | 31892641 | T | 0.018 | 0.0024 | 3.66E-14 |
| rs429150 | 6 | 32075563 | T | 0.0085 | 0.0014 | 1.50E-09 |
| rs1061801 | 6 | 33282338 | A | -0.011 | 0.0018 | 1.06E-09 |
| rs1094743 | 6 | 33774948 | T | 0.009 | 0.0015 | 1.11E-09 |
| rs5734979 | 6 | 37486052 | A | 0.0093 | 0.0014 | 8.63E-11 |
| rs2436760 | 6 | 40374288 | T | 0.0232 | 0.0042 | 4.52E-08 |
| rs912883 | 6 | 41552040 | T | 0.0087 | 0.0015 | 6.20E-09 |
| rs7864810 | 6 | 50683009 | T | -0.0141 | 0.0025 | 1.70E-08 |
| rs7449561 | 6 | 66215549 | A | 0.0102 | 0.0017 | 9.73E-10 |
| rs9294770 | 6 | 68242866 | T | 0.0083 | 0.0014 | 8.74E-09 |
| rs9446060 | 6 | 69552271 | A | 0.0077 | 0.0014 | 4.33E-08 |
| rs9442750 | 6 | 72886012 | A | -0.0089 | 0.0016 | 3.15E-08 |
| rs1321204 | 6 | 78171124 | T | 0.0101 | 0.0018 | 7.42E-09 |
| rs1094358 | 6 | 79510994 | A | 0.0084 | 0.0015 | 1.05E-08 |
| rs4352658 | 6 | 88279872 | T | -0.019 | 0.0026 | 1.71E-13 |
| rs9359939 | 6 | 92133241 | A | -0.0105 | 0.0016 | 1.51E-10 |
| rs1768664 | 6 | 93568527 | T | -0.0098 | 0.0017 | 6.37E-09 |
| rs633279 | 6 | 93840705 | A | -0.0089 | 0.0015 | 8.12E-09 |
| rs1408284 | 6 | 93893586 | C | -0.0127 | 0.002 | 5.71E-10 |
| rs7775100 | 6 | 96519657 | T | -0.0087 | 0.0014 | 1.20E-09 |
| rs3510449 | 6 | 97390463 | A | 0.0124 | 0.0018 | 6.81E-12 |
| rs6569077 | 6 | 98212409 | T | 0.0118 | 0.0014 | 3.29E-16 |
| rs1123757 | 6 | 98224217 | A | 0.0184 | 0.0021 | 6.00E-19 |
| rs6242038 | 6 | 98404243 | T | -0.0184 | 0.0027 | 6.16E-12 |
| rs9490512 | 6 | 98537993 | A | -0.0136 | 0.0014 | 4.00E-22 |
| rs9375188 | 6 | 98555272 | T | 0.0212 | 0.0014 | 8.78E-52 |
| rs6932108 | 6 | 98713992 | C | -0.021 | 0.0027 | 6.10E-15 |
| rs9375403 | 6 | 98831636 | C | 0.0126 | 0.0016 | 1.66E-14 |
| rs1728118 | 6 | 108029518 | A | 0.0089 | 0.0016 | 1.32E-08 |
| rs1170059 | 6 | 114218608 | T | 0.015 | 0.0022 | 9.35E-12 |
| rs1008064 | 6 | 114742335 | A | 0.0124 | 0.002 | 7.05E-10 |
| rs6745686 | 6 | 119079800 | A | 0.0119 | 0.0018 | 5.37E-11 |
| rs1154266 | 6 | 119215402 | A | 0.0112 | 0.0015 | 2.44E-13 |
| rs9388490 | 6 | 126704795 | T | 0.0091 | 0.0014 | 1.15E-10 |
| rs1720590 | 6 | 127764305 | T | -0.0092 | 0.0015 | 7.71E-10 |
| rs1319725 | 6 | 128333682 | T | 0.0109 | 0.0016 | 2.83E-12 |
| rs9492774 | 6 | 131302489 | C | 0.008 | 0.0015 | 4.33E-08 |
| rs9373363 | 6 | 143150043 | A | -0.0111 | 0.0016 | 5.54E-12 |
| rs4895650 | 6 | 145024122 | T | 0.0093 | 0.0014 | 5.31E-11 |
| rs6917204 | 6 | 145284716 | T | 0.0119 | 0.0017 | 8.14E-12 |
| rs9386110 | 6 | 145626208 | T | -0.0163 | 0.0023 | 5.66E-13 |
| rs3428683 | 6 | 148273664 | T | -0.0089 | 0.0014 | 3.44E-10 |
| rs1115581 | 6 | 152149435 | T | -0.0129 | 0.0023 | 1.57E-08 |

| | | | | | | |
|---|---|---|---|---|---|---|
| rs6557171 | 6 | 152234593 | T | -0.0155 | 0.0015 | 5.48E-25 |
| rs1933264 | 6 | 153391618 | T | 0.01 | 0.0016 | 5.85E-10 |
| rs9371883 | 6 | 155644978 | C | -0.0087 | 0.0015 | 3.42E-09 |
| rs1135883 | 6 | 157236426 | T | -0.0104 | 0.0017 | 1.67E-09 |
| rs2023016 | 6 | 162981266 | C | 0.0107 | 0.0017 | 4.91E-10 |
| rs6917154 | 6 | 167135156 | T | 0.0127 | 0.0021 | 2.55E-09 |
| rs6924023 | 6 | 170076041 | A | 0.0107 | 0.0016 | 5.17E-11 |
| rs4719460 | 7 | 859610 | T | -0.0088 | 0.0015 | 2.34E-09 |
| rs1177223 | 7 | 1856273 | T | 0.0177 | 0.0019 | 1.20E-20 |
| rs1026457 | 7 | 1944591 | A | 0.0175 | 0.0032 | 3.79E-08 |
| rs5608518 | 7 | 2143594 | A | -0.0269 | 0.0037 | 5.94E-13 |
| rs1637770 | 7 | 2190385 | T | 0.0191 | 0.0028 | 7.32E-12 |
| rs1324040 | 7 | 2194396 | T | 0.0176 | 0.0017 | 2.36E-24 |
| rs5892170 | 7 | 2322943 | T | 0.0109 | 0.0015 | 7.84E-14 |
| rs1713329 | 7 | 3455401 | A | -0.0146 | 0.0023 | 2.28E-10 |
| rs4719944 | 7 | 3496032 | T | -0.0121 | 0.0014 | 9.25E-18 |
| rs3574545 | 7 | 5824100 | A | 0.008 | 0.0014 | 1.43E-08 |
| rs6946362 | 7 | 6574807 | T | 0.0091 | 0.0015 | 1.89E-09 |
| rs929511 | 7 | 8003017 | T | -0.016 | 0.0021 | 3.91E-14 |
| rs1756838 | 7 | 8091876 | A | 0.0127 | 0.0014 | 1.62E-19 |
| rs1181348 | 7 | 11500372 | T | -0.0223 | 0.003 | 1.50E-13 |
| rs1223436 | 7 | 11893275 | A | 0.0087 | 0.0015 | 9.61E-09 |
| rs6977237 | 7 | 14027991 | T | 0.0082 | 0.0014 | 5.59E-09 |
| rs6461536 | 7 | 21131807 | A | 0.0088 | 0.0015 | 1.28E-08 |
| rs3592992 | 7 | 21400525 | A | -0.0114 | 0.0016 | 1.77E-12 |
| rs1049953 | 7 | 21690606 | A | 0.008 | 0.0014 | 1.40E-08 |
| rs2529069 | 7 | 24571038 | T | -0.0106 | 0.0016 | 2.66E-11 |
| rs7926543 | 7 | 24621381 | A | -0.0197 | 0.0022 | 1.98E-19 |
| rs9771228 | 7 | 32322496 | T | 0.0096 | 0.0015 | 5.91E-11 |
| rs1176538 | 7 | 32494790 | T | 0.0086 | 0.0015 | 9.40E-09 |
| rs1717051 | 7 | 32861230 | C | 0.0132 | 0.0023 | 1.99E-08 |
| rs1095159 | 7 | 39091043 | T | -0.0109 | 0.0015 | 3.62E-13 |
| rs1132057 | 7 | 39246987 | A | -0.0135 | 0.0024 | 2.00E-08 |
| rs1324622 | 7 | 39324253 | T | -0.0093 | 0.0014 | 8.53E-11 |
| rs7796103 | 7 | 41820155 | C | -0.0077 | 0.0014 | 3.81E-08 |
| rs4724083 | 7 | 42007071 | T | 0.0083 | 0.0015 | 3.53E-08 |
| rs2850528 | 7 | 44377654 | C | 0.0106 | 0.0018 | 3.50E-09 |
| rs3735478 | 7 | 44800176 | T | 0.0108 | 0.0016 | 3.66E-12 |
| rs1267037 | 7 | 48823154 | A | 0.0092 | 0.0014 | 5.99E-11 |
| rs1755106 | 7 | 49869771 | A | 0.0141 | 0.0019 | 1.53E-13 |
| rs2299156 | 7 | 50764404 | T | 0.0099 | 0.0018 | 3.15E-08 |
| rs5895008 | 7 | 53857821 | T | 0.0095 | 0.0017 | 4.77E-08 |
| rs1220779 | 7 | 54734673 | A | -0.0079 | 0.0014 | 1.58E-08 |
| rs7870239 | 7 | 68618770 | C | 0.0167 | 0.003 | 2.38E-08 |
| rs7803932 | 7 | 70203673 | A | 0.0127 | 0.0019 | 1.74E-11 |
| rs7819315 | 7 | 71720681 | A | 0.0191 | 0.0029 | 2.76E-11 |
| rs3541770 | 7 | 71739916 | T | -0.0152 | 0.0014 | 2.40E-27 |

| | | | | | | |
|---|---|---|---|---|---|---|
| rs2718277 | 7 | 74133929 | T | 0.0142 | 0.0026 | 2.99E-08 |
| rs1167827 | 7 | 75163169 | A | 0.0104 | 0.0014 | 2.93E-13 |
| rs6071774 | 7 | 75846083 | C | -0.0118 | 0.0019 | 1.08E-09 |
| rs2373124 | 7 | 85830272 | A | -0.0096 | 0.0016 | 4.35E-09 |
| rs1405876 | 7 | 86237581 | T | 0.0109 | 0.0015 | 7.84E-14 |
| rs1021508 | 7 | 92657985 | A | -0.0149 | 0.0014 | 7.03E-26 |
| rs6951996 | 7 | 96646437 | A | -0.0182 | 0.0032 | 1.41E-08 |
| rs6956283 | 7 | 98756597 | T | 0.0098 | 0.0016 | 6.44E-10 |
| rs8192465 | 7 | 99566007 | A | -0.0272 | 0.0049 | 2.69E-08 |
| rs2406253 | 7 | 100077273 | A | 0.0135 | 0.0018 | 3.58E-14 |
| rs401966 | 7 | 101695817 | C | -0.0097 | 0.0014 | 1.72E-11 |
| rs201495 | 7 | 101755522 | T | -0.0112 | 0.0019 | 5.92E-09 |
| rs4730020 | 7 | 104189082 | T | 0.009 | 0.0016 | 1.21E-08 |
| rs2470966 | 7 | 104430889 | C | -0.0103 | 0.0015 | 1.62E-11 |
| rs1075228 | 7 | 104520528 | A | -0.0081 | 0.0014 | 2.18E-08 |
| rs9655780 | 7 | 104667334 | A | -0.0153 | 0.0019 | 3.70E-16 |
| rs567003 | 7 | 105341608 | A | 0.0094 | 0.0015 | 2.27E-10 |
| rs1048045 | 7 | 108283606 | A | 0.0158 | 0.0028 | 2.05E-08 |
| rs1392441 | 7 | 111912738 | A | -0.0193 | 0.003 | 7.71E-11 |
| rs1211363 | 7 | 111996952 | T | 0.0088 | 0.0015 | 2.50E-09 |
| rs1176685 | 7 | 112816099 | T | 0.0234 | 0.0042 | 3.49E-08 |
| rs2100249 | 7 | 113848497 | T | -0.0089 | 0.0015 | 1.91E-09 |
| rs1136151 | 7 | 114519339 | T | -0.0123 | 0.0021 | 2.98E-09 |
| rs6969783 | 7 | 117593308 | A | -0.0098 | 0.0014 | 2.87E-12 |
| rs1714446 | 7 | 122098782 | A | 0.0098 | 0.0015 | 8.74E-11 |
| rs2283076 | 7 | 126478190 | A | 0.0115 | 0.0017 | 1.05E-11 |
| rs6175538 | 7 | 126541317 | T | 0.0125 | 0.0017 | 3.88E-13 |
| rs7799982 | 7 | 127572977 | A | 0.0294 | 0.0039 | 5.08E-14 |
| rs322744 | 7 | 127780401 | T | -0.009 | 0.0016 | 2.04E-08 |
| rs4731413 | 7 | 127836132 | A | 0.0111 | 0.0017 | 2.31E-10 |
| rs339057 | 7 | 128376900 | A | -0.0081 | 0.0014 | 1.26E-08 |
| rs1135204 | 7 | 128402782 | A | 0.0155 | 0.0016 | 1.03E-22 |
| rs7799141 | 7 | 132660489 | A | 0.0099 | 0.0015 | 5.31E-11 |
| rs5735273 | 7 | 133304694 | A | -0.0152 | 0.0017 | 2.51E-18 |
| rs3812281 | 7 | 135082751 | T | 0.0113 | 0.0014 | 2.70E-15 |
| rs6946136 | 7 | 135423424 | A | 0.0081 | 0.0014 | 1.99E-08 |
| rs1025143 | 7 | 137006969 | A | 0.0078 | 0.0014 | 2.65E-08 |
| rs728054 | 7 | 137072364 | A | -0.0128 | 0.0015 | 2.34E-18 |
| rs4726070 | 7 | 151328218 | A | 0.0122 | 0.0014 | 1.36E-17 |
| rs7543427 | 7 | 155424887 | A | 0.0151 | 0.0026 | 8.54E-09 |
| rs7319131 | 8 | 4796094 | A | -0.0094 | 0.0015 | 2.03E-10 |
| rs7016302 | 8 | 4833041 | C | -0.0125 | 0.0019 | 2.26E-11 |
| rs6994287 | 8 | 9340932 | A | 0.0102 | 0.0014 | 1.11E-12 |
| rs5948070 | 8 | 9653130 | C | -0.0123 | 0.0018 | 3.97E-12 |
| rs7655249 | 8 | 12679680 | T | -0.012 | 0.0019 | 1.17E-10 |
| rs2517086 | 8 | 17042416 | C | -0.0088 | 0.0014 | 8.00E-10 |
| rs1326628 | 8 | 17871598 | A | -0.0085 | 0.0015 | 1.75E-08 |

| | | | | | | |
|---|---|---|---|---|---|---|
| rs7816777 | 8 | 19371767 | T | -0.0088 | 0.0015 | 2.35E-09 |
| rs4739235 | 8 | 21287105 | A | 0.0109 | 0.0019 | 5.26E-09 |
| rs7321980 | 8 | 26279173 | A | 0.0106 | 0.0019 | 1.97E-08 |
| rs3480707 | 8 | 28680769 | A | 0.0128 | 0.0019 | 3.29E-11 |
| rs2725370 | 8 | 30852826 | T | -0.0141 | 0.0015 | 4.96E-20 |
| rs6250607 | 8 | 30906089 | T | 0.0091 | 0.0015 | 8.64E-10 |
| rs6250610 | 8 | 31020008 | A | -0.0098 | 0.0018 | 3.93E-08 |
| rs763553 | 8 | 31445496 | A | 0.0087 | 0.0014 | 6.41E-10 |
| rs2980813 | 8 | 40050010 | A | 0.0088 | 0.0014 | 4.28E-10 |
| rs7823700 | 8 | 42363231 | T | 0.0132 | 0.0024 | 4.00E-08 |
| rs2923424 | 8 | 42382222 | A | 0.012 | 0.0014 | 6.31E-17 |
| rs2929032 | 8 | 56371745 | A | -0.0081 | 0.0014 | 8.40E-09 |
| rs1866823 | 8 | 57436577 | A | 0.0107 | 0.0014 | 3.09E-14 |
| rs6472208 | 8 | 66474730 | T | -0.0091 | 0.0014 | 2.00E-10 |
| rs2926702 | 8 | 71167994 | T | -0.0124 | 0.0021 | 4.91E-09 |
| rs5609937 | 8 | 77372988 | T | 0.012 | 0.0017 | 3.10E-13 |
| rs8003790 | 8 | 87170913 | T | -0.0107 | 0.0019 | 2.16E-08 |
| rs1768071 | 8 | 87300775 | T | -0.0092 | 0.0014 | 8.63E-11 |
| rs4320563 | 8 | 87769961 | A | 0.0109 | 0.0014 | 2.24E-14 |
| rs2740795 | 8 | 91927455 | A | 0.0098 | 0.0016 | 7.71E-10 |
| rs1905616 | 8 | 93235675 | A | 0.0083 | 0.0015 | 2.74E-08 |
| rs3560643 | 8 | 93327532 | A | 0.0105 | 0.0016 | 3.98E-11 |
| rs4735297 | 8 | 95589340 | A | -0.0086 | 0.0015 | 1.62E-08 |
| rs1693584 | 8 | 101748917 | T | -0.0087 | 0.0015 | 8.79E-09 |
| rs7012546 | 8 | 105067737 | T | 0.0092 | 0.0014 | 1.25E-10 |
| rs7267205 | 8 | 115798424 | A | -0.0101 | 0.0018 | 4.87E-08 |
| rs6469654 | 8 | 117632965 | C | -0.0103 | 0.0017 | 1.10E-09 |
| rs4298514 | 8 | 118932887 | T | 0.0104 | 0.0015 | 1.07E-11 |
| rs1328156 | 8 | 119483316 | A | 0.0088 | 0.0014 | 4.02E-10 |
| rs1485300 | 8 | 120082253 | C | 0.0098 | 0.0015 | 1.14E-10 |
| rs1326177 | 8 | 120187090 | C | 0.0122 | 0.0019 | 3.12E-10 |
| rs401526 | 8 | 130929166 | T | -0.0104 | 0.0014 | 1.15E-13 |
| rs7446262 | 8 | 133732856 | T | 0.0125 | 0.002 | 1.21E-09 |
| rs7454533 | 8 | 135440608 | A | -0.0158 | 0.0022 | 9.06E-13 |
| rs2977464 | 8 | 141545193 | T | 0.0102 | 0.0018 | 1.46E-08 |
| rs1178002 | 8 | 141992778 | T | 0.0085 | 0.0014 | 1.17E-09 |
| rs746839 | 8 | 142617261 | C | 0.0135 | 0.0015 | 1.19E-20 |
| rs6989141 | 8 | 142681460 | A | 0.01 | 0.0017 | 8.40E-09 |
| rs902820 | 8 | 143099872 | A | -0.0096 | 0.0014 | 1.28E-11 |
| rs1009807 | 8 | 143309504 | A | -0.0091 | 0.0014 | 1.03E-10 |
| rs1443367 | 8 | 143356114 | A | 0.0388 | 0.0053 | 2.56E-13 |
| rs7770262 | 8 | 143367857 | A | -0.0213 | 0.0029 | 3.62E-13 |
| rs7460106 | 8 | 143534777 | T | -0.0123 | 0.0017 | 2.06E-13 |
| rs6667163 | 8 | 143680772 | T | -0.0133 | 0.0021 | 2.24E-10 |
| rs2976397 | 8 | 143764613 | T | 0.0081 | 0.0014 | 1.30E-08 |
| rs1177421 | 8 | 145686505 | T | 0.0128 | 0.0014 | 1.40E-19 |
| rs4741571 | 9 | 1667814 | A | -0.0102 | 0.0015 | 8.09E-12 |

| | | | | | | |
|---|---|---|---|---|---|---|
| rs7849487 | 9 | 1757274 | T | -0.0167 | 0.0015 | 1.25E-29 |
| rs3847228 | 9 | 1798805 | T | 0.0083 | 0.0014 | 7.47E-09 |
| rs1097425 | 9 | 3952892 | A | -0.0093 | 0.0015 | 4.12E-10 |
| rs7364371 | 9 | 4163612 | T | -0.0165 | 0.0027 | 1.24E-09 |
| rs7364845 | 9 | 14077378 | T | -0.0183 | 0.0026 | 1.99E-12 |
| rs1081009 | 9 | 14161927 | A | -0.0146 | 0.0016 | 1.91E-20 |
| rs1223801 | 9 | 14210897 | T | -0.0141 | 0.0026 | 4.67E-08 |
| rs1081014 | 9 | 14430692 | A | 0.0078 | 0.0014 | 4.00E-08 |
| rs7875078 | 9 | 14494845 | A | -0.0078 | 0.0014 | 2.72E-08 |
| rs7849480 | 9 | 14648130 | A | -0.0084 | 0.0014 | 3.54E-09 |
| rs1329634 | 9 | 14791348 | T | 0.009 | 0.0015 | 5.35E-10 |
| rs7345793 | 9 | 23233667 | A | 0.0144 | 0.0025 | 1.11E-08 |
| rs7029718 | 9 | 23358495 | A | 0.0244 | 0.0014 | 7.82E-65 |
| rs1172734 | 9 | 23385738 | T | -0.0211 | 0.0034 | 8.59E-10 |
| rs1328451 | 9 | 23423785 | C | -0.0082 | 0.0014 | 6.78E-09 |
| rs4977885 | 9 | 23687983 | A | -0.0084 | 0.0014 | 6.78E-09 |
| rs1329044 | 9 | 23747791 | T | 0.0134 | 0.0019 | 6.28E-13 |
| rs2805064 | 9 | 24530615 | C | 0.0093 | 0.0016 | 3.54E-09 |
| rs1758747 | 9 | 25300643 | A | 0.0092 | 0.0015 | 1.64E-09 |
| rs7041702 | 9 | 33053430 | A | -0.0111 | 0.0016 | 4.53E-12 |
| rs1113894 | 9 | 72110562 | T | 0.0105 | 0.0016 | 2.36E-11 |
| rs4434676 | 9 | 73032476 | A | -0.0089 | 0.0014 | 2.83E-10 |
| rs9886703 | 9 | 82246351 | A | -0.0136 | 0.0019 | 4.46E-13 |
| rs4877516 | 9 | 82442731 | A | -0.0101 | 0.0014 | 7.78E-13 |
| rs7035315 | 9 | 83231511 | A | -0.009 | 0.0015 | 5.85E-10 |
| rs7855503 | 9 | 86381638 | C | 0.0088 | 0.0015 | 2.39E-09 |
| rs995698 | 9 | 87997869 | A | -0.0099 | 0.0014 | 1.53E-12 |
| rs7040995 | 9 | 92226172 | C | 0.0097 | 0.0014 | 5.39E-12 |
| rs1076120 | 9 | 96019970 | T | 0.0088 | 0.0014 | 4.28E-10 |
| rs1113705 | 9 | 96203756 | T | 0.0151 | 0.0027 | 1.61E-08 |
| rs2989751 | 9 | 96366647 | A | -0.0092 | 0.0017 | 2.89E-08 |
| rs1076125 | 9 | 96421546 | A | 0.0113 | 0.0015 | 2.52E-14 |
| rs1118210 | 9 | 99084793 | T | 0.0124 | 0.0019 | 1.58E-10 |
| rs1178901 | 9 | 109677417 | T | 0.0106 | 0.0016 | 1.47E-10 |
| rs1097961 | 9 | 111687584 | T | -0.0117 | 0.0015 | 1.25E-15 |
| rs1012266 | 9 | 116555771 | A | -0.0079 | 0.0014 | 3.00E-08 |
| rs1098332 | 9 | 119485337 | A | 0.0084 | 0.0015 | 3.55E-08 |
| rs7623588 | 9 | 121096681 | A | 0.0241 | 0.0039 | 4.52E-10 |
| rs7030373 | 9 | 121161926 | A | -0.0098 | 0.0017 | 1.49E-08 |
| rs1098444 | 9 | 121980586 | A | 0.0115 | 0.0014 | 2.58E-16 |
| rs1009470 | 9 | 122289187 | T | 0.0086 | 0.0014 | 1.34E-09 |
| rs2416759 | 9 | 123358262 | A | -0.0093 | 0.0015 | 1.45E-09 |
| rs1098540 | 9 | 124578782 | T | 0.0089 | 0.0016 | 4.47E-08 |
| rs1237594 | 9 | 124617900 | T | -0.0138 | 0.0014 | 1.61E-22 |
| rs2416845 | 9 | 124755432 | T | 0.0263 | 0.0048 | 4.19E-08 |
| rs1467737 | 9 | 124982500 | T | -0.0084 | 0.0014 | 3.52E-09 |
| rs1234254 | 9 | 126329270 | A | -0.0128 | 0.0017 | 1.11E-14 |

| | | | | | | | |
|---|---|---|---|---|---|---|---|
| rs631287 | 9 | 128412676 | A | 0.0084 | 0.0014 | 2.52E-09 | |
| rs2039204 | 9 | 130371107 | A | 0.0082 | 0.0014 | 5.23E-09 | |
| rs913509 | 9 | 134737751 | A | 0.0096 | 0.0015 | 7.42E-11 | |
| rs9411331 | 9 | 134883419 | A | 0.0145 | 0.0015 | 7.63E-22 | |
| rs1079390 | 9 | 134976652 | A | 0.0126 | 0.0022 | 1.37E-08 | |
| rs1200515 | 9 | 135030649 | A | 0.0087 | 0.0015 | 3.73E-09 | |
| rs1268277 | 9 | 135490491 | T | -0.0119 | 0.0017 | 1.26E-12 | |
| rs214626 | 9 | 135718584 | A | 0.0101 | 0.0018 | 1.58E-08 | |
| rs7358158 | 9 | 140251458 | A | -0.0181 | 0.0022 | 5.78E-16 | Not available |
| rs4881269 | 10 | 4031519 | A | 0.01 | 0.0014 | 3.97E-12 | |
| rs7894722 | 10 | 9972046 | T | 0.0085 | 0.0014 | 5.41E-09 | |
| rs1714899 | 10 | 10909401 | A | 0.0101 | 0.0017 | 6.98E-09 | |
| rs1079583 | 10 | 10943379 | T | -0.0099 | 0.0017 | 6.98E-09 | |
| rs1747714 | 10 | 11161316 | T | 0.0106 | 0.0014 | 3.82E-14 | |
| rs1075226 | 10 | 12395100 | T | 0.0102 | 0.0014 | 1.28E-12 | |
| rs7905192 | 10 | 12711949 | T | 0.0084 | 0.0014 | 3.14E-09 | |
| rs2183271 | 10 | 21957229 | T | 0.0084 | 0.0015 | 1.91E-08 | |
| rs1159876 | 10 | 23953725 | A | -0.0106 | 0.0018 | 1.73E-09 | |
| rs806816 | 10 | 32285942 | A | 0.0101 | 0.0016 | 4.80E-10 | |
| rs2007655 | 10 | 34586689 | T | 0.0086 | 0.0014 | 8.64E-10 | |
| rs1100346 | 10 | 51801793 | T | 0.0091 | 0.0015 | 5.58E-10 | |
| rs2588959 | 10 | 63598957 | T | -0.008 | 0.0014 | 9.88E-09 | |
| rs6185333 | 10 | 64884737 | T | -0.0109 | 0.0018 | 3.29E-09 | |
| rs7924036 | 10 | 65191645 | T | 0.0133 | 0.0014 | 2.55E-21 | |
| rs1159187 | 10 | 65494036 | T | 0.0098 | 0.0017 | 2.25E-08 | |
| rs1099563 | 10 | 65573195 | T | 0.0078 | 0.0014 | 4.26E-08 | |
| rs1099616 | 10 | 66813928 | C | 0.0113 | 0.002 | 7.47E-09 | |
| rs1235937 | 10 | 67767951 | T | -0.0092 | 0.0015 | 6.33E-10 | |
| rs1050925 | 10 | 67828436 | T | -0.0093 | 0.0017 | 3.61E-08 | |
| rs7920624 | 10 | 67963186 | A | 0.012 | 0.0014 | 1.06E-17 | |
| rs4469771 | 10 | 68192714 | T | 0.0101 | 0.0015 | 1.88E-11 | |
| rs6480234 | 10 | 68636894 | T | -0.0084 | 0.0014 | 1.94E-09 | |
| rs2657283 | 10 | 76920199 | T | 0.0085 | 0.0014 | 2.08E-09 | |
| rs7910403 | 10 | 87039251 | T | 0.0122 | 0.0018 | 5.08E-12 | |
| rs1426619 | 10 | 90091540 | T | 0.0096 | 0.0014 | 1.40E-11 | |
| rs482787 | 10 | 99767024 | T | 0.01 | 0.0015 | 1.95E-11 | |
| rs1788280 | 10 | 101983413 | A | 0.0084 | 0.0014 | 3.00E-09 | |
| rs1119095 | 10 | 103112165 | T | -0.0083 | 0.0015 | 1.74E-08 | |
| rs1159638 | 10 | 103606122 | T | 0.0258 | 0.0046 | 2.20E-08 | |
| rs1882515 | 10 | 103744305 | A | 0.0428 | 0.007 | 1.07E-09 | Excluded Low MAF |
| rs7334483 | 10 | 103816828 | A | 0.017 | 0.0014 | 1.00E-32 | |
| rs1119123 | 10 | 103949787 | T | 0.0132 | 0.0024 | 1.96E-08 | |
| rs4919624 | 10 | 104021085 | A | 0.0193 | 0.0018 | 6.52E-28 | |
| rs1159229 | 10 | 104225941 | A | 0.0097 | 0.0018 | 4.19E-08 | |
| rs1277457 | 10 | 104959487 | T | 0.0142 | 0.002 | 8.69E-13 | |
| rs1074884 | 10 | 104996309 | A | -0.0111 | 0.002 | 1.58E-08 | |
| rs7695767 | 10 | 105030157 | T | -0.0297 | 0.0054 | 4.72E-08 | |

| | | | | | | |
|---|---|---|---|---|---|---|
| rs3781339 | 10 | 105428152 | T | -0.0104 | 0.0018 | 7.55E-09 |
| rs1159923 | 10 | 106454672 | T | -0.0106 | 0.0014 | 9.94E-14 |
| rs7914674 | 10 | 106738675 | T | -0.0139 | 0.0022 | 1.94E-10 |
| rs790647 | 10 | 106776484 | A | -0.0145 | 0.0017 | 3.81E-18 |
| rs1276459 | 10 | 107491895 | C | 0.0202 | 0.0029 | 6.00E-12 |
| rs1276172 | 10 | 107619932 | T | -0.0098 | 0.0016 | 1.87E-09 |
| rs6072648 | 10 | 110882068 | T | -0.009 | 0.0016 | 2.35E-08 |
| rs7899270 | 10 | 111258134 | A | -0.0084 | 0.0015 | 1.74E-08 |
| rs1257154 | 10 | 111770013 | A | 0.0151 | 0.002 | 1.45E-13 |
| rs1088601 | 10 | 118550021 | C | 0.0078 | 0.0014 | 4.24E-08 |
| rs1865955 | 10 | 118837990 | T | 0.0122 | 0.0019 | 4.06E-11 |
| rs4384309 | 10 | 133110596 | A | 0.0102 | 0.0014 | 4.89E-13 |
| rs1276176 | 10 | 133775375 | T | 0.0153 | 0.0017 | 4.79E-20 |
| rs7084508 | 10 | 133932946 | T | -0.0085 | 0.0015 | 8.79E-09 |
| rs1276518 | 10 | 134977077 | A | -0.0088 | 0.0016 | 2.96E-08 |
| rs1083165 | 11 | 11660815 | T | 0.009 | 0.0015 | 3.10E-09 |
| rs4757957 | 11 | 12881398 | C | 0.0133 | 0.0015 | 1.79E-18 |
| rs1102376 | 11 | 15910939 | A | 0.0091 | 0.0015 | 5.32E-10 |
| rs1880088 | 11 | 20959185 | A | -0.0097 | 0.0016 | 1.11E-09 |
| rs1278931 | 11 | 25058937 | T | 0.0092 | 0.0014 | 6.79E-11 |
| rs1103008 | 11 | 27643725 | T | 0.0106 | 0.0018 | 4.07E-09 |
| rs1103010 | 11 | 27681596 | C | 0.0101 | 0.0016 | 4.60E-10 |
| rs7288376 | 11 | 29331969 | A | 0.0101 | 0.0018 | 8.64E-09 |
| rs322614 | 11 | 29755547 | A | -0.0087 | 0.0014 | 1.43E-09 |
| rs795230 | 11 | 30774525 | T | 0.0081 | 0.0014 | 9.56E-09 |
| rs2199409 | 11 | 39915234 | T | 0.0103 | 0.0017 | 3.14E-09 |
| rs1044428 | 11 | 41155897 | T | 0.01 | 0.0018 | 1.47E-08 |
| rs1074259 | 11 | 41548956 | A | -0.0084 | 0.0014 | 4.26E-09 |
| rs7972801 | 11 | 46062245 | A | -0.0134 | 0.002 | 2.06E-11 |
| rs1279019 | 11 | 57497847 | T | -0.0101 | 0.0015 | 3.29E-11 |
| rs7503301 | 11 | 61347033 | C | -0.0258 | 0.0038 | 1.38E-11 |
| rs7931563 | 11 | 61446425 | T | -0.0109 | 0.0014 | 5.15E-14 |
| rs628993 | 11 | 61539691 | A | 0.0128 | 0.0023 | 3.83E-08 |
| rs1180930 | 11 | 62413673 | T | 0.0146 | 0.0021 | 3.78E-12 |
| rs5877994 | 11 | 64007159 | A | 0.011 | 0.0019 | 3.84E-09 |
| rs3511150 | 11 | 65334712 | C | -0.0175 | 0.003 | 7.90E-09 |
| rs585557 | 11 | 65663547 | A | 0.0128 | 0.0018 | 3.25E-13 |
| rs479018 | 11 | 66060546 | A | 0.0109 | 0.0015 | 3.10E-13 |
| rs7687866 | 11 | 66092567 | C | 0.0141 | 0.0017 | 3.10E-17 |
| rs7351880 | 11 | 68282986 | C | -0.0128 | 0.0021 | 1.15E-09 |
| rs7296216 | 11 | 72365669 | T | -0.0174 | 0.0019 | 1.26E-19 |
| rs1089928 | 11 | 76504698 | A | 0.014 | 0.0017 | 1.05E-16 |
| rs7127580 | 11 | 79144856 | C | -0.015 | 0.0023 | 3.12E-11 |
| rs1880692 | 11 | 80338069 | A | 0.0086 | 0.0014 | 1.09E-09 |
| rs580652 | 11 | 84904742 | T | -0.0147 | 0.0024 | 4.65E-10 |
| rs9666728 | 11 | 87907712 | T | -0.0082 | 0.0014 | 7.10E-09 |
| rs1792602 | 11 | 90439633 | A | -0.0097 | 0.0014 | 1.90E-11 |

| | | | | | | | |
|---|---|---|---|---|---|---|---|
| rs1083085 | 11 | 91896638 | T | 0.0092 | 0.0014 | 5.31E-11 | |
| rs488476 | 11 | 95543715 | C | -0.0113 | 0.0015 | 8.29E-15 | |
| rs1102143 | 11 | 95837674 | A | 0.0103 | 0.0015 | 1.25E-12 | |
| rs1301838 | 11 | 99670514 | T | -0.0096 | 0.0015 | 1.29E-10 | |
| rs7248602 | 11 | 107167486 | T | -0.0104 | 0.0016 | 1.61E-10 | Not available |
| rs2212430 | 11 | 109289538 | T | -0.01 | 0.0016 | 1.32E-10 | |
| rs1121348 | 11 | 110435636 | A | -0.0106 | 0.0019 | 2.68E-08 | |
| rs1753605 | 11 | 110957335 | C | -0.0108 | 0.0019 | 6.45E-09 | |
| rs1756597 | 11 | 111586950 | A | -0.0106 | 0.0014 | 5.54E-14 | |
| rs7928017 | 11 | 113448762 | A | 0.0096 | 0.0014 | 1.43E-11 | |
| rs648044 | 11 | 114030799 | A | 0.0083 | 0.0015 | 8.89E-09 | |
| rs2126069 | 11 | 115473374 | T | 0.008 | 0.0014 | 1.11E-08 | |
| rs1228507 | 11 | 116762028 | A | 0.0161 | 0.0023 | 5.11E-12 | |
| rs4938815 | 11 | 120441647 | T | 0.0085 | 0.0015 | 2.99E-08 | |
| rs1089280 | 11 | 121998254 | T | 0.0109 | 0.0014 | 1.09E-14 | |
| rs4127499 | 11 | 122176383 | A | 0.0087 | 0.0015 | 3.56E-09 | |
| rs1257428 | 11 | 131205421 | A | -0.008 | 0.0015 | 3.04E-08 | |
| rs7108020 | 11 | 131289820 | A | 0.0111 | 0.0015 | 3.91E-14 | |
| rs563954 | 11 | 132391656 | A | -0.0081 | 0.0014 | 1.34E-08 | |
| rs9633970 | 11 | 132502212 | T | 0.0091 | 0.0016 | 2.81E-08 | |
| rs6766127 | 11 | 132662671 | A | 0.0134 | 0.0025 | 4.40E-08 | |
| rs1075053 | 11 | 133548873 | A | -0.0095 | 0.0015 | 1.11E-10 | |
| rs1176779 | 11 | 133772904 | A | -0.018 | 0.0032 | 2.03E-08 | |
| rs7303429 | 11 | 133822133 | A | 0.0099 | 0.0018 | 2.11E-08 | |
| rs1227343 | 11 | 133825855 | A | -0.0132 | 0.0017 | 4.72E-14 | |
| rs5720426 | 11 | 133844024 | A | 0.0114 | 0.002 | 9.94E-09 | |
| rs2368831 | 12 | 188285 | T | -0.0092 | 0.0014 | 1.63E-10 | |
| rs4766424 | 12 | 1954096 | C | 0.0142 | 0.0021 | 2.47E-11 | |
| rs1084505 | 12 | 10291446 | A | 0.0083 | 0.0014 | 3.75E-09 | |
| rs1077264 | 12 | 13417617 | C | 0.0142 | 0.0022 | 1.69E-10 | |
| rs1861786 | 12 | 14000467 | A | -0.0081 | 0.0014 | 2.39E-08 | |
| rs7974852 | 12 | 14511679 | A | 0.0117 | 0.0014 | 1.14E-16 | |
| rs1117152 | 12 | 15438032 | T | 0.0106 | 0.0015 | 4.47E-12 | |
| rs2160514 | 12 | 16756508 | A | -0.0094 | 0.0014 | 2.54E-11 | |
| rs1603460 | 12 | 23092626 | T | 0.009 | 0.0014 | 2.20E-10 | |
| rs7965154 | 12 | 23697725 | A | -0.0085 | 0.0015 | 2.33E-08 | |
| rs1995181 | 12 | 24195048 | A | -0.0078 | 0.0014 | 3.61E-08 | |
| rs1173980 | 12 | 26523628 | C | -0.0141 | 0.0025 | 1.16E-08 | |
| rs1161343 | 12 | 26684343 | A | -0.0094 | 0.0016 | 6.71E-09 | |
| rs7972246 | 12 | 27209328 | T | 0.011 | 0.0015 | 1.01E-13 | |
| rs1084417 | 12 | 32469601 | A | 0.0104 | 0.0017 | 5.03E-10 | |
| rs6779023 | 12 | 46079172 | A | 0.0101 | 0.0017 | 6.90E-09 | |
| rs1078324 | 12 | 48653003 | A | -0.0092 | 0.0014 | 5.80E-11 | |
| rs1054442 | 12 | 49389320 | A | -0.0136 | 0.0015 | 1.10E-20 | |
| rs8025797 | 12 | 49402562 | T | -0.0252 | 0.004 | 3.31E-10 | |
| rs1178957 | 12 | 49729637 | T | 0.0362 | 0.0064 | 1.51E-08 | |
| rs933738 | 12 | 49943122 | A | -0.011 | 0.0018 | 2.08E-09 | |

| | | | | | | |
|---|---|---|---|---|---|---|
| rs1711010 | 12 | 54668908 | T | -0.0093 | 0.0014 | 1.47E-10 |
| rs8022341 | 12 | 56367966 | T | -0.0118 | 0.002 | 8.64E-09 |
| rs1689510 | 12 | 56396768 | C | 0.0194 | 0.0015 | 1.35E-38 |
| rs7616722 | 12 | 56507702 | T | -0.0187 | 0.0033 | 1.88E-08 |
| rs3751331 | 12 | 58290278 | A | -0.0097 | 0.0014 | 1.75E-11 |
| rs1087728 | 12 | 59837734 | T | 0.0085 | 0.0014 | 2.86E-09 |
| rs5981332 | 12 | 60879673 | T | 0.0097 | 0.0018 | 4.15E-08 |
| rs1146079 | 12 | 62485472 | C | -0.0087 | 0.0016 | 4.24E-08 |
| rs1117439 | 12 | 62646556 | A | 0.0111 | 0.0018 | 2.42E-10 |
| rs710629 | 12 | 67697856 | A | 0.0082 | 0.0015 | 2.09E-08 |
| rs1087967 | 12 | 74152402 | T | -0.0083 | 0.0014 | 5.29E-09 |
| rs1711373 | 12 | 74936526 | A | -0.017 | 0.0027 | 4.10E-10 |
| rs5766153 | 12 | 78629917 | T | -0.0114 | 0.0021 | 3.24E-08 |
| rs1245829 | 12 | 79648170 | A | -0.0099 | 0.0014 | 3.29E-12 |
| rs1111505 | 12 | 82258775 | A | 0.0187 | 0.0023 | 1.67E-15 |
| rs717996 | 12 | 84133820 | T | -0.0122 | 0.0014 | 1.89E-17 |
| rs1427829 | 12 | 89760744 | A | -0.0083 | 0.0014 | 3.29E-09 |
| rs7967550 | 12 | 92145307 | A | -0.0086 | 0.0014 | 1.79E-09 |
| rs2216144 | 12 | 97666478 | T | 0.0098 | 0.0014 | 2.58E-12 |
| rs1012888 | 12 | 97799524 | A | -0.0083 | 0.0015 | 4.04E-08 |
| rs1230418 | 12 | 97981272 | A | 0.0111 | 0.0019 | 8.17E-09 |
| rs2131167 | 12 | 99633189 | A | 0.008 | 0.0014 | 1.52E-08 |
| rs7958371 | 12 | 103456084 | A | -0.0085 | 0.0015 | 5.99E-09 |
| rs1196760 | 12 | 105606068 | C | 0.0139 | 0.0025 | 2.00E-08 |
| rs2866988 | 12 | 106722701 | A | -0.0084 | 0.0015 | 8.31E-09 |
| rs7575684 | 12 | 110111274 | A | 0.0288 | 0.0049 | 5.79E-09 |
| rs7977614 | 12 | 110115286 | A | -0.0134 | 0.0016 | 6.84E-17 |
| rs3809169 | 12 | 113911532 | T | 0.014 | 0.0025 | 3.85E-08 |
| rs7340529 | 12 | 117522917 | A | 0.0113 | 0.002 | 7.42E-09 |
| rs1671770 | 12 | 120946568 | A | 0.0125 | 0.0018 | 9.23E-12 |
| rs1718188 | 12 | 121343893 | A | 0.0096 | 0.0014 | 9.05E-12 |
| rs2954114 | 12 | 122169884 | A | -0.0081 | 0.0014 | 9.40E-09 |
| rs1281058 | 12 | 122675576 | T | 0.0094 | 0.0015 | 3.70E-10 |
| rs1077320 | 12 | 122970156 | T | -0.011 | 0.0016 | 8.09E-12 |
| rs7844061 | 12 | 123144189 | A | -0.0147 | 0.0024 | 5.71E-10 |
| rs1077300 | 12 | 123746961 | A | 0.0185 | 0.0016 | 2.50E-30 |
| rs1107871 | 12 | 132240659 | A | -0.0083 | 0.0014 | 4.18E-09 |
| rs6490618 | 13 | 21338343 | T | 0.0088 | 0.0015 | 3.48E-09 |
| rs4941735 | 13 | 31641248 | T | 0.0092 | 0.0014 | 8.63E-11 |
| rs9529146 | 13 | 31785404 | T | 0.0127 | 0.0017 | 3.48E-14 |
| rs936496 | 13 | 36100541 | A | -0.0079 | 0.0014 | 4.38E-08 |
| rs9545395 | 13 | 36370970 | T | 0.0138 | 0.0021 | 3.81E-11 |
| rs9568798 | 13 | 53614554 | T | -0.0093 | 0.0016 | 2.30E-09 |
| rs9536462 | 13 | 54124650 | A | 0.0133 | 0.002 | 2.10E-11 |
| rs9563168 | 13 | 54247827 | A | 0.0101 | 0.0017 | 4.07E-09 |
| rs1115301 | 13 | 54342123 | T | 0.0269 | 0.0048 | 1.62E-08 |
| rs1287533 | 13 | 55722033 | A | -0.0108 | 0.0015 | 1.71E-13 |

| | | | | | | |
|---|---|---|---|---|---|---|
| rs1137205 | 13 | 57477706 | A | 0.0164 | 0.0029 | 2.34E-08 |
| rs9527662 | 13 | 58020265 | A | 0.0094 | 0.0014 | 3.37E-11 |
| rs7737094 | 13 | 58037032 | A | -0.0147 | 0.0027 | 3.65E-08 |
| rs1427471 | 13 | 58271461 | A | 0.0329 | 0.0052 | 2.84E-10 |
| rs1334297 | 13 | 58335375 | A | 0.0257 | 0.0016 | 2.53E-58 |
| rs1169673 | 13 | 58466434 | A | 0.0315 | 0.0056 | 1.53E-08 |
| rs6195817 | 13 | 58680250 | A | -0.0198 | 0.0034 | 8.22E-09 |
| rs7323027 | 13 | 58693790 | A | 0.013 | 0.0015 | 3.91E-19 |
| rs1864567 | 13 | 58698035 | A | 0.0455 | 0.0079 | 7.26E-09 |
| rs2321157 | 13 | 58736901 | A | -0.0081 | 0.0014 | 7.64E-09 |
| rs1719041 | 13 | 58746132 | T | -0.0213 | 0.0031 | 8.42E-12 |
| rs9527905 | 13 | 59403033 | A | -0.0094 | 0.0014 | 4.11E-11 |
| rs9597907 | 13 | 59496462 | T | 0.0152 | 0.0027 | 1.56E-08 |
| rs341504 | 13 | 60421219 | A | 0.0099 | 0.0015 | 2.22E-11 |
| rs4497562 | 13 | 62612604 | A | 0.0112 | 0.0016 | 1.05E-12 |
| rs9540718 | 13 | 66920915 | A | -0.0089 | 0.0014 | 1.88E-10 |
| rs7687659 | 13 | 67141932 | A | 0.0126 | 0.002 | 5.82E-10 |
| rs7321274 | 13 | 69146186 | A | 0.0114 | 0.0018 | 8.21E-11 |
| rs716513 | 13 | 69488908 | A | 0.0084 | 0.0014 | 2.83E-09 |
| rs2478208 | 13 | 81471113 | C | -0.0103 | 0.0014 | 2.89E-13 |
| rs7326331 | 13 | 91615838 | A | -0.0128 | 0.0016 | 3.42E-16 |
| rs1162035 | 13 | 92044587 | A | 0.0182 | 0.0025 | 1.09E-13 |
| rs9513416 | 13 | 99055774 | A | -0.0116 | 0.0019 | 2.18E-09 |
| rs9556958 | 13 | 99100046 | T | -0.011 | 0.0014 | 5.96E-15 |
| rs7317761 | 13 | 100753796 | C | 0.0109 | 0.0016 | 7.77E-12 |
| rs9513754 | 13 | 101043420 | T | 0.0096 | 0.0016 | 6.64E-10 |
| rs9300612 | 13 | 101283219 | T | 0.0084 | 0.0015 | 3.07E-08 |
| rs1014552 | 14 | 21930932 | T | -0.0122 | 0.0018 | 4.78E-12 |
| rs1115793 | 14 | 23403193 | A | -0.0139 | 0.0014 | 3.86E-22 |
| rs1162328 | 14 | 24557642 | T | -0.0135 | 0.0021 | 5.58E-11 |
| rs178183 | 14 | 26979412 | T | 0.0145 | 0.0016 | 4.81E-19 |
| rs176218 | 14 | 29600506 | T | 0.02 | 0.0018 | 1.10E-29 |
| rs7267145 | 14 | 29662737 | A | -0.0174 | 0.002 | 1.27E-17 |
| rs7147473 | 14 | 30608538 | A | -0.0103 | 0.0015 | 6.33E-12 |
| rs8016504 | 14 | 33308021 | A | -0.0087 | 0.0014 | 6.84E-10 |
| rs1289104 | 14 | 34018986 | T | 0.0092 | 0.0014 | 1.02E-10 |
| rs4981245 | 14 | 35088635 | A | 0.008 | 0.0014 | 1.58E-08 |
| rs1007731 | 14 | 37054590 | A | -0.0143 | 0.0022 | 1.05E-10 |
| rs8009933 | 14 | 39600212 | A | 0.0088 | 0.0015 | 4.72E-09 |
| rs1123285 | 14 | 57274519 | C | -0.0094 | 0.0015 | 2.38E-10 |
| rs6199766 | 14 | 57280046 | T | -0.0144 | 0.002 | 2.24E-13 |
| rs198262 | 14 | 57283283 | T | -0.0223 | 0.0035 | 1.32E-10 |
| rs1289119 | 14 | 58100283 | C | 0.008 | 0.0014 | 1.21E-08 |
| rs1051860 | 14 | 58838668 | A | -0.0079 | 0.0014 | 3.29E-08 |
| rs4899012 | 14 | 61003889 | C | -0.0103 | 0.0014 | 8.06E-13 |
| rs6573552 | 14 | 64750233 | T | -0.009 | 0.0014 | 1.17E-10 |
| rs6573559 | 14 | 64923166 | T | 0.0105 | 0.0015 | 3.11E-12 |

| | | | | | | |
|---|---|---|---|---|---|---|
| rs1115880 | 14 | 69747046 | A | -0.0091 | 0.0014 | 7.15E-11 |
| rs1288861 | 14 | 72435705 | T | 0.012 | 0.0018 | 9.42E-12 |
| rs7158218 | 14 | 73934880 | A | -0.009 | 0.0015 | 5.63E-09 |
| rs2358628 | 14 | 74631757 | A | -0.0091 | 0.0015 | 1.78E-09 |
| rs730384 | 14 | 74889870 | A | 0.0101 | 0.0014 | 9.38E-13 |
| rs1779549 | 14 | 84640016 | A | 0.0081 | 0.0014 | 8.45E-09 |
| rs7146625 | 14 | 84788153 | A | 0.0091 | 0.0016 | 7.55E-09 |
| rs1431483 | 14 | 84896824 | T | -0.0211 | 0.0039 | 4.62E-08 |
| rs2998299 | 14 | 84997363 | T | -0.0136 | 0.0017 | 1.51E-15 |
| rs7269447 | 14 | 85112638 | T | 0.0127 | 0.002 | 4.94E-10 |
| rs1184578 | 14 | 89276431 | T | -0.0079 | 0.0014 | 2.43E-08 |
| rs4904523 | 14 | 89723630 | A | -0.0082 | 0.0014 | 6.27E-09 |
| rs1952183 | 14 | 89844111 | A | -0.0089 | 0.0014 | 2.76E-10 |
| rs736281 | 14 | 94287830 | T | 0.0086 | 0.0014 | 2.49E-09 |
| rs2496482 | 14 | 99750164 | T | 0.0121 | 0.0015 | 9.20E-17 |
| rs1243168 | 14 | 101539999 | T | -0.0093 | 0.0015 | 1.73E-10 |
| rs8008382 | 14 | 104054425 | T | -0.0106 | 0.0015 | 4.07E-12 |
| rs7167688 | 15 | 26813137 | T | -0.0094 | 0.0014 | 1.71E-11 |
| rs891793 | 15 | 27256021 | C | -0.0099 | 0.0014 | 1.87E-12 |
| rs4423373 | 15 | 27988340 | A | -0.0087 | 0.0014 | 6.00E-10 |
| rs1177994 | 15 | 34659517 | C | 0.0106 | 0.0016 | 1.19E-11 |
| rs1696627 | 15 | 38423730 | T | 0.0092 | 0.0016 | 4.21E-09 |
| rs6493265 | 15 | 47513253 | T | -0.0122 | 0.0014 | 2.26E-17 |
| rs6493275 | 15 | 47676579 | A | 0.0131 | 0.0017 | 2.98E-14 |
| rs1656614 | 15 | 47800513 | C | 0.0122 | 0.0015 | 6.91E-16 |
| rs1898111 | 15 | 47892298 | A | 0.0101 | 0.0018 | 4.72E-08 |
| rs2414072 | 15 | 50850562 | A | -0.0083 | 0.0014 | 4.45E-09 |
| rs6201821 | 15 | 51387404 | T | 0.0113 | 0.0019 | 4.83E-09 |
| rs1243817 | 15 | 55951432 | A | 0.0093 | 0.0015 | 1.48E-10 |
| rs1529597 | 15 | 57190064 | A | 0.0223 | 0.0038 | 3.36E-09 |
| rs2431023 | 15 | 57553832 | A | 0.0093 | 0.0014 | 5.76E-11 |
| rs7171405 | 15 | 61466893 | A | -0.0101 | 0.0016 | 7.53E-10 |
| rs3426265 | 15 | 63849287 | T | -0.024 | 0.0038 | 2.84E-10 |
| rs4984613 | 15 | 65023232 | T | 0.0174 | 0.0028 | 7.76E-10 |
| rs1259164 | 15 | 65959729 | T | -0.0152 | 0.0018 | 7.78E-17 |
| rs3608585 | 15 | 70379236 | C | 0.0201 | 0.0036 | 1.52E-08 |
| rs1564347 | 15 | 73438848 | T | 0.0088 | 0.0015 | 2.39E-09 |
| rs7495033 | 15 | 75206225 | A | 0.0095 | 0.0015 | 5.92E-10 |
| rs2469226 | 15 | 77246990 | A | 0.0096 | 0.0017 | 1.28E-08 |
| rs1291246 | 15 | 77309459 | T | -0.0104 | 0.0016 | 1.17E-10 |
| rs6200730 | 15 | 77723195 | A | 0.0264 | 0.0047 | 1.80E-08 |
| rs5639134 | 15 | 78006899 | A | 0.0148 | 0.0016 | 7.49E-20 |
| rs7182216 | 15 | 84422506 | C | -0.0096 | 0.0017 | 1.10E-08 |
| rs4778058 | 15 | 93456069 | T | -0.0092 | 0.0014 | 4.92E-11 |
| rs1697527 | 15 | 96046556 | A | -0.0102 | 0.0017 | 8.14E-10 |
| rs4984541 | 15 | 96911139 | A | -0.0138 | 0.0017 | 3.40E-16 |
| rs8030487 | 15 | 97119595 | A | 0.0085 | 0.0015 | 2.52E-08 |

| | | | | | | |
|---|---|---|---|---|---|---|
| rs4984682 | 16 | 740404 | C | -0.0138 | 0.0017 | 1.82E-16 |
| rs3431627 | 16 | 1018697 | A | 0.0097 | 0.0017 | 3.86E-09 |
| rs9933256 | 16 | 1246748 | A | 0.0119 | 0.0014 | 6.06E-17 |
| rs8046072 | 16 | 3562075 | A | 0.0101 | 0.0018 | 9.04E-09 |
| rs2077235 | 16 | 3629227 | T | 0.0097 | 0.0017 | 5.20E-09 |
| rs1107696 | 16 | 5811367 | T | 0.0086 | 0.0016 | 4.29E-08 |
| rs1270918 | 16 | 7249472 | A | -0.0091 | 0.0015 | 1.12E-09 |
| rs4787028 | 16 | 7533021 | T | 0.0086 | 0.0015 | 8.08E-09 |
| rs9929993 | 16 | 7664875 | T | -0.0102 | 0.0015 | 3.18E-12 |
| rs1164444 | 16 | 7943552 | A | 0.0117 | 0.0019 | 1.61E-09 |
| rs8052523 | 16 | 9293246 | T | -0.0084 | 0.0014 | 2.73E-09 |
| rs1174687 | 16 | 10205467 | A | -0.031 | 0.005 | 4.57E-10 |
| rs1097784 | 16 | 10249416 | T | -0.0083 | 0.0014 | 3.05E-09 |
| rs3501681 | 16 | 10266115 | T | 0.0176 | 0.0023 | 3.56E-14 |
| rs387027 | 16 | 12085343 | A | -0.0083 | 0.0015 | 1.36E-08 |
| rs350281 | 16 | 12231739 | T | -0.0117 | 0.0015 | 1.41E-14 |
| rs3409877 | 16 | 12237812 | A | -0.016 | 0.002 | 2.95E-16 |
| rs1035578 | 16 | 12531365 | A | -0.0094 | 0.0014 | 2.57E-11 |
| rs1122109 | 16 | 14674204 | A | 0.0105 | 0.0017 | 2.66E-10 |
| rs9927842 | 16 | 15153717 | T | -0.013 | 0.002 | 6.22E-11 |
| rs9924031 | 16 | 19267144 | C | 0.0093 | 0.0015 | 1.81E-10 |
| rs2851246 | 16 | 24711806 | C | 0.0112 | 0.0014 | 9.77E-15 |
| rs39998 | 16 | 28133152 | A | 0.009 | 0.0015 | 5.02E-09 |
| rs4255791 | 16 | 28274804 | A | -0.0084 | 0.0015 | 2.51E-08 |
| rs4787457 | 16 | 28555400 | A | 0.0162 | 0.0015 | 1.23E-28 |
| rs4788115 | 16 | 28998111 | A | 0.0111 | 0.0019 | 2.27E-09 |
| rs9927137 | 16 | 30634300 | A | 0.0082 | 0.0014 | 5.11E-09 |
| rs1164056 | 16 | 48791392 | A | 0.0112 | 0.002 | 2.77E-08 |
| rs4785187 | 16 | 49766772 | A | -0.01 | 0.0017 | 3.12E-09 |
| rs2052285 | 16 | 51188432 | A | 0.0112 | 0.0014 | 7.25E-15 |
| rs3809634 | 16 | 53538157 | A | -0.0109 | 0.0015 | 6.82E-13 |
| rs2542673 | 16 | 54206715 | A | 0.0088 | 0.0015 | 6.30E-09 |
| rs1724875 | 16 | 61579618 | A | -0.0136 | 0.0017 | 1.28E-15 |
| rs1438128 | 16 | 61773849 | A | -0.0113 | 0.0019 | 1.03E-09 |
| rs9929556 | 16 | 62193640 | T | -0.0099 | 0.0014 | 2.31E-12 |
| rs3575474 | 16 | 63127870 | T | 0.0083 | 0.0014 | 4.77E-09 |
| rs818415 | 16 | 65448079 | T | -0.011 | 0.0018 | 8.19E-10 |
| rs9926649 | 16 | 66750040 | T | 0.0177 | 0.0031 | 1.58E-08 |
| rs255053 | 16 | 68020492 | A | -0.0111 | 0.0018 | 6.68E-10 |
| rs9888796 | 16 | 68297589 | T | 0.0113 | 0.0016 | 1.20E-12 |
| rs7461509 | 16 | 69122043 | A | 0.0145 | 0.0026 | 3.67E-08 |
| rs6175720 | 16 | 70358495 | A | 0.0359 | 0.0063 | 1.27E-08 |
| rs9927049 | 16 | 71860008 | A | 0.0099 | 0.0016 | 9.28E-10 |
| rs1760434 | 16 | 72210865 | A | 0.0127 | 0.0018 | 2.36E-12 |
| rs6205114 | 16 | 72497721 | A | -0.0134 | 0.0022 | 1.79E-09 |
| rs9929762 | 16 | 78169675 | A | 0.0096 | 0.0014 | 1.03E-11 |
| rs1164718 | 16 | 82648514 | A | 0.0079 | 0.0014 | 4.92E-08 |

| | | | | | | |
|---|---|---|---|---|---|---|
| rs6152721 | 16 | 83608599 | A | 0.011 | 0.0014 | 1.13E-14 |
| rs1050847 | 16 | 87443734 | T | 0.0092 | 0.0014 | 9.20E-11 |
| rs7441546 | 16 | 89990843 | T | 0.0176 | 0.0026 | 6.25E-12 |
| rs8066044 | 17 | 1367352 | A | 0.009 | 0.0016 | 1.17E-08 |
| rs2447097 | 17 | 2278064 | T | 0.0096 | 0.0014 | 1.10E-11 |
| rs7218235 | 17 | 2564267 | A | -0.0097 | 0.0017 | 2.44E-08 |
| rs1773287 | 17 | 7362359 | T | -0.0095 | 0.0017 | 4.06E-08 |
| rs4925109 | 17 | 17661802 | A | -0.0096 | 0.0015 | 1.91E-10 |
| rs854796 | 17 | 18070761 | A | 0.009 | 0.0015 | 4.66E-09 |
| rs1260228 | 17 | 19236954 | T | 0.016 | 0.0021 | 4.62E-14 |
| rs4925065 | 17 | 19914546 | T | -0.0078 | 0.0014 | 2.59E-08 |
| rs870589 | 17 | 31614454 | A | -0.0079 | 0.0014 | 2.24E-08 |
| rs1165797 | 17 | 32900837 | A | -0.0091 | 0.0017 | 4.57E-08 |
| rs1842713 | 17 | 33200385 | A | -0.0117 | 0.0017 | 9.67E-12 |
| rs9649 | 17 | 34050934 | T | 0.0108 | 0.0019 | 6.13E-09 |
| rs1260138 | 17 | 34904985 | A | -0.0082 | 0.0014 | 7.42E-09 |
| rs1245368 | 17 | 37770005 | T | 0.0102 | 0.0015 | 1.81E-11 |
| rs2314338 | 17 | 38344485 | T | 0.0093 | 0.0016 | 9.50E-09 |
| rs4793090 | 17 | 40686342 | A | 0.0084 | 0.0015 | 1.67E-08 |
| rs10208 | 17 | 42300278 | T | 0.0091 | 0.0015 | 2.07E-09 |
| rs2521602 | 17 | 42336424 | A | -0.0266 | 0.0049 | 4.17E-08 |
| rs1187142 | 17 | 42920929 | A | 0.0128 | 0.0017 | 2.52E-14 |
| rs1165252 | 17 | 43055579 | A | -0.0154 | 0.0023 | 2.29E-11 |
| rs6503409 | 17 | 43058742 | T | 0.0087 | 0.0015 | 4.11E-09 |
| rs721579 | 17 | 43370481 | T | -0.011 | 0.0016 | 5.96E-12 |
| rs5631990 | 17 | 43871982 | T | -0.0206 | 0.0017 | 5.81E-33 |
| rs4458044 | 17 | 43873727 | C | 0.0101 | 0.0016 | 5.71E-10 |
| rs1232572 | 17 | 47028472 | A | 0.0088 | 0.0014 | 4.10E-10 |
| rs1760925 | 17 | 50274170 | T | -0.0087 | 0.0014 | 7.90E-10 |
| rs1051500 | 17 | 50395539 | T | 0.0143 | 0.0019 | 8.07E-14 |
| rs1762237 | 17 | 50726846 | T | -0.0113 | 0.0018 | 4.52E-10 |
| rs1295219 | 17 | 52077386 | T | -0.0081 | 0.0014 | 6.52E-09 |
| rs1695855 | 17 | 55659233 | T | 0.0096 | 0.0017 | 2.07E-08 |
| rs181214 | 17 | 56179482 | T | -0.0104 | 0.0017 | 8.19E-10 |
| rs5679481 | 17 | 60001978 | A | 0.0122 | 0.0019 | 1.69E-10 |
| rs6814558 | 17 | 75864805 | T | -0.0127 | 0.0021 | 7.53E-10 |
| rs7223311 | 17 | 78940614 | C | -0.0079 | 0.0014 | 1.77E-08 |
| rs1165734 | 17 | 79355294 | A | 0.014 | 0.0019 | 2.32E-13 |
| rs7190 | 18 | 5889765 | A | -0.0095 | 0.0015 | 1.50E-10 |
| rs1296701 | 18 | 9645035 | T | 0.0114 | 0.0017 | 1.33E-11 |
| rs8097125 | 18 | 13005473 | T | 0.0086 | 0.0014 | 2.31E-09 |
| rs303752 | 18 | 21074255 | A | -0.011 | 0.0014 | 1.47E-14 |
| rs1046009 | 18 | 22624708 | A | -0.0124 | 0.0014 | 2.61E-18 |
| rs1296284 | 18 | 22654102 | C | -0.0123 | 0.0021 | 2.38E-09 |
| rs1118522 | 18 | 25596884 | T | 0.0156 | 0.0022 | 4.85E-13 |
| rs7226824 | 18 | 27676827 | T | -0.0078 | 0.0014 | 2.72E-08 |
| rs1295342 | 18 | 31192529 | A | -0.008 | 0.0014 | 1.89E-08 |

| | | | | | | | |
|---|---|---|---|---|---|---|---|
| rs4426420 | 18 | 35104386 | T | -0.014 | 0.0019 | 2.63E-13 | |
| rs1108201 | 18 | 35145122 | T | 0.0194 | 0.0015 | 2.06E-38 | |
| rs5617131 | 18 | 35272539 | T | -0.0137 | 0.002 | 7.09E-12 | |
| rs1963381 | 18 | 36125986 | A | 0.0099 | 0.0016 | 8.80E-10 | |
| rs7290252 | 18 | 36565421 | T | -0.0103 | 0.0016 | 9.85E-11 | |
| rs1399808 | 18 | 36648364 | T | 0.018 | 0.0029 | 5.78E-10 | |
| rs978807 | 18 | 36898459 | A | 0.0139 | 0.0018 | 9.14E-15 | |
| rs2276209 | 18 | 36942457 | A | 0.0085 | 0.0015 | 3.49E-08 | |
| rs2852349 | 18 | 37182907 | T | -0.0093 | 0.0014 | 3.78E-11 | |
| rs1295746 | 18 | 37412228 | A | 0.0146 | 0.0018 | 9.20E-17 | |
| rs977143 | 18 | 38028165 | A | 0.0085 | 0.0015 | 3.95E-09 | |
| rs3429858 | 18 | 39026695 | A | 0.0111 | 0.0019 | 3.67E-09 | |
| rs9916901 | 18 | 40238335 | T | 0.0092 | 0.0016 | 8.40E-09 | |
| rs6209051 | 18 | 42671636 | A | -0.009 | 0.0015 | 6.88E-10 | |
| rs1166367 | 18 | 42908726 | T | -0.0121 | 0.0022 | 3.42E-08 | |
| rs1338155 | 18 | 44513371 | A | -0.0087 | 0.0014 | 7.40E-10 | |
| rs6209294 | 18 | 48779673 | T | -0.0087 | 0.0014 | 4.86E-10 | |
| rs1671269 | 18 | 50109971 | T | -0.01 | 0.0016 | 7.40E-10 | |
| rs1415869 | 18 | 50167764 | A | -0.009 | 0.0015 | 3.91E-09 | Not available |
| rs2043187 | 18 | 50394405 | A | 0.0107 | 0.0015 | 5.94E-13 | |
| rs1741133 | 18 | 50807359 | A | 0.014 | 0.0014 | 5.10E-23 | |
| rs1187662 | 18 | 52737309 | T | 0.0143 | 0.0024 | 1.52E-09 | |
| rs2958182 | 18 | 53149021 | A | 0.0082 | 0.0015 | 3.46E-08 | |
| rs613872 | 18 | 53210302 | T | -0.0163 | 0.0019 | 1.95E-18 | |
| rs1901024 | 18 | 57048571 | T | -0.021 | 0.0037 | 2.19E-08 | |
| rs2220926 | 18 | 58943386 | T | -0.0095 | 0.0014 | 1.84E-11 | |
| rs6567288 | 18 | 60218334 | A | -0.0079 | 0.0014 | 2.81E-08 | |
| rs7294406 | 18 | 63516255 | T | -0.0098 | 0.0016 | 9.39E-10 | |
| rs2554835 | 18 | 74141190 | A | 0.0088 | 0.0014 | 8.95E-10 | |
| rs1108152 | 18 | 75902735 | T | 0.0126 | 0.0015 | 5.22E-16 | |
| rs1420147 | 18 | 77511268 | A | 0.0102 | 0.0018 | 1.74E-08 | |
| rs1166360 | 18 | 77578191 | A | -0.013 | 0.0016 | 9.27E-17 | |
| rs1297026 | 18 | 77627950 | A | -0.0085 | 0.0014 | 1.61E-09 | |
| rs6210319 | 18 | 77634276 | T | -0.0147 | 0.0025 | 2.69E-09 | |
| rs1041175 | 19 | 1857297 | A | 0.0116 | 0.002 | 2.88E-09 | |
| rs8103741 | 19 | 2325005 | A | -0.0108 | 0.0019 | 5.17E-09 | |
| rs312927 | 19 | 3274770 | A | -0.0147 | 0.0022 | 2.52E-11 | |
| rs7517713 | 19 | 4386911 | T | 0.0298 | 0.0036 | 1.01E-16 | |
| rs1788333 | 19 | 4954455 | A | -0.0108 | 0.0018 | 1.29E-09 | |
| rs2287838 | 19 | 9959014 | A | -0.0104 | 0.0014 | 1.27E-13 | |
| rs6210986 | 19 | 13011809 | A | 0.0136 | 0.002 | 2.85E-11 | |
| rs1180401 | 19 | 13109531 | A | -0.0248 | 0.0038 | 8.47E-11 | |
| rs1215124 | 19 | 13212025 | T | -0.0149 | 0.0023 | 3.81E-11 | |
| rs1298140 | 19 | 19651577 | T | -0.0112 | 0.0019 | 2.75E-09 | |
| rs7254263 | 19 | 30741257 | T | -0.0119 | 0.0016 | 1.84E-14 | |
| rs1176234 | 19 | 32204489 | A | -0.0118 | 0.002 | 4.30E-09 | |
| rs7255223 | 19 | 32824310 | A | 0.0097 | 0.0016 | 1.06E-09 | |

| | | | | | | |
|---|---|---|---|---|---|---|
| rs3859523 | 19 | 32972296 | T | 0.0124 | 0.0018 | 9.92E-12 |
| rs173003 | 19 | 36240960 | A | -0.0082 | 0.0014 | 5.50E-09 |
| rs1040274 | 19 | 45815248 | T | 0.0077 | 0.0014 | 4.79E-08 |
| rs7624610 | 19 | 50121274 | A | -0.0162 | 0.0026 | 6.52E-10 |
| rs1924366 | 19 | 54960747 | T | -0.0319 | 0.0045 | 2.06E-12 |
| rs1408430 | 20 | 14917048 | C | 0.0096 | 0.0016 | 7.09E-10 |
| rs6043521 | 20 | 15739745 | T | -0.0083 | 0.0014 | 9.72E-09 |
| rs3612053 | 20 | 17438121 | T | 0.01 | 0.0018 | 3.77E-08 |
| rs175325 | 20 | 22317310 | A | -0.01 | 0.0014 | 2.24E-12 |
| rs6060308 | 20 | 33794378 | A | 0.0095 | 0.0016 | 1.76E-09 |
| rs1774754 | 20 | 41211913 | A | -0.009 | 0.0014 | 1.74E-10 |
| rs1006749 | 20 | 41716653 | A | 0.0079 | 0.0014 | 2.43E-08 |
| rs4369924 | 20 | 41954383 | A | 0.0125 | 0.0019 | 5.58E-11 |
| rs4812697 | 20 | 41975395 | T | 0.0171 | 0.0027 | 1.88E-10 |
| rs7310613 | 20 | 43491477 | T | -0.0147 | 0.0026 | 1.05E-08 |
| rs6065784 | 20 | 43717435 | C | 0.011 | 0.0015 | 5.78E-13 |
| rs3848715 | 20 | 44564682 | A | -0.008 | 0.0014 | 1.07E-08 |
| rs2024568 | 20 | 44732089 | T | -0.0097 | 0.0016 | 1.61E-09 |
| rs4810894 | 20 | 47532999 | A | -0.0085 | 0.0015 | 4.38E-09 |
| rs6020560 | 20 | 49119419 | T | -0.0083 | 0.0014 | 3.84E-09 |
| rs6091570 | 20 | 51299884 | A | 0.0081 | 0.0015 | 3.20E-08 |
| rs2252098 | 20 | 51620301 | T | -0.0077 | 0.0014 | 4.74E-08 |
| rs911149 | 20 | 58214637 | T | 0.0091 | 0.0017 | 3.93E-08 |
| rs6123924 | 20 | 58219764 | A | 0.0129 | 0.0019 | 3.20E-11 |
| rs6065080 | 20 | 59832791 | T | -0.013 | 0.0015 | 7.49E-19 |
| rs1305013 | 21 | 20018118 | A | -0.0081 | 0.0015 | 4.40E-08 |
| rs232496 | 21 | 22734409 | T | 0.009 | 0.0015 | 7.14E-10 |
| rs5930099 | 21 | 33072363 | T | -0.0164 | 0.0028 | 4.85E-09 |
| rs1544 | 21 | 33085585 | A | -0.0096 | 0.0016 | 1.76E-09 |
| rs2898191 | 21 | 34288509 | A | 0.0093 | 0.0015 | 1.88E-09 |
| rs6173971 | 21 | 34925689 | A | -0.0099 | 0.0016 | 3.74E-10 |
| rs743316 | 21 | 35252696 | T | 0.0112 | 0.0017 | 8.21E-11 |
| rs9974899 | 21 | 42006441 | T | -0.0093 | 0.0016 | 1.22E-08 |
| rs2838006 | 21 | 42653567 | T | 0.0132 | 0.0015 | 1.38E-19 |
| rs6648232 | 21 | 42688931 | C | 0.0124 | 0.0022 | 2.57E-08 |
| rs7958541 | 21 | 46487730 | T | -0.0178 | 0.0031 | 5.92E-09 |
| rs9977825 | 21 | 46494995 | T | -0.0092 | 0.0015 | 3.37E-10 |
| rs2297293 | 21 | 47125046 | C | 0.0099 | 0.0015 | 8.80E-11 |
| rs165633 | 22 | 29880773 | A | -0.0116 | 0.0016 | 1.27E-12 |
| rs737945 | 22 | 30202774 | C | -0.0115 | 0.0014 | 3.73E-16 |
| rs4128255 | 22 | 31286928 | A | 0.0237 | 0.004 | 2.09E-09 |
| rs5754581 | 22 | 33910736 | T | -0.0082 | 0.0014 | 5.85E-09 |
| rs7315454 | 22 | 34271689 | A | 0.0197 | 0.0036 | 4.57E-08 |
| rs5754753 | 22 | 34296751 | T | -0.0126 | 0.0016 | 6.15E-16 |
| rs5754762 | 22 | 34329224 | A | 0.0197 | 0.003 | 2.95E-11 |
| rs1170394 | 22 | 38817047 | A | -0.0168 | 0.0024 | 1.26E-12 |
| rs1437435 | 22 | 39289014 | A | 0.0118 | 0.002 | 4.99E-09 |

| | | | | | | |
|---|---|---|---|---|---|---|
| rs1217045 | 22 | 40019773 | A | 0.0105 | 0.0014 | 7.78E-14 |
| rs9611597 | 22 | 41864190 | A | -0.0107 | 0.0019 | 2.30E-08 |
| rs137079 | 22 | 42977907 | T | 0.0128 | 0.002 | 3.06E-10 |
| rs9616906 | 22 | 51104680 | A | 0.0122 | 0.0014 | 8.31E-18 |
| rs2873599 | 22 | 51135595 | A | 0.0162 | 0.0022 | 3.03E-13 |
| rs9616947 | 22 | 51151631 | T | -0.0134 | 0.0015 | 3.58E-20 |

| Okbay et al. 2016 | | | | | |
|---|---|---|---|---|---|
| Chromosome | Position hg19 (bp) | Effect Allele | Beta | P-value | Note |
| 1 | 8490603 | T | 0.019 | 1.79E-08 | |
| 1 | 43982527 | A | 0.017 | 2.36E-10 | |
| 1 | 72733610 | A | 0.035 | 3.76E-14 | |
| 1 | 72762169 | T | -0.017 | 1.80E-08 | |
| 1 | 91189731 | T | -0.016 | 6.01E-10 | |
| 1 | 2.05E+08 | A | 0.02 | 5.27E-10 | |
| 1 | 2.12E+08 | A | 0.015 | 1.55E-08 | |
| 1 | 2.44E+08 | A | 0.017 | 8.23E-09 | |
| 2 | 10977585 | T | 0.02 | 3.63E-08 | |
| 2 | 15621917 | C | 0.016 | 1.28E-08 | |
| 2 | 51873599 | A | 0.022 | 2.80E-08 | |
| 2 | 57387094 | A | 0.015 | 1.99E-08 | |
| 2 | 60757419 | T | -0.017 | 7.11E-10 | |
| 2 | 60976384 | A | -0.02 | 2.41E-08 | |
| 2 | 61482261 | A | -0.018 | 5.62E-10 | |
| 2 | 1E+08 | T | -0.018 | 1.70E-11 | |
| 2 | 1.01E+08 | T | -0.017 | 1.91E-08 | |
| 2 | 1.01E+08 | A | 0.027 | 2.69E-24 | |
| 2 | 1.44E+08 | T | -0.016 | 2.77E-09 | |
| 2 | 1.62E+08 | T | 0.016 | 2.65E-09 | |
| 2 | 1.63E+08 | T | -0.016 | 3.75E-10 | |
| 2 | 1.94E+08 | T | -0.015 | 4.70E-08 | |
| 2 | 1.94E+08 | A | -0.016 | 4.54E-09 | |
| 2 | 2.37E+08 | A | -0.022 | 5.37E-10 | |
| 3 | 48623124 | A | 0.034 | 3.82E-08 | |
| 3 | 48939052 | A | 0.048 | 2.63E-09 | |
| 3 | 49406708 | A | 0.025 | 1.36E-18 | |
| 3 | 49914397 | T | 0.024 | 3.30E-19 | |
| 3 | 50075494 | A | 0.036 | 4.61E-08 | |
| 3 | 85674790 | A | -0.016 | 7.01E-09 | |
| 3 | 1.61E+08 | C | -0.015 | 2.82E-08 | |
| 4 | 3249828 | T | 0.016 | 4.00E-08 | |
| 4 | 18037231 | T | -0.016 | 2.01E-08 | |
| 4 | 28801221 | T | 0.024 | 3.91E-08 | |
| 4 | 42649935 | T | -0.015 | 1.43E-08 | |
| 4 | 1.41E+08 | T | 0.018 | 1.56E-10 | |
| 5 | 45188024 | C | 0.019 | 3.32E-08 | |
| 5 | 57535206 | T | 0.015 | 3.02E-08 | |
| 5 | 60111579 | A | -0.017 | 3.49E-10 | |
| 5 | 87896602 | T | -0.015 | 1.91E-08 | |
| 5 | 87934707 | A | 0.021 | 2.46E-09 | |
| 5 | 1.04E+08 | T | 0.016 | 5.27E-09 | |
| 5 | 1.14E+08 | T | 0.017 | 3.42E-08 | |

| | | | | | |
|---|---|---|---|---|---|
| 5 | 1.2E+08 | T | 0.016 | 3.30E-08 | |
| 6 | 98187291 | T | -0.017 | 2.07E-09 | |
| 6 | 98584733 | A | 0.024 | 2.46E-19 | |
| 6 | 1.53E+08 | T | 0.017 | 2.44E-08 | |
| 7 | 23402104 | A | -0.037 | 4.71E-08 | |
| 7 | 39090698 | A | 0.014 | 3.11E-08 | |
| 7 | 92654365 | A | 0.016 | 9.15E-10 | |
| 7 | 1.28E+08 | A | 0.017 | 1.97E-08 | |
| 7 | 1.33E+08 | A | 0.02 | 1.14E-09 | |
| 7 | 1.35E+08 | A | -0.017 | 9.90E-10 | |
| 8 | 1.46E+08 | A | -0.016 | 3.93E-09 | |
| 9 | 1746016 | T | -0.016 | 4.35E-10 | |
| 9 | 23358875 | A | -0.023 | 2.20E-17 | |
| 9 | 88003668 | A | 0.015 | 2.25E-08 | |
| 9 | 1.25E+08 | T | -0.015 | 1.29E-08 | |
| 10 | 1.04E+08 | A | 0.018 | 5.44E-11 | |
| 10 | 1.04E+08 | T | -0.015 | 1.56E-08 | |
| 11 | 12748819 | A | 0.015 | 1.54E-08 | |
| 12 | 14653667 | A | 0.017 | 4.49E-10 | |
| 12 | 56416928 | A | -0.02 | 1.06E-12 | |
| 12 | 92159557 | A | 0.015 | 9.02E-09 | |
| 12 | 1.21E+08 | A | 0.014 | 3.46E-08 | |
| 12 | 1.24E+08 | A | 0.023 | 1.26E-12 | |
| 13 | 58402771 | A | 0.024 | 1.50E-16 | |
| 14 | 23373986 | A | 0.018 | 1.82E-11 | |
| 14 | 27098611 | A | -0.018 | 7.19E-09 | Not available |
| 14 | 84913111 | A | -0.019 | 3.55E-10 | |
| 17 | 43991515 | T | 0.025 | 1.47E-12 | Not available |
| 18 | 35186122 | A | -0.016 | 7.24E-09 | |
| 21 | 42620520 | T | 0.015 | 3.80E-08 | |
| 22 | 29880773 | A | -0.018 | 2.86E-09 | |
