## Supplementary Table 3 for "Using adopted individuals to partition maternal genetic effects into prenatal and postnatal effects on offspring phenotypes"

**Table S3.** Comparison of power estimated from asymptotic calculations and simulations using varying sizes
offspring pairs, 300,000 singletons, 6000 adopted individuals, and 50 biological mother-adopted offspring

| | | | | | | | | **Power to detect pre-nat** |
|---|---|---|---|---|---|---|---|---|
| | **Asymptotic power** | | | | | | | |
| | **Varying $\beta_m$ with $\beta_p$ and $\beta_o$ = 0** | | | | | | | |
| **$\gamma_m$** | **0** | **0.05** | **0.1** | **0.3** | **0.5** | | **$\gamma_m$** | **0** |
| **0** | 0.050 | 0.050 | 0.050 | 0.050 | 0.050 | | **0** | 0.042 (0.030, 0.054) |
| **0.05** | 0.202 | 0.202 | 0.204 | 0.215 | 0.237 | | **0.05** | 0.193 (0.169, 0.217) |
| **0.1** | 0.614 | 0.618 | 0.623 | 0.661 | 0.726 | | **0.1** | 0.600 (0.570, 0.630) |
| **0.15** | 0.924 | 0.927 | 0.931 | 0.953 | 0.977 | | **0.15** | 0.914 (0.897, 0.931) |
| **0.2** | 0.995 | 0.996 | 0.996 | 0.999 | 1.000 | | **0.2** | 0.995 (0.991, 0.999) |
| **0.25** | 1.000 | 1.000 | 1.000 | 1.000 | 1.000 | | **0.25** | 1 (1, 1) |
| | **Varying $\beta_p$ with $\beta_m$ and $\beta_o$ = 0** | | | | | | | |
| **$\gamma_m$** | **0** | **0.05** | **0.1** | **0.3** | **0.5** | | **$\gamma_m$** | **0** |
| **0** | 0.050 | 0.050 | 0.050 | 0.050 | 0.050 | | **0** | 0.042 (0.030, 0.054) |
| **0.05** | 0.202 | 0.202 | 0.204 | 0.218 | 0.250 | | **0.05** | 0.193 (0.169, 0.217) |
| **0.1** | 0.614 | 0.614 | 0.617 | 0.652 | 0.726 | | **0.1** | 0.600 (0.570, 0.630) |
| **0.15** | 0.924 | 0.924 | 0.925 | 0.942 | 0.970 | | **0.15** | 0.914 (0.897, 0.931) |
| **0.2** | 0.995 | 0.995 | 0.995 | 0.997 | 0.999 | | **0.2** | 0.995 (0.991, 0.999) |
| **0.25** | 1.000 | 1.000 | 1.000 | 1.000 | 1.000 | | **0.25** | 1 (1, 1) |
| | **Varying $\beta_o$ with $\beta_m$ and $\beta_p$ = 0** | | | | | | | |
| **$\gamma_m$** | **0** | **0.05** | **0.1** | **0.3** | **0.5** | | **$\gamma_m$** | **0** |
| **0** | 0.050 | 0.050 | 0.050 | 0.050 | 0.050 | | **0** | 0.042 (0.030, 0.054) |
| **0.05** | 0.202 | 0.203 | 0.204 | 0.220 | 0.261 | | **0.05** | 0.193 (0.169, 0.217) |
| **0.1** | 0.614 | 0.617 | 0.623 | 0.670 | 0.769 | | **0.1** | 0.600 (0.570, 0.630) |
| **0.15** | 0.924 | 0.927 | 0.930 | 0.955 | 0.986 | | **0.15** | 0.914 (0.897, 0.931) |
| **0.2** | 0.995 | 0.996 | 0.996 | 0.999 | 1.000 | | **0.2** | 0.995 (0.991, 0.999) |
| **0.25** | 1.000 | 1.000 | 1.000 | 1.000 | 1.000 | | **0.25** | 1 (1, 1) |

; of prenatal maternal genetic effects ($\gamma_m$), paternal genetic effects ($\beta_p$) or offspring genetic effects ($\beta_o$) with sa
pairs). Covariance between parental genotypes was fixed at 0.

| al maternal effect ($\gamma_m$) | | | |
|---|---|---|---|
| Simulation power (95% confidence interval) | | | |
| Varying $\beta_m$ with $\beta_p$ and $\beta_o$ = 0 | | | |
| **0.05** | **0.1** | **0.3** | **0.5** |
| 0.048 (0.035, 0.061) | 0.050 (0.036, 0.064) | 0.053 (0.039, 0.067) | 0.055 (0.041, 0.069) |
| 0.194 (0.169, 0.219) | 0.196 (0.171, 0.221) | 0.217 (0.191, 0.243) | 0.249 (0.222, 0.276) |
| 0.596 (0.566, 0.626) | 0.599 (0.569, 0.629) | 0.644 (0.614, 0.674) | 0.728 (0.700, 0.756) |
| 0.917 (0.900, 0.934) | 0.921 (0.904, 0.938) | 0.945 (0.931, 0.959) | 0.979 (0.970, 0.988) |
| 0.998 (0.995, 1.000) | 0.998 (0.995, 1.000) | 0.998 (0.995, 1.000) | 0.999 (0.997, 1.000) |
| 1 (1, 1) | 1 (1, 1) | 1 (1, 1) | 1 (1, 1) |
| Varying $\beta_p$ with $\beta_m$ and $\beta_o$ = 0 | | | |
| **0.05** | **0.1** | **0.3** | **0.5** |
| 0.045 (0.032, 0.058) | 0.046 (0.033, 0.059) | 0.047 (0.034, 0.060) | 0.054 (0.040, 0.068) |
| 0.195 (0.170, 0.220) | 0.204 (0.179, 0.229) | 0.224 (0.198, 0.250) | 0.238 (0.212, 0.264) |
| 0.598 (0.568, 0.628) | 0.605 (0.575, 0.635) | 0.647 (0.617, 0.677) | 0.717 (0.689, 0.745) |
| 0.913 (0.896, 0.930) | 0.915 (0.898, 0.932) | 0.935 (0.920, 0.950) | 0.965 (0.954, 0.976) |
| 0.997 (0.994, 1.000) | 0.996 (0.992, 1.000) | 0.998 (0.995, 1.000) | 0.999 (0.997, 1.000) |
| 1 (1, 1) | 1 (1, 1) | 1 (1, 1) | 1 (1, 1) |
| Varying $\beta_o$ with $\beta_m$ and $\beta_p$ = 0 | | | |
| **0.05** | **0.1** | **0.3** | **0.5** |
| 0.043 (0.030, 0.056) | 0.043 (0.030, 0.056) | 0.043 (0.030, 0.056) | 0.043 (0.030, 0.056) |
| 0.193 (0.169, 0.217) | 0.196 (0.171, 0.221) | 0.212 (0.187, 0.237) | 0.245 (0.218, 0.272) |
| 0.605 (0.575, 0.635) | 0.612 (0.582, 0.642) | 0.649 (0.619, 0.679) | 0.748 (0.721, 0.775) |
| 0.917 (0.900, 0.934) | 0.918 (0.901, 0.935) | 0.946 (0.932, 0.960) | 0.986 (0.979, 0.993) |
| 0.997 (0.994, 1.000) | 0.998 (0.995, 1.000) | 0.998 (0.995, 1.000) | 1 (1, 1) |
| 1 (1, 1) | 1 (1, 1) | 1 (1, 1) | 1 (1, 1) |

ample sizes approximating the number of individuals in UK Biobank with educational attainment informatio

| | | | | | | | | Power to detect post-nat |
|---|---|---|---|---|---|---|---|---|
| Asymptotic power | | | | | | | | |
| | Varying $\gamma_m$ with $\beta_p$ and $\beta_o$ = 0 | | | | | | | |
| **$\beta_m$** | **0** | **0.05** | **0.1** | **0.3** | **0.5** | | **$\beta_m$** | **0** |
| **0** | 0.050 | 0.050 | 0.050 | 0.050 | 0.050 | | **0** | 0.058 (0.044, 0.072) |
| **0.05** | 0.306 | 0.306 | 0.307 | 0.320 | 0.352 | | **0.05** | 0.326 (0.297, 0.355) |
| **0.1** | 0.828 | 0.830 | 0.832 | 0.854 | 0.897 | | **0.1** | 0.835 (0.812, 0.858) |
| **0.15** | 0.992 | 0.992 | 0.993 | 0.995 | 0.999 | | **0.15** | 0.990 (0.984, 0.996) |
| **0.2** | 1.000 | 1.000 | 1.000 | 1.000 | 1.000 | | **0.2** | 1 (1, 1) |
| **0.25** | 1.000 | 1.000 | 1.000 | 1.000 | 1.000 | | **0.25** | 1 (1, 1) |
| | Varying $\beta_p$ with $\gamma_m$ and $\beta_o$ = 0 | | | | | | | |
| **$\beta_m$** | **0** | **0.05** | **0.1** | **0.3** | **0.5** | | **$\beta_m$** | **0** |
| **0** | 0.050 | 0.050 | 0.050 | 0.050 | 0.050 | | **0** | 0.058 (0.044, 0.072) |
| **0.05** | 0.306 | 0.306 | 0.308 | 0.327 | 0.365 | | **0.05** | 0.326 (0.297, 0.355) |
| **0.1** | 0.828 | 0.829 | 0.832 | 0.860 | 0.902 | | **0.1** | 0.835 (0.812, 0.858) |
| **0.15** | 0.992 | 0.992 | 0.993 | 0.996 | 0.998 | | **0.15** | 0.990 (0.984, 0.996) |
| **0.2** | 1.000 | 1.000 | 1.000 | 1.000 | 1.000 | | **0.2** | 1 (1, 1) |
| **0.25** | 1.000 | 1.000 | 1.000 | 1.000 | 1.000 | | **0.25** | 1 (1, 1) |
| | Varying $\beta_o$ with $\gamma_m$ and $\beta_p$ = 0 | | | | | | | |
| **$\beta_m$** | **0** | **0.05** | **0.1** | **0.3** | **0.5** | | **$\beta_m$** | **0** |
| **0** | 0.050 | 0.050 | 0.050 | 0.050 | 0.050 | | **0** | 0.058 (0.044, 0.072) |
| **0.05** | 0.306 | 0.307 | 0.309 | 0.333 | 0.393 | | **0.05** | 0.326 (0.297, 0.355) |
| **0.1** | 0.828 | 0.830 | 0.834 | 0.867 | 0.926 | | **0.1** | 0.835 (0.812, 0.858) |
| **0.15** | 0.992 | 0.992 | 0.993 | 0.996 | 0.999 | | **0.15** | 0.990 (0.984, 0.996) |
| **0.2** | 1.000 | 1.000 | 1.000 | 1.000 | 1.000 | | **0.2** | 1 (1, 1) |
| **0.25** | 1.000 | 1.000 | 1.000 | 1.000 | 1.000 | | **0.25** | 1 (1, 1) |

n (1000 biological parent-offspring trios, 4000 biological mother-offspring pairs, 1800 biological father-

| cal maternal effect ($\beta_m$) | | | |
| --- | --- | --- | --- |
| Simulation power (95% confidence interval) | | | |
| Varying $\gamma_m$ with $\beta_p$ and $\beta_o$ = 0 | | | |
| **0.05** | **0.1** | **0.3** | **0.5** |
| 0.057 (0.043, 0.071) | 0.063 (0.048, 0.078) | 0.056 (0.042, 0.070) | 0.048 (0.035, 0.061) |
| 0.324 (0.295, 0.353) | 0.325 (0.296, 0.354) | 0.334 (0.305, 0.363) | 0.360 (0.330, 0.390) |
| 0.835 (0.812, 0.858) | 0.839 (0.816, 0.862) | 0.863 (0.842, 0.884) | 0.899 (0.880, 0.918) |
| 0.990 (0.984, 0.996) | 0.991 (0.985, 0.997) | 0.994 (0.989, 0.999) | 0.998 (0.995, 1.000) |
| 1 (1, 1) | 1 (1, 1) | 1 (1, 1) | 1 (1, 1) |
| 1 (1, 1) | 1 (1, 1) | 1 (1, 1) | 1 (1, 1) |
| Varying $\beta_p$ with $\gamma_m$ and $\beta_o$ = 0 | | | |
| **0.05** | **0.1** | **0.3** | **0.5** |
| 0.056 (0.042, 0.070) | 0.057 (0.043, 0.071) | 0.057 (0.043, 0.071) | 0.063 (0.048, 0.078) |
| 0.318 (0.289, 0.347) | 0.318 (0.289, 0.347) | 0.329 (0.300, 0.358) | 0.367 (0.337, 0.397) |
| 0.835 (0.812, 0.858) | 0.836 (0.813, 0.859) | 0.858 (0.836, 0.880) | 0.898 (0.879, 0.917) |
| 0.990 (0.984, 0.996) | 0.991 (0.985, 0.997) | 0.993 (0.988, 0.998) | 0.997 (0.994, 1.000) |
| 1 (1, 1) | 1 (1, 1) | 1 (1, 1) | 1 (1, 1) |
| 1 (1, 1) | 1 (1, 1) | 1 (1, 1) | 1 (1, 1) |
| Varying $\beta_o$ with $\gamma_m$ and $\beta_p$ = 0 | | | |
| **0.05** | **0.1** | **0.3** | **0.5** |
| 0.058 (0.044, 0.072) | 0.058 (0.044, 0.072) | 0.058 (0.044, 0.072) | 0.058 (0.044, 0.072) |
| 0.327 (0.298, 0.356) | 0.327 (0.298, 0.356) | 0.346 (0.317, 0.375) | 0.395 (0.365, 0.425) |
| 0.838 (0.815, 0.861) | 0.842 (0.819, 0.865) | 0.874 (0.853, 0.895) | 0.925 (0.909, 0.941) |
| 0.990 (0.984, 0.996) | 0.990 (0.984, 0.996) | 0.993 (0.988, 0.998) | 1 (1, 1) |
| 1 (1, 1) | 1 (1, 1) | 1 (1, 1) | 1 (1, 1) |
| 1 (1, 1) | 1 (1, 1) | 1 (1, 1) | 1 (1, 1) |
