## Supplementary Table 4 for "Using adopted individuals to partition maternal genetic effects into prenatal and postnatal effects on offspring phenotypes"

**Table S4.** Comparison of power to detect pre-natal ($\gamma_m$) and post-natal ($\beta_m$) maternal genetic effects est
number of individuals in the UK Biobank with educational attainment data (1000 biological parent-offsp
offspring pairs). Path coefficients representing postnatal or prenatal maternal genetic effects, paternal g

**a. Power to detect pre-natal maternal effect ($\gamma_m$)**

| Family structure | Asymptotic power | | | | |
|---|---|---|---|---|---|
| | Sample size | | | | |
| | **0** | **2000** | **4000** | **6000** | **8000** |
| G1: Parent offspring trios | 0.499 | 0.725 | 0.817 | 0.864 | 0.891 |
| G2: Mother offspring pairs | 0.442 | 0.593 | 0.640 | 0.663 | 0.676 |
| G3: Father offspring pairs | 0.453 | 0.651 | 0.719 | 0.752 | 0.772 |
| G4: Singletons | 0.544 | 0.564 | 0.577 | 0.586 | 0.593 |
| G5: Adopted singletons | 0.105 | 0.457 | 0.581 | 0.640 | 0.674 |
| G6: Adoptive mother adopted child pairs | 0.640 | 0.957 | 0.989 | 0.996 | 0.998 |
| G7: Biological mother adopted child pairs | 0.607 | 0.995 | 1.000 | 1.000 | 1.000 |

**b. Power to detect post-natal maternal effect ($\beta_m$)**

| Family structure | Asymptotic power | | | | |
|---|---|---|---|---|---|
| | Sample size | | | | |
| | **0** | **2000** | **4000** | **6000** | **8000** |
| G1: Parent offspring trios | 0.751 | 0.889 | 0.925 | 0.941 | 0.949 |
| G2: Mother offspring pairs | 0.842 | 0.846 | 0.847 | 0.847 | 0.847 |
| G3: Father offspring pairs | 0.658 | 0.856 | 0.905 | 0.926 | 0.937 |
| G4: Singletons | 0.705 | 0.735 | 0.755 | 0.769 | 0.780 |
| G5: Adopted singletons | 0.118 | 0.565 | 0.759 | 0.847 | 0.892 |
| G6: Adoptive mother adopted child pairs | 0.847 | 1.000 | 1.000 | 1.000 | 1.000 |
| G7: Biological mother adopted child pairs | 0.820 | 0.999 | 1.000 | 1.000 | 1.000 |

imated from asymptotic calculations and simulations using varying sample size for each of t
ring trios, 4000 biological mother-offspring pairs, 1800 biological father-offspring pairs, 300
genetic effects ($\beta_p$) and offspring genetic effects ($\beta_o$) were fixed to 0.1. Covariance between

| | | Simulation power (95 | | |
|---|---|---|---|---|
| | | Sampl | | |
| 10000 | | 0 | 2000 | 4000 |
| 0.908 | | 0.508 (0.477, 0.539) | 0.734 (0.707, 0.761) | 0.791 (0.766, 0.816) |
| 0.685 | | 0.429 (0.398, 0.460) | 0.602 (0.572, 0.632) | 0.624 (0.594, 0.654) |
| 0.785 | | 0.466 (0.435, 0.497) | 0.664 (0.635, 0.693) | 0.732 (0.705, 0.759) |
| 0.599 | | 0.559 (0.528, 0.590) | 0.586 (0.555, 0.617) | 0.579 (0.548, 0.610) |
| 0.696 | | 0.144 (0.122, 0.166) | 0.450 (0.419, 0.481) | 0.604 (0.574, 0.634) |
| 0.999 | | 0.624 (0.594, 0.654) | 0.954 (0.941, 0.967) | 0.988 (0.981, 0.995) |
| 1.000 | | 0.599 (0.569, 0.629) | 0.998 (0.995, 1.000) | 1 (1, 1) |

| | | Simulation power (95 | | |
|---|---|---|---|---|
| | | Sampl | | |
| 10000 | | 0 | 2000 | 4000 |
| 0.954 | | 0.725 (0.697, 0.753) | 0.895 (0.876, 0.914) | 0.905 (0.887, 0.923) |
| 0.848 | | 0.842 (0.819, 0.865) | 0.849 (0.827, 0.871) | 0.849 (0.827, 0.871) |
| 0.944 | | 0.660 (0.631, 0.689) | 0.839 (0.816, 0.862) | 0.904 (0.886, 0.922) |
| 0.788 | | 0.699 (0.671, 0.727) | 0.731 (0.704, 0.758) | 0.758 (0.731, 0.785) |
| 0.919 | | 0.137 (0.116, 0.159) | 0.585 (0.554, 0.616) | 0.755 (0.728, 0.782) |
| 1.000 | | 0.849 (0.827, 0.871) | 1 (1, 1) | 1 (1, 1) |
| 1.000 | | 0.821 (0.797, 0.845) | 0.998 (0.995, 1.000) | 1 (1, 1) |

he 7 family structures and the remaining family structures approximating the ,000 singletons, 6000 adopted individuals, and 50 biological mother-adopted parental genotypes was fixed to 0.

| **% confidence interval)** | | |
|---|---|---|
| **le size** | | |
| **6000** | **8000** | **10000** |
| 0.844 (0.822, 0.866) | 0.883 (0.863, 0.903) | 0.907 (0.889, 0.925) |
| 0.656 (0.627, 0.685) | 0.672 (0.643, 0.701) | 0.680 (0.651, 0.709) |
| 0.764 (0.738, 0.790) | 0.772 (0.746, 0.798) | 0.783 (0.757, 0.809) |
| 0.595 (0.565, 0.625) | 0.575 (0.544, 0.606) | 0.590 (0.560, 0.620) |
| 0.624 (0.594, 0.654) | 0.676 (0.647, 0.705) | 0.718 (0.690, 0.746) |
| 0.998 (0.995, 1.000) | 0.998 (0.995, 1.000) | 0.999 (0.997, 1.000) |
| 1 (1, 1) | 1 (1, 1) | 1 (1, 1) |

| **% confidence interval)** | | |
|---|---|---|
| **le size** | | |
| **6000** | **8000** | **10000** |
| 0.938 (0.923, 0.953) | 0.947 (0.933, 0.961) | 0.953 (0.940, 0.966) |
| 0.842 (0.819, 0.865) | 0.853 (0.831, 0.875) | 0.860 (0.838, 0.882) |
| 0.925 (0.909, 0.941) | 0.938 (0.923, 0.953) | 0.954 (0.941, 0.967) |
| 0.769 (0.743, 0.795) | 0.794 (0.769, 0.819) | 0.784 (0.758, 0.810) |
| 0.849 (0.827, 0.871) | 0.894 (0.875, 0.913) | 0.905 (0.887, 0.923) |
| 1 (1, 1) | 1 (1, 1) | 1 (1, 1) |
| 1 (1, 1) | 1 (1, 1) | 1 (1, 1) |
